# Large scale PVA modelling of insects in cultivated grasslands: the role of dispersal in mitigating the effects of management schedules under climate change

**DOI:** 10.1101/2021.11.21.469124

**Authors:** Johannes A. Leins, Volker Grimm, Martin Drechsler

## Abstract

In many species, dispersal is decisive for survival in a changing climate. Simulation models for population dynamics under climate change thus need to account for this factor. Moreover, large numbers of species inhabiting agricultural landscapes are subject to disturbances induced by human land use. We included dispersal in the HiLEG model that we previously developed to study the interaction between climate change and agricultural land use in single populations. Here, the model was parameterized for the large marsh grasshopper (LMG) in cultivated grasslands of North Germany to analyze (1) the species development and dispersal success depending on severity of climate change in sub regions, (2) the additional effect of grassland cover on dispersal success, and (3) the role of dispersal in compensating for detrimental grassland mowing. Our model simulated population dynamics in 60-year periods (2020-2079) on a fine temporal (daily) and high spatial (250 × 250 m^2^) scale in 107 sub regions, altogether encompassing a range of different grassland cover, climate change projections and mowing schedules. We show that climate change alone would allow the LMG to thrive and expand, while grassland cover played a minor role. Some mowing schedules that were harmful to the LMG nevertheless allowed the species to moderately expand its range. Especially under minor climate change, in many sub regions dispersal allowed for mowing early in the year, which is economically beneficial for farmers. More severe climate change could facilitate LMG expansion to uninhabited regions, but would require suitable mowing schedules along the path.

These insights can be transferred to other species, given that the LMG is considered a representative of grassland communities. For more specific predictions on the dynamics of other species affected by climate change and land use, the publicly available HiLEG model can be easily adapted to the characteristics of their life cycle.

## 1 Introduction

The 2021 IPCC assessment report (Masson-Delmotte et al., 2021) confirms that climate change poses a great threat to global biodiversity. Distribution of species is expected to change (Van der Putten et al., 2010), potentially leading to increased extinction risk as ranges shrink or species must persist in new communities. Species distribution models (SDMs) are therefore widely used to predict future distributions based on climate, habitat, and occurrence data (Srivastava et al., 2019). However, in fragmented and agricultural landscapes, extinction risk is at the same time severely affected by land use practices (Oliver and Morecroft, 2014). Accounting for these effects with SDMs, which are correlative, is particularly difficult for insects that require a representation of their life cycle on a fine temporal scale. Here, the timing of anthropogenic processes such as management schedules relative to the species life stage can be critical for population viability (Leins et al. 2021).

Population Viability Analyses (PVA) using mechanistic models are an important complement to SDMs for estimating the risk of species loss in changing and disturbed environments (Naujokaitis-Lewis et al., 2013). PVA models describe a species’ viability as a function of its life cycle, environmental conditions such as forage supply, and anthropogenic influences such as mechanical disruption of habitats (Beissinger and McCullough, 2002; Coulson et al., 2001).

Incorporating a dispersal process into such model analysis is considered another important factor for predicting both population viability (Driscoll et al., 2014) and species distribution (Bateman et al., 2013). According to metapopulation theory (Hanski, 1999; Levins, 1969), dispersal between habitats in a fragmented landscape can prevent extinction. Moreover, it is critical to the interpretation of SDMs whether and how quickly species can actually reach regions that have been projected to be suitable (Bateman et al., 2013).

In this study, we explore the combined effect of climate change and disturbance through land use on a dispersing species by conducting a PVA of the large marsh grasshopper (*Stethophyma grossum, hereafter referred to as* LMG, Figure 1) in cultivated grasslands of North Germany. The LMG, a well-studied species inhabiting wet meadows and marshes, is considered an indicator for the quality of grassland communities, similar to other grasshopper species (Báldi and Kisbenedek, 1997; Keßler et al., 2012; Sörens, 1996). It is a slow, yet fairly good disperser due to its flight ability (Sörens, 1996) and believed to extend its range in response to climate change (Leins et al., 2021; Poniatowski et al., 2020; Trautner and Hermann, 2008). Anthropogenic disturbances, in particular the timing of mowing events, affect the species differently depending on the current stage of the LMG’s life cycle. The German federal state of Schleswig-Holstein (SH) serves as study region, for which we extracted a highly resolved map of its grasslands (72,969 plots of roughly 6.25 ha each) using the software DSS-Ecopay (Mewes et al., 2012; Sturm et al., 2018). Three climate change projections of increasing severity up to the year 2080 function as environmental conditions, while mechanical grassland mowing applies as anthropogenic disturbance.

**Figure 1:**
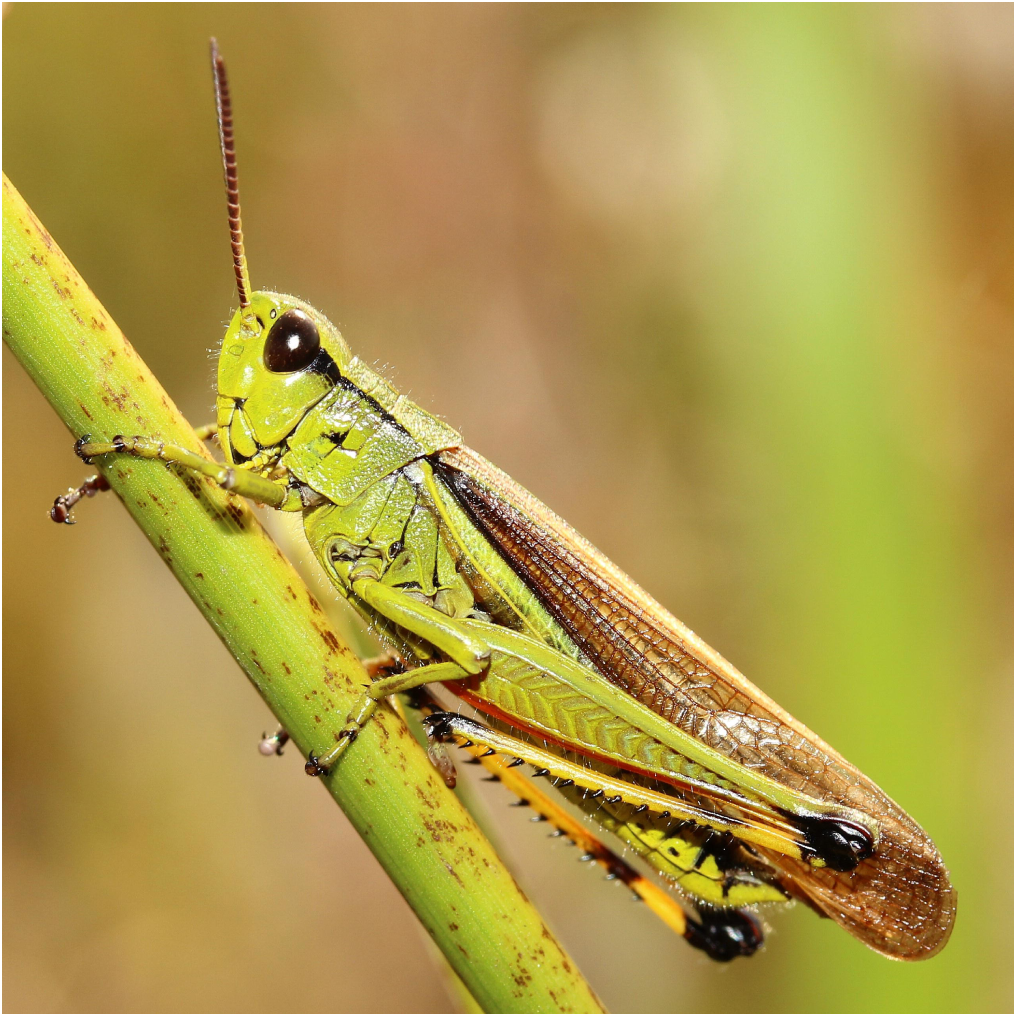
A male adult of the large marsh grasshopper, *Stethophyma grossum* (photo: Daniel Konn-Vetterlein)

We address the following three research questions regarding the LMG:

1. Are there (regional) differences in dispersal success depending on climate change scenario?
2. Is the success of dispersal additionally affected by spatial patterns such as grassland cover?
3. Can dispersal compensate for otherwise detrimental grassland mowing?

We used the PVA model HiLEG (*Hi*gh-resolution *L*arge *E*nvironmental *G*radient) introduced in Leins et al. (2021) and extended it by two features for the analysis of this study: the actual dispersal process within a predefined neighborhood of species habitats; and bilinear interpolation of the available, spatially coarse climate projections to achieve heterogeneous, gradual values of high spatial resolution throughout the study region. Originally, HiLEG is a spatially differentiated stage- and cohort-based simulation model that can be parameterized to represent the life cycle of terrestrial animal species, particularly insects. The new features together with the high-resolution grassland map render HiLEG from a spatially *differentiated* to a spatially *explicit* simulation model.

In Leins et al. (2021), we explored the effects of climate change on the LMG at a rather low spatial (12 × 12 km^2^) but high temporal resolution (daily time steps) while the 21 predefined management schedules were timed in one-week intervals between calendar week 20 and 40. We found that although the LMG mostly benefits from climate change, the timing of land use, i.e., the mowing schedule, is the most critical factor for the species’ survival. This is particularly relevant, because the intensification of anthropogenic grassland use in Germany is advancing (Bundesamt für Naturschutz, 2017). Moreover we showed that the high temporal resolution was required to detect the long-term impact of management schedules and climate change.

The original model, however, represented isolated local effects on the LMG and ignored dispersal between habitats. Due to the relevance of dispersal effects, we extended HiLEG by the features described above to analyze the implications of external drivers on dispersal success of the LMG. Using the realistic grassland map allows us to additionally consider the interaction between grassland distribution and dispersal at the landscape-scale.

The results of the simulations depend on the relative timing of dispersal and mowing events, but local effects of climate change and management may still dominate. Since high-resolution maps of species occurrence are often not available, a more general question is therefore, what the additive value of introducing higher spatial resolution and dispersal to a large-scale PVA model might be.

## 2 Material and methods

There are four main elements to our analysis: the study region (German federal state SH), the target species (LMG), climate data (projections from 2020 to 2080) and land use (grassland mowing). The following subsections include a description of these elements. Simulations for the present study are performed using an extension of the HiLEG model introduced in Leins et al. (2021). Section 2.5 includes a description of the model along with the relevant changes made to it for this study.

### 2.1 North German grasslands

Germany’s most northern federal state SH serves as study region. The state’s grassland areas, i.e., 72,969 cells of roughly 6.25 ha in size each, were extracted using the Software DSS-Ecopay (Mewes et al., 2012; Sturm et al., 2018) and mapped to 107 climate cells (Figure 2) of available climate projections (Section 2.3). Supplement S4 describes the mapping of all cells including their further specifications. Compared to the agricultural area of Germany as a whole (50.6 %), SH is the most intensively farmed federal state with 68.5 % (Statistisches Bundesamt, 2021).

**Figure 2:**
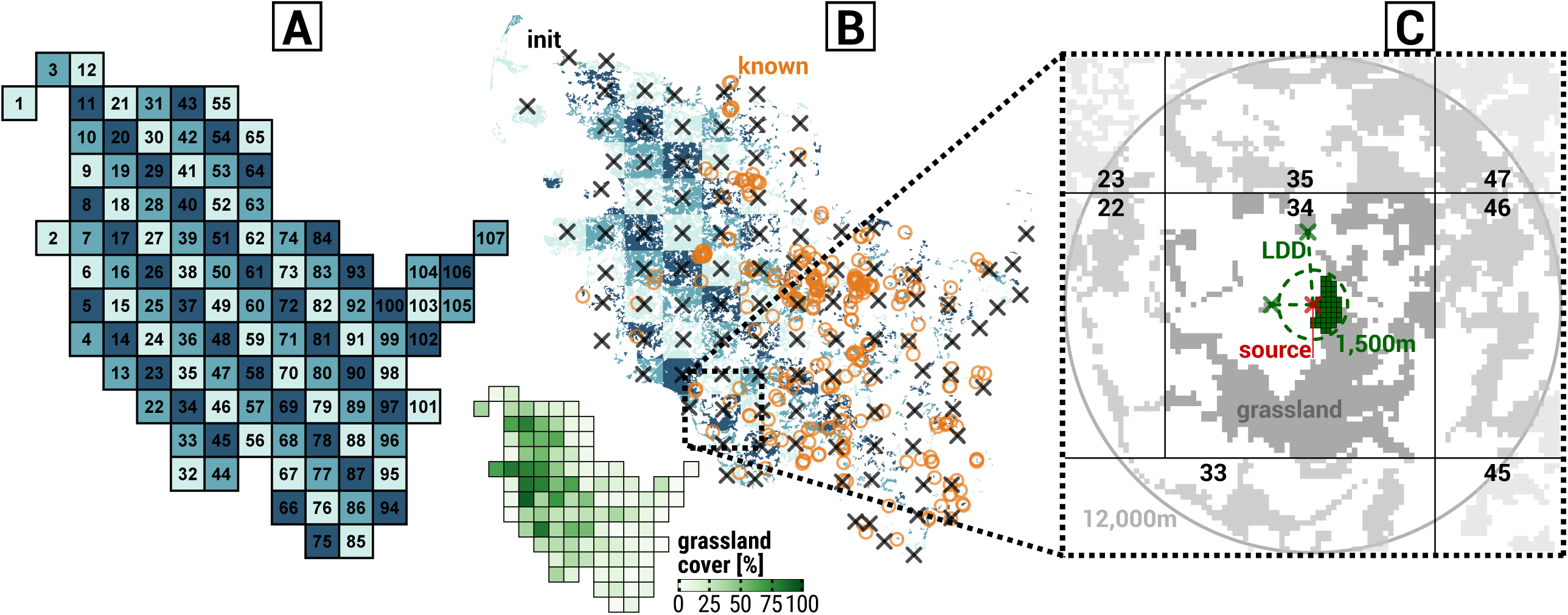
Distribution of 107 climate cells (A) and 72,929 grassland cells including grassland cover [%] per climate cell (B) in Schleswig-Holstein (SH), and grassland cells within a dispersal distance of > 12,000 m around the source habitat of a selected initial population (C). The numbers in A assign a unique ID to each climate cell. Turquoise colors in plots A and B are used to highlight mapping of grassland cells to respective climate cells. In plot B, black crosses mark the 107 initial populations that are closest to the respective climate cell’s geometric center and orange circles known LMG locations in SH in the years 2000 to 2016. The bottom subplot of B depicts the percental grassland cover within 1,500 m of the geometric per climate cell. In plot C, a red cross marks the source habitat of the initial population in climate cell 34. Green colors highlight grassland involved in dispersal: the dashed green circle around the source habitat represents the starting population’s dispersal radius of 1,500 m; green cells are available grassland within this radius; and the two green crosses connected to the source habitat by dashed lines represent the habitats reached via long distance dispersal (LDD). LDD applies, if there are no cells within the 1,500 m range of the source cell in either one of eight cardinal directions (North, Northeast, etc.) to reflect the assumption that nearby grassland, if present, is prioritized for colonization or as stepping stone for farther dispersal. Grey cells depict the remaining grassland that can be reached over time through dispersal from cells other than the source habitat, where the shades of grey from dark to light represent: grassland within the source climate cell 34; grassland outside the source cell but within a 12,000 m dispersal distance; and grassland in a dispersal distance farther than 12,000 m. The four vertical and horizontal black lines delimit the source climate cell from its seven neighbors identified by black numbers. Note: here, there is no neighboring climate cell to the Southwest

### 2.2 The large marsh grasshopper

The well-studied LMG (*Stethophyma grossum*, Linné 1758) is widely distributed in Central European grass- and wetlands (Heydenreich, 1999). Due to the high water requirements of its eggs, the species is bound to wet habitats such as meadows and marshes, although the grasshopper itself tolerates a wide range of temperatures and humidity (Ingrisch and Köhler, 1998; Koschuh, 2004). It used to be considered threatened in SH state (Winkler, 2000) but was recently given the status of “least concern” (Winkler and Haacks, 2019). Still, the LMG is regarded an indicator for the quality of grassland biotopes (Keßler et al., 2012; Sörens, 1996), similar to other grasshopper species (Báldi and Kisbenedek, 1997). The annual life cycle of the LMG (Figure 3) can be divided into the following five life stages, beginning with the stage after oviposition: (1) prediapause development inside egg, roughly occurring between July and November, below ground; (2) diapause (preventing too early development during mild winter months), November-March, below ground; (3) embryo development before egg hatching, March-June, below ground; (4) larva maturation, May-October, above ground; (5) imago (including oviposition), July-October, above ground; (Heydenreich, 1999; Ingrisch and Köhler, 1998; Kleukers et al., 1997; Köhler and Weipert, 1991; Malkus, 1997; Marshall and Haes, 1988; Oschmann, 1969). Although the majority of an LMG population usually stays within a close range of its hatching location (Malkus, 1997), it has been shown that new populations could establish in habitats several hundred meters from their origin within two years (Marzelli, 1994) while some offspring even reached distances of three or more kilometers (Keller, 2012; Van Strien, 2013). The latter is likely to be facilitated by the LMG’s flight ability (Sörens, 1996).

**Figure 3:**
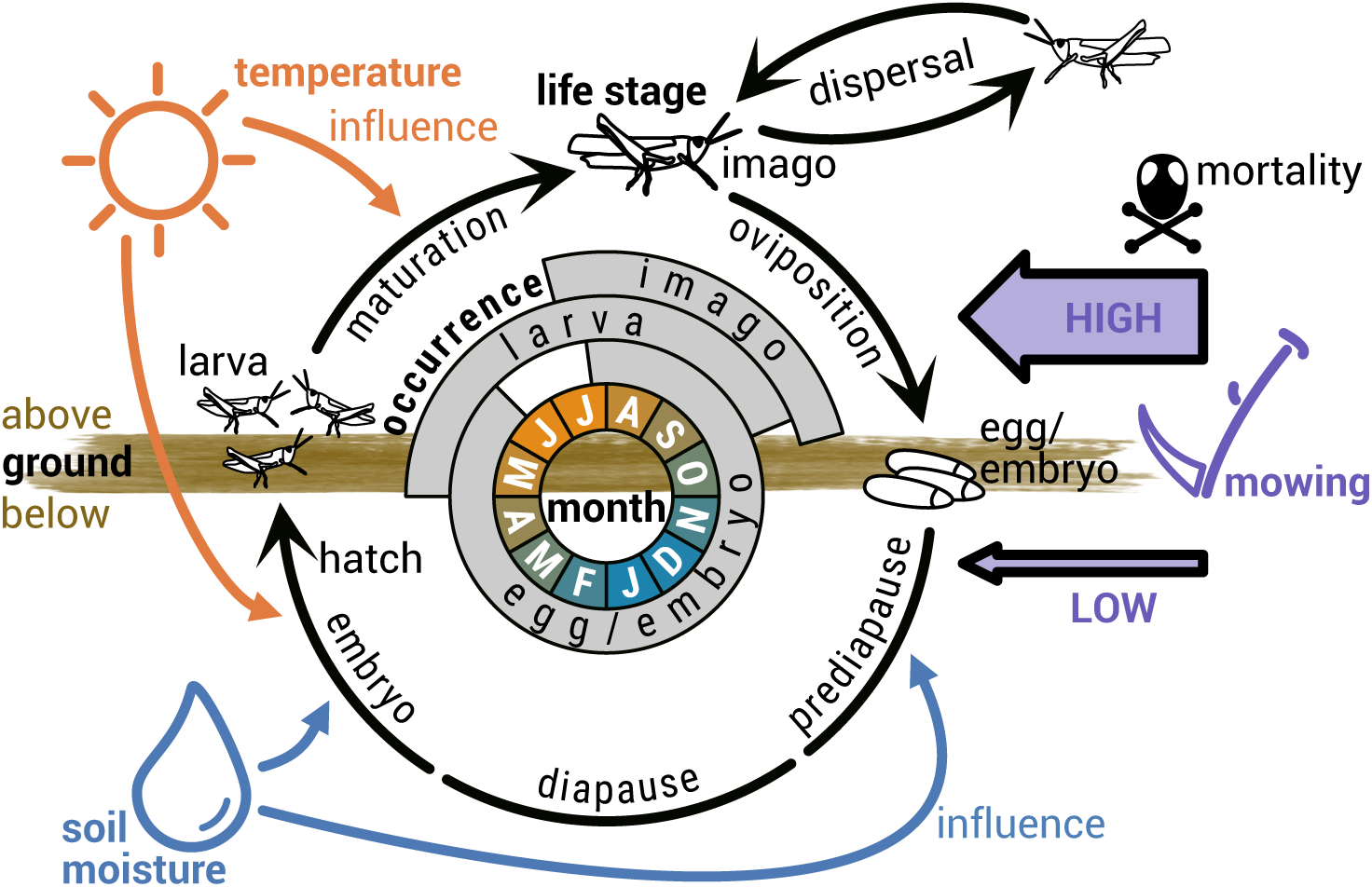
Yearly life cycle of the large marsh grasshopper, including the influence of external drivers. Black life stage symbols and arrows represent processes between and during life stages, where the life stage ‘egg/embryo’ is subdivided into three phases (broken arrow) and the dispersal process is directed to neighboring habitats. The typical ranges of the life stage occurrences are indicated in grey. The inner circle depicts months, where the color indicates seasonal changes in temperature. The influence of the external drivers of temperature, soil moisture and mowing is shown by colored symbols and arrows. Mowing impact is distinguished into high (aboveground) and low (belowground) mortality

Population development is affected differently by the climate conditions in an LMG habitat. Embryo hatching in spring (Wingerden et al., 1991) and larval development during summer (Ingrisch and Köhler, 1998; Uvarov, 1977) is accelerated by warm temperatures. Eggs/embryos experience stress in the event of a sustained dry soil before and after winter (Ingrisch, 1983). In the face of climate change, increasing temperatures might benefit the species by accelerating its development and expansion (Poniatowski et al., 2018; Trautner and Hermann, 2008) while extended droughts might prove detrimental for hygrophilous species like the LMG (Löffler et al., 2019).

### 2.3 High-resolution climate projections

We obtain climate data from high-resolution scenario simulations of the COSMO-CLM^1^ regional climate model (CCLM4-8-17) published by Keuler et al. (2016). In our analysis, the lateral boundaries of COSMO-CLM were controlled by simulation results from the global model ICHEC^2^-EC-EARTH and three Representative Concentration Pathways (RCPs) distinguished by action taken towards reducing CO_2_ emissions (in parenthesis): RCP2.6 (full force, Clim-FF), RCP4.5 (moderate, Clim-MOD) and RCP8.5 (business as usual, Clim-BAU). Time series of daily climate data (mean or sum) are provided by the regional model, spatially resolved to grid cells of size 12 × 12 km^2^. We used the years 2015-2080 of these time series and resampled them without losing long-term trends by randomly rearranging years within a 20-year time window (see Supplements S1, Section 5). This was necessary because the stochastic model processes (section 2.5) would otherwise have been limited by the fact that only a single, deterministic climate projection was available per global model, RCP and grid cell. Three climate parameters were relevant for the LMG population dynamics as implemented in our model: *surface temperature* [°C], *contact water* [kg m^-2^] and *relative humidity upper ground* [%].

We applied bilinear interpolation to the climate values of the four adjacent climate cells of each grassland cell to achieve heterogeneous, gradual climate data values of high spatial resolution throughout the grassland of the study region (Supplement S1, Section 7.6, Equations S1-25-27). This was done by weighing the distances from the center of the adjacent climate cells to a grassland cell of interest, multiplying their climate values by the resulting weights and summing up the results (Section 2.5). Figure 4 illustrates the calculation of the directional weights for a single grassland cell using a simplified geometric example. The calculated values of the weights per grassland cell are referenced in Supplement S4.

**Figure 4:**
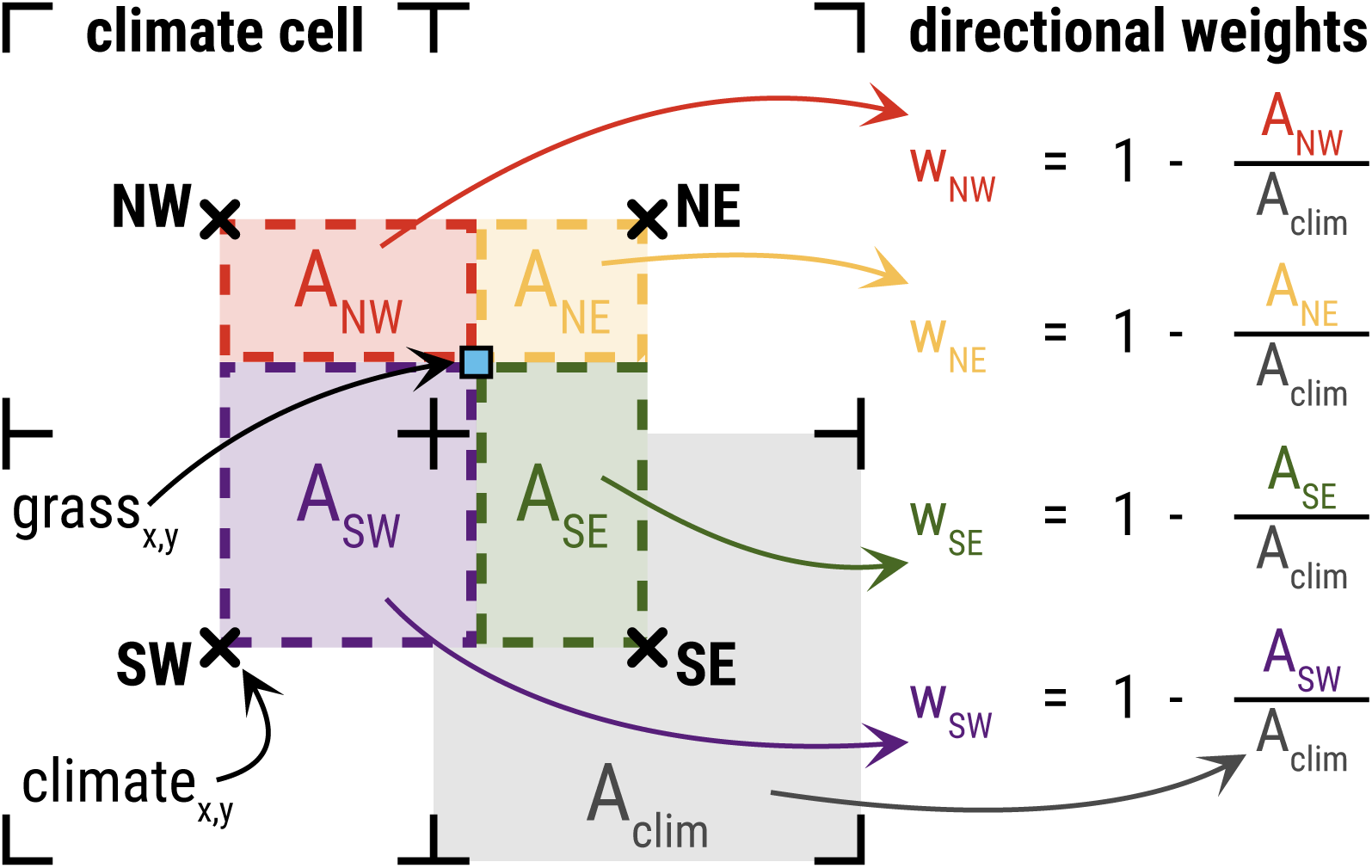
Calculation of the weights applied to bilinearly interpolate the climate values of four climate cells (black crosses) to achieve a distinct value for a grassland cell (blue square) that is enclosed by the climate cells. These directional weights are determined using the square area of a climate cell (A_clim_, grey) and the rectangular directional areas (A_NW_, red; A_NE_, yellow; A_SE_, green; A_SW_, purple) formed between the grassland cell coordinate and the center of the respective climate cell in secondary cardinal direction (Northwest, NW; Northeast, NE; Southeast, SE; Southwest, SW). Climate cells closer to the grassland cell result in smaller areas while receiving larger directional weights (w_NW_, w_NE_, w_SE_, w_SW_), which is accounted for by building the inverse of the ratio from directional area to climate cell area

### 2.4 Grassland mowing

Anthropogenic disturbances to the LMG are represented by mechanical grassland mowing that occurs two to three times per year depending on the mowing schedule (Table 1). In terms of our model, the impact of mowing on the model species is exclusively negative, though of different magnitude with respect to the above- and belowground life stages. Indirect effects of mowing, e.g. the observation that an early and/or late cut could maintain a beneficial vegetation structure for the species (Malkus, 1997; Miller and Gardiner, 2018; Sonneck et al., 2008), are not included in the model. However, such low-impact maintenance cuts with only a minor mortality effect on the LMG are accounted for by the base mowing schedule named *M20+00+44* (acronym: *M00*) that stands for an undisturbed environment and always takes effect where no other schedules apply. The first number of the schedule’s name stands for early mowing calendar week 20 (day 133) and the last number for late mowing week 44 (day 301). The middle number defines the (additional) mowing weeks 22-38 of more intensive grassland mowing schedules (acronyms: M22-M38). If either of the numbers in the schedule name is a double zero, the respective mowing time is omitted, so there are only two mowing events rather than three.

**Table 1:**
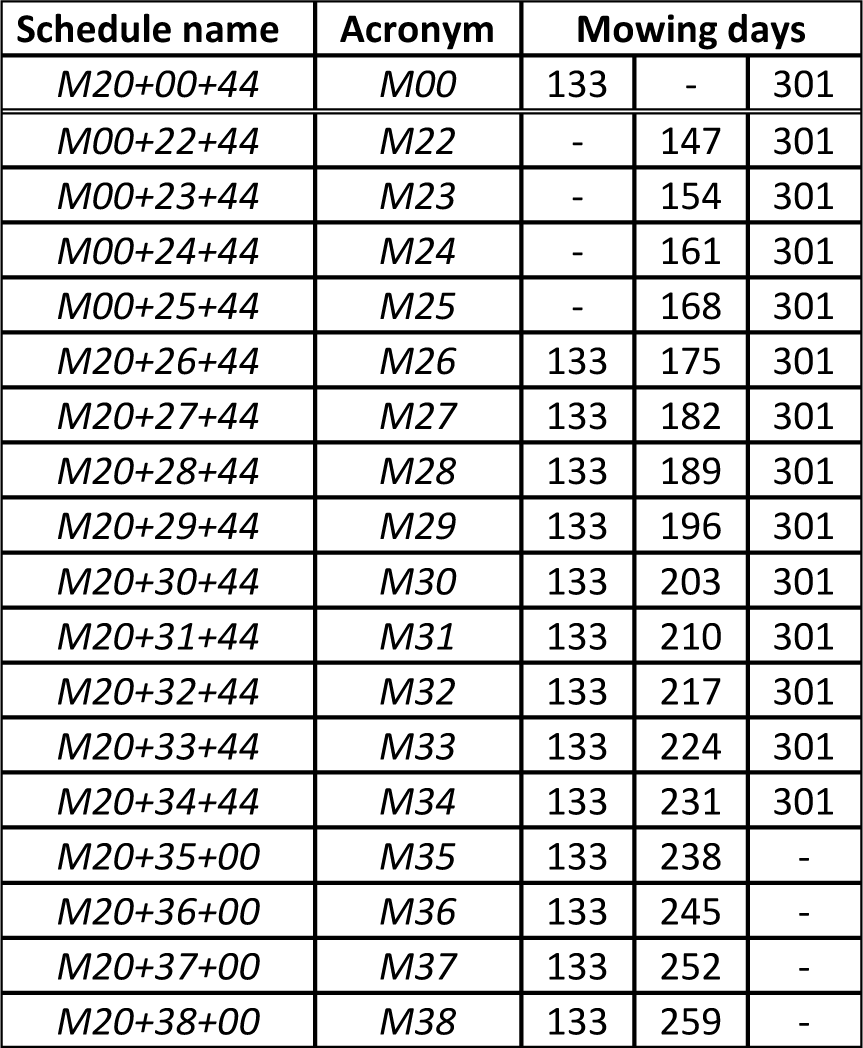
**Yearly grassland mowing schedules as applied in the simulation runs. First column gives the names of the 18 mowing schedules that encode the calendar weeks of yearly mowing occurrence divided by a plus (+) symbol. An acronym of the schedule name is provided in the second column, encoding the relevant mowing week in its name. The last three columns give the actual yearly mowing days (first day of respective calendar week) per mowing schedule. Schedules that include cells containing a dash (encoded by ‘00’ in the respective name) only have two mowing occurrences per year, all others have three. The first mowing schedule *M20+00+44* represents low-impact mowing, while more intensive mowing schedules follow in the rows below the double line**.

Gerling et al. (2022) gave the main lead for the rationale of these two-cut schedules: they are used for (1) the intensive schedules with mowing weeks 22-25 which omit early mowing in week 20, because grassland cuts are usually at least six weeks apart; and (2) the schedules of mowing weeks 35-38 which omit mowing in week 44, because an additional late maintenance cut is neither necessary, because of slowed grassland growth in autumn, nor economically beneficial for farmers.

### 2.5 Extended HiLEG model

A comprehensive description of the HiLEG model following the delta-ODD (Overview, Design concepts, Details) protocol (Grimm et al., 2020, 2006) is provided in Supplement S1. Here, we give a ‘Summary ODD’ (Grimm et al., 2020), which includes a digest of HiLEG’s main model description as introduced in Leins et al. (2021) and an overview of the extensions applied to the model for the present study. Unaffected mechanisms and parameters are either briefly described for general understanding or omitted in the main text. ODD keywords are in italics and capital letters hereafter.

We applied the HiLEG model to the LMG’s life cycle, with its life stages influenced by climate and land use. Both, the species’ development and mortality were affected by climate conditions, while the latter additionally increased during mowing events, especially in the species’ aboveground phase (Section 2.2). The model extension added a dispersal module rendering HiLEG spatially explicit, thus allowing dispersal between populations within a predefined radius. Essential climate variables were spatially differentiated on a large scale (12 × 12 km^2^) and resolved to a higher scale of 6.25 ha in the model extension through bilinear interpolation (Figure 4) to achieve relevant spatial gradients within the grasslands of the North German federal state SH (Figure 2B). We ignored other spatial heterogeneity in land use and biotic variables such as vegetation height or habitat size. Hereafter, all descriptions that neither concerned the dispersal process nor bilinear interpolation of the climate data were already included in the original model version.

While the ultimate *PURPOSE* of the HiLEG model is to analyze the regional effects of different climate change scenarios (CCS) and mowing schedules on the population viability of species such as the LMG (Leins et al., 2021), we here focuse on exploring the potential role of dispersal, which was ignored in the original version of HiLEG. The *PURPOSE* of the extended model is to answer the following questions:

- Are there (regional) differences in dispersal success depending on CCS?
- Is the success of dispersal additionally affected by spatial patterns such as grassland cover?
- Can dispersal compensate for otherwise detrimental grassland mowing?

We have drawn from literature the empirical *PATTERNS* that ensure the model is sufficiently realistic for its purpose, namely the observed characteristics of the species’ life cycle with its sensitivity to environmental conditions (Leins et al., 2021) and dispersal metrics (Griffioen, 1996; Malkus, 1997; Marzelli, 1994). The model’s design allowed for these empirical patterns to in principle emerge in the model as well (“pattern-oriented modelling”, Grimm and Railsback, 2012). In terms of population structure, density, persistence and dispersal, the model output was not compared to other data, since they are scarce. All model predictions are, thus, relative, not absolute. However, 251 known LMG habitats (Figure 2B, orange circles) adapted from survey data^3^ recorded in the years 2000 to 2016 were used to analyze some implications of regional effects. We used C++ for the implementation of the model’s source code. It is available for both, the original model and extensions, via a GitLab repository^4^ along with the executable program and the input files used for the simulation runs.

The following *ENTITIES* build the model’s core: *Climate Cells* (defining large scale climate conditions in a 12 × 12 km^2^ region), *Grassland Cells* (defining environmental conditions, e.g. interpolated climate values, on a scale of 250 × 250 m^2^), and per *Grassland Cell* a *Population* comprised of *Life Stages*, which are comprised of age-distinguished *Cohorts*. The most relevant *STATE VARIABLE* for the interpretation of the simulation results is the *density* [in individuals/eggs m^-2^] of a *Cohort, Life Stage* or *Population*. During a year, the LMG develops through five consecutive *Life Stages* (Figure 3): (1) prediapause, (2) diapause, (3) embryo, (4) larva, and (5) imago. *Density* is transferred to the next *Life Stage* (life cycle) or neighboring *Populations* (imago dispersal), and lost through mortality. The auxiliary *ENTITY Flow* controls the *density* transfer and in this function connects both the stages of the life cycle and habitats within a neighborhood. Environmental conditions such as climate, disturbances and grassland cover influence the amount of transferred/lost *density*. Daily hatching of eggs, maturation from larva to imago as well as dispersal rate and mortality are stochastically determined by drawing from density dependent binominal distributions. See Supplement S1, Sections 2, 4 and 7 for details on *ENTITIES, STATE VARIABLES* and for an overview of all stochastic elements of the model.

The model runs on basis of daily time steps where the *SCALE* corresponds to the sampling of the climate data. By definition, a year has 364 days (52 full calendar weeks) to account for the weekly mowing schedules. A simulation run takes 21,840 time steps (60 years) starting in the beginning of 2020 and terminating by the end of 2079. In the case of premature extinction of all *Populations*, simulations stop earlier. Each *Grassland Cell* in the study region (Section 2.1) represents a potential species habitat and is connected to cells within a predefined radius to allow dispersal between habitats.

To be able to better observe dispersal effects and explore the potential role of dispersal for population viability, we chose an artificial *INITIAL* setting for each simulation run in terms of species distribution: a single *Population* was placed at the center of one of the 107 *Climate Cells* (i.e., the *Grassland Cell* closest to the geometric center of the *Climate Cell*, cf. black crosses in Figure 2B), while all other *Grassland Cells* initially remained unoccupied. This initial setup was repeated separately for each of the 107 distinct Climate Cells. Furthermore, a simulation run was *INITIALIZED* with one out of three CCS (section 2.3) and one of 18 mowing schedules (Table 1). The artificial setup with a single starting location per simulation run, thus no initial populations at other locations of the study region, allowed us studying regional dispersal effects independent of potential immigration from other starting locations.

A similar simplification is inherent in the mowing schedules, which were comprised of fixed dates, whereas in reality farmers would to some degree respond to, e.g., an earlier beginning of vegetation growth due to climate change by shifting mowing to earlier dates. However, to account for this would require both a grassland model capable of predicting climate change response and a model of farmer decision making. This would have made our model considerably more complex and uncertain. Instead, we focused on the changing influence of a fixed mowing date on the climate-related shifting life cycle of the grasshopper. While dynamic schedules would likely change the quantitative model results and might shift potential thresholds in output parameters in time, our approach was sufficient for the objective of comparing the qualitative long-term effects on dispersal success between mowing schedules.

The *Population* at a starting location received an initial density per *Life Stage* (i.e., 0.725 eggs m^-2^ in the diapause stage, zero density for all other *Life Stages*). Independent of the defined mowing schedule, a starting location was only exposed to the low-impact mowing schedule M20+00+44 by default to serve as rather undisturbed source of dispersal to their close vicinity. *Populations* at non-starting locations were initialized empty and receive their density through potential immigration. All non-starting locations were subject to the initially defined mowing schedule.

For comparability with the original model setup of spatially stationary populations, we additionally ran the simulations with low-impact mowing schedule M20+00+44 while the dispersal process was disabled. Thereby, the starting population remained confined to its source habitat and was predominantly affected by climate. Comparison with the other mowing schedules was not practical because, as described above, they were not applied to the source habitats in the simulations with dispersal.

Distinct climate data time series were employed as *INPUT DATA* per *Climate Cell* to drive the model dynamics. To have heterogeneous, gradual values at the location of each *Grassland Cell* as well, bilinear interpolation was applied using the climate data of the (up to) four closest adjacent neighbors. This was achieved by weighing the distances from a *Grassland Cell G*_*a*_ to the center of the (up to) four *Climate Cells* {Ω_*a,NE*_, Ω_*a,SE*_, Ω_*a,SW*_, Ω_*a,NW*_} into secondary cardinal directions of *G*_*a*_. The resulting bilinear weights 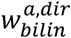 were multiplied with their respective climate values 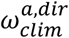 and then summed to achieve the interpolated value 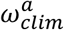 at *G*_*a*_.

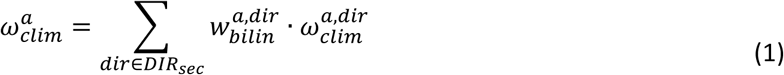

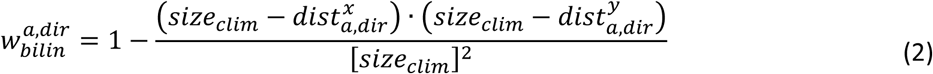

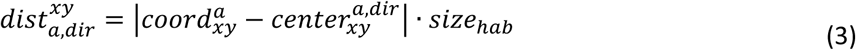

Here, *DIR*_*sec*_ ⊂ *DIR* = {*N, NE, E, SE, S, SW, W, NW*} are the secondary cardinal directions NE, SE, SW and NW of the cardinal directions North (N), Northeast (NE), East (E), Southeast (SE), South (S), Southwest (SW), West (W) and Northwest (NW) and 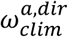 is the projected value in the *Climate Cell* Ω_*a,dir*_ into direction *dir* of *G*_*a*_. Size of *Climate Cell* and *Grassland Cell* are given by *size*_*clim*_ and *size*_*hab*_ (Table 2). The value 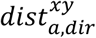 for the distances in x- and y-direction was calculated using the geometric center of a *Climate Cell* 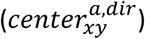.

**Table 2:**
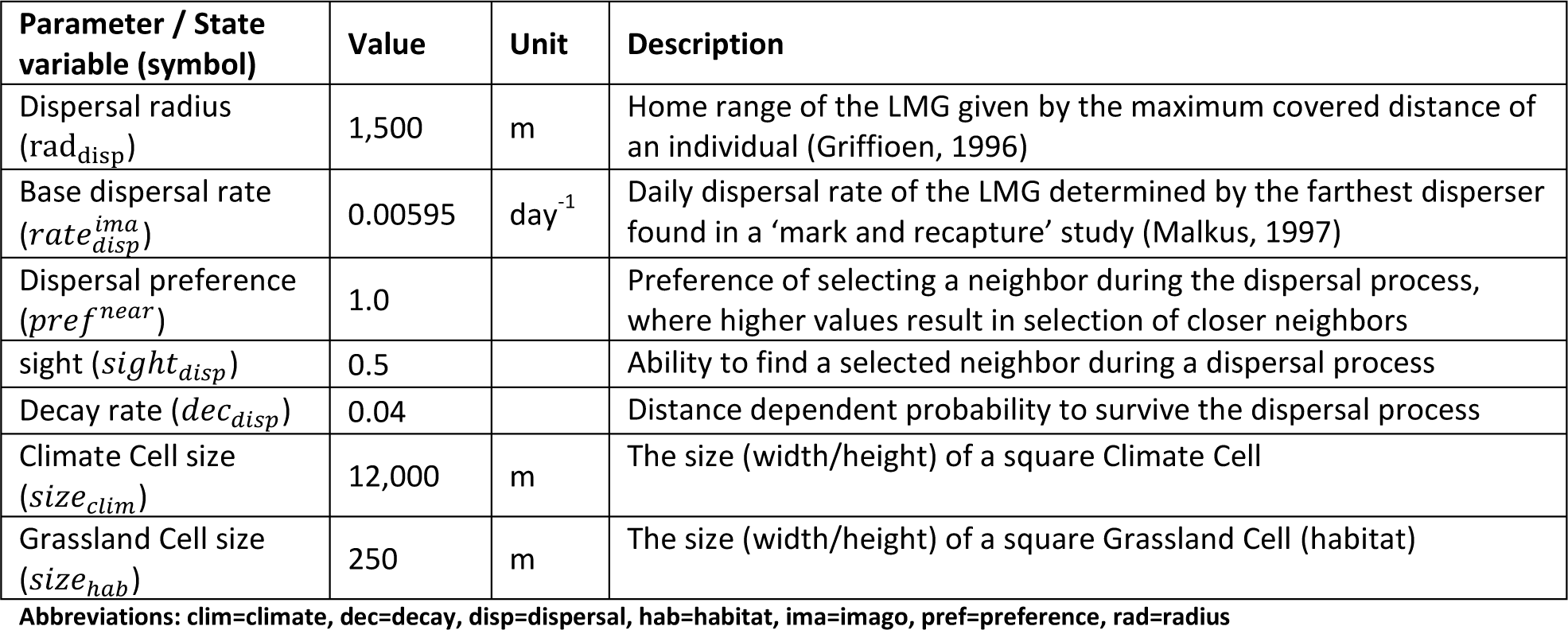
**Simulation parameters as used for the model extension and dispersal process of the large marsh grasshopper (LMG). The first column gives the parameter name and the respective symbol applied in equations. Second and third column contain the parameter’s value and unit (if any). The fourth column gives a more detailed description of the parameter**.

Six main *PROCESSES* are included in the model: ‘Update environmental drivers’, ‘Flow update’, ‘Life Stage update’, ‘Cohort update’, ‘Bilinear climate interpolation’ and ‘Dispersal setup’. Their scheduling is described in Pseudocode S1-1 of Supplement S1, Section 3. The first three of these processes are *SCHEDULED* for every inhabited Grassland *Cell* and during each time step of a simulation run. ‘Cohort update’ and ‘Bilinear climate interpolation’ are submodels of ‘Life Stage update’ and ‘Update environmental drivers’, respectively, and thus *SCHEDULED* every time step, as well. ‘Dispersal setup’ is only *SCHEDULED* in the event of an empty *Grassland Cell* becoming inhabited. Additionally, five types of sub *PROCESSES* can be associated with a *Life Stage* that apply depending on parametrization: (1) mortality (all stages), (2) development (prediapause, diapause), (3) transfer (all except imago), (4) reproduction (imago only), and (5) dispersal (imago only). For all sub processes, a daily *base rate* representing benign or observed average environmental conditions is assumed. Environmental drivers can modify (‘influence’) these base rates of the processes using predefined functions called *Influences* (Supplement S1, Section 7.1)

While the first four above sub-processes were already part of the original model, bilinear climate interpolation (see above) and the dispersal process are introduced in the present version. Figure 2C depicts the relevant grassland cells included for calculating the dispersal from an exemplary source population inside the climate cell with ID 34 to the neighborhood in reach as it was applied for the LMG. Cells belonging to this neighborhood are either within a predefined radius rad_disp_ (Table 2) from the source population in question; or the closest (if any) grassland cells in each of the eight cardinal directions *DIR* (see above) that have no neighbors within the predefined radius. The latter is called long distance dispersal (LDD, see Supplement S1, Sections 7.1.7 and 7.5) and is included to account for the LMG’s flight ability (Sörens, 1996) that is especially relevant in otherwise isolated habitats. Figure 5 illustrates how and in which cases grassland cells are selected for the LDD process.

**Figure 5:**
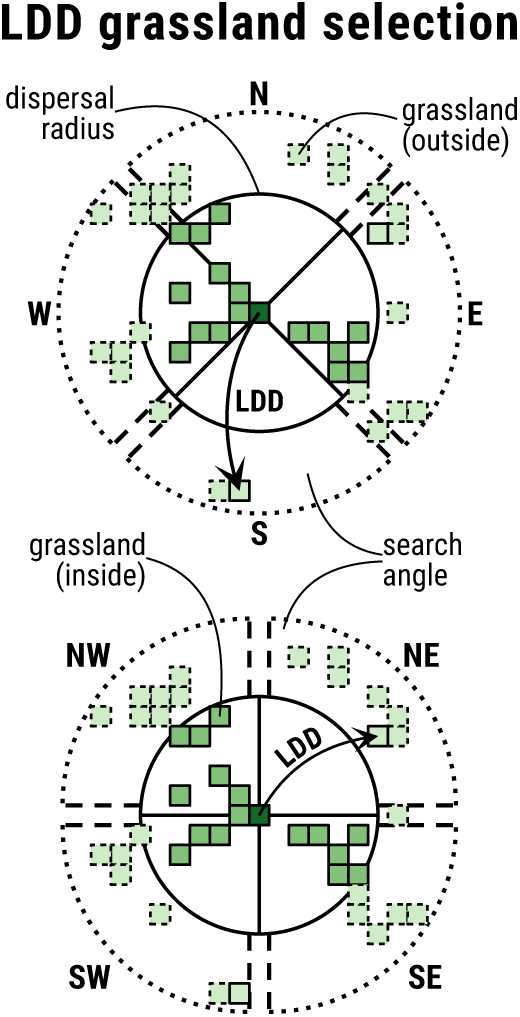
Determination of grassland cells for long distance dispersal (LDD) in cardinal (TOP) and secondary cardinal direction (BOTTOM) of a source habitat (dark green). The solid black circle encompasses grassland cells (medium green) within a defined dispersal radius. Light green cells represent grassland outside of the dispersal radius, where dashed cells are unreachable from the source habitat and solid cells are selected for LDD. Each two of the dashed black lines in the outer ring of the plot, which are approximately perpendicular to each other, enclose the angle at which cells for LDD are searched in that direction. Longer distances than indicated here are possible. Cells for LDD are only searched in case no grassland is found within the dispersal radius of either one of the directions to reflect the assumption that nearby grassland, if present, is prioritized for colonization or as stepping stone for farther dispersal Cells in straight secondary cardinal or cardinal direction (spaces between parallel dashed lines in the outer ring) are ignored for the search in cardinal or straight cardinal direction, respectively.

The dispersal rate (Eq. 4) between any population *P*_*a*_ and a neighbor *P*_*b*_ within the neighborhood N_*a*_ is stochastically determined each time step using the *base dispersal rate* defined for the *Life Stage* of interest (here, 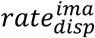 for the LMG’s imago stage) and a dispersal probability 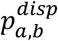 (Eq. 5):

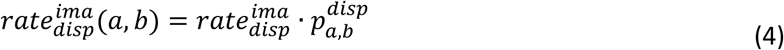

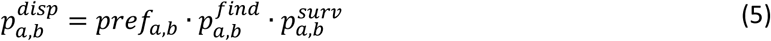

The dispersal probability itself is calculated using a preference factor (*pref*_*a,b*_) to select nearby target populations depending on the distance to all neighbors, a probability 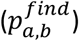 to find the selected neighbor during the dispersal process and a probability 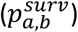 to survive the dispersal, where both latter probabilities depend on grassland cover. Parameters applied to adjust the three factors/probabilities are given in Table 2. Furthermore, the dispersers are subject to dispersal mortality which is the difference of the sum of all the dispersal probabilities multiplied by the *base dispersal rate*:

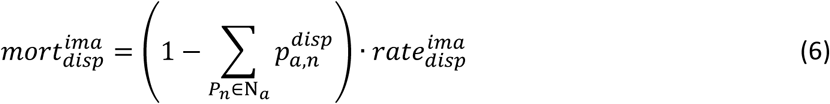

Table 2 gives an overview of the simulation parameters additionally defined model extension. We followed the maximum covered distance of 1,500 m described by Griffioen (1996) for the LMG’s dispersal radius and defined the individuals that travelled the largest distance during one day (1 out of 168) in a ‘mark and recapture’ study by Malkus (1997) as dispersers to determine the daily base dispersal rate. The remaining dispersal parameters were approximated using initial test simulations and their usage is explained in more detail in Supplement S1, Section 7.5.

The model output was *DESIGNED* in such a way that different aspects of a population development and dispersal success could be *OBSERVED* or rather analyzed with respect to the study’s *PURPOSE*. This data is distinguished into direct parameters produced for each inhabited cell during the simulation runs of the model itself, and indirect parameters calculated for a region’s whole population in the post-processing of the direct output. Relevant evaluation parameters of the first category are the daily *life stage densities* given in individuals/eggs m^-2^ depending on the respective stage. They allow *OBSERVING* the actual dispersal process over time. The indirect evaluation parameters, used to facilitate the *OBSERVATION* of a population’s dispersal success, are the following four: (1) the *dispersal distance* in meters from a source habitat to an occupied habitat, (2) the *established distance* in meters from a source habitat to a habitat with imago density ≥ 0.002 individuals m^-2^ during a year, (3) the *population size* in total number of individuals/eggs in all established habitats, and (4) the *population density* in individuals/eggs m^-2^ for all established habitats. For the analysis, it was convenient to consider parameters (1) and (2) on the basis of their maximum value, i.e., the habitat farthest from the source, to compare dispersal success. All parameters were determined only using inhabited cells at the end of a simulation year to match values in the same life stage (typically diapause) and after mowing schedules had been fulfilled. In the following, *population size* and *density* are thus given in *eggs*, because populations are usually in the diapause life stage by the end of the year.

## 3 Results

We added a representation of dispersal to the model of Leins et al. (2021). To make sure to achieve a realistic representation, despite sparse quantitative data on dispersal of LMG, we compared dispersal distances in the model to the findings of Marzelli (1994). The author found that in a natural and anthropogenically undisturbed grassland environment new LMG populations could establish in a distance of 400 m from an existing population within two years. Considering the defined measure for established distance (Section 2.5) our model confirmed these findings: within an environment of moderate grassland mowing (Table 1, M20+00+44) the LMG on average established in distances of about 14,000 m in 60 years (i.e., roughly 467 m every two years).

With this model version, we obtained the following key results: (1) qualitative patterns of dispersal success are similar in the study region independent of CCS, but there are regions that allow more successful dispersal depending on severity of climate change; (2) spatial patterns have an effect on foremost population size (high grassland cover) and dispersal distance (low grassland cover); and (3) mowing schedules that might seem problematic when looking at an isolated habitat could still allow (slowed) dispersal outside of a population’s home range. These results are described in more detailed in the following.

The dispersal success of the LMG differed depending on region, climate change scenario (CCS) and mowing schedule. Figure 6 depicts different outcomes of the dispersal process exemplary for an initial population in the center of climate cell 34 (black arrow in top left subplot). The first two rows of Figure 6 show the distribution of LMG populations chronologically in 15-year-steps in an undisturbed (first row) and disturbed environment (second row) using the Clim-MOD scenario: Under ideal conditions with low-impact mowing (Table 1, M20+00+44), the LMG continuously spread out until it occupied grasslands in a distance > 20,500 meters and established in grasslands in a distance > 12,500 meters in the year 2079 (Figure 6, first row). The dispersal process became significantly slowed down (10,250 / 8,000 m) when a mowing schedule with a deviating early cut in calendar week 23 (M00+23+44) was applied (Figure 6, second row). The three bottom columns of Figure 6 compare the final dispersal success in the year 2079 between simulation scenarios. It became evident that the mowing schedule had a different impact on the dispersal success depending on which CCS occurred: Deviating early mowing in, for instance, week 23 (Figure 6, third row, M00+23+44) still allowed substantial dispersal success for the LMG in the event of the less severe Clim-FF scenario (Figure 6, first bottom column), while it already became quite inhibited for both other scenarios (Figure 6, second/third bottom column), especially Clim-MOD. Mowing just one week later (Figure 6, M00+24+44) already had a great negative impact on the dispersal success in all three CCS, allowing population establishment only in close vicinity of < 5,000 m for the Clim-FF scenario while restricting it to grassland roughly within the dispersal radius of 1,500 m from the source habitat for Clim-MOD and Clim-BAU. The strong negative impact in climate cell 34 continued for several weeks and the dispersal success afterwards became more inhibited for the Clim-FF scenario, shown on the example of additional mowing in calendar week 34 (Figure 6, M20+34+44). Later schedules starting with additional mowing in calendar week 35 (Figure 6, M20+35+00) allowed for gradual improvement in dispersal success. In these cases, the dispersal process became slightly more successful in the Clim-MOD than in the Clim-BAU scenario and remained restricted the most for the Clim-FF scenario.

**Figure 6:**
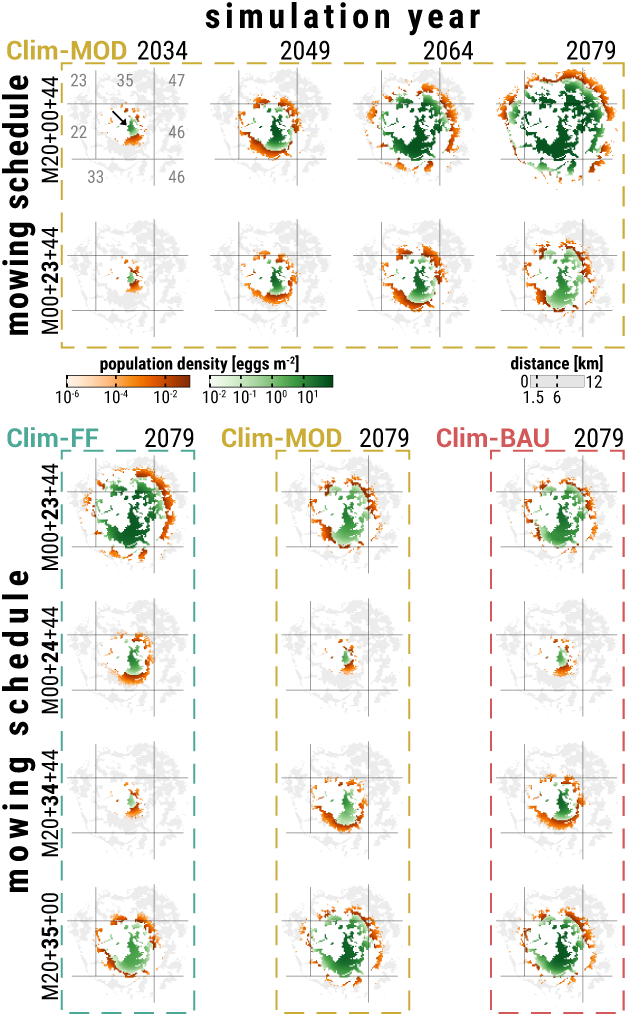
Spatial distribution of an LMG population dispersing from a singular source habitat in the center (black arrow in top left subplot) of climate cell 34. Each colored cell within the 20 subplots represents the population density in eggs m^-2^ (mean over 10 replicates) in a 6.25 ha grassland habitat inside a 16 km radius of the source habitat at the end of a simulation year. GREEN cells are considered habitats with an established LMG population, i.e., an imago life stage density ≥ 0.002 individuals m^-2^ during a simulation year; ORANGE cells represent unestablished populations at the cutting edge of the dispersal process, i.e., a imago life stage density < 0.002 individuals m^-2^; GREY cells are habitats that are reachable at the end of the 60 year simulation time in case of ideal conditions with minimal disturbances (cf. top right subplot); WHITE areas were either unreachable or do not qualify as grassland. The grey grid lines delineate the climate cells from each other, where the grey numbers in the top left plot label the ID of the respective climate cell. The top two rows show – from left to right – the chronological LMG distribution progress after 15, 30, 45 and 60 simulation years exemplarily for the Clim-MOD climate change scenario (CCS), where the first row is the progress under ideal conditions (low-impact mowing) and the second row in an environment disturbed by mowing schedule M00+23+44 (mowing in calendar week 23 instead of 20). Each of the 12 plots in the three bottom columns depict the LMG distribution at the end of the final simulation year 2079 depending on the CCS Clim-FF (first column), Clim-MOD (second) and Clim-BAU (third) as well as the applied mowing schedules M00+23+44 (first row), M00+24+44 (second), M20+34+44 (third) and M20+35+44 (last)

Figure 7 provides an overview of the dispersal success in terms of *maximum established distance* at the end of a simulation run depending on CCS, mowing schedule and source habitat. In an undisturbed environment (M00), established populations on average reached distances of roughly 14,000 m and at most up to 40,000 m. More importantly the figure highlights, in which simulations population establishment basically remained restricted to the dispersal radius of the source habitat (Figure 7, dots below black horizontal dashed line). This was the case for virtually all of the regions (or rather source habitats) when mowing schedules M20+26+44 (M26) to M20+31+44 (M31) were applied. Only outside of this time window, i.e., early mowing before calendar week 26 or late mowing after week 31, dispersal could be successful to some extent, depending on region and CCS. As described above on the example of climate cell 34, early mowing schedules were in favor of the Clim-FF scenario while late mowing was in favor of Clim-MOD and especially Clim-BAU.

**Figure 7:**
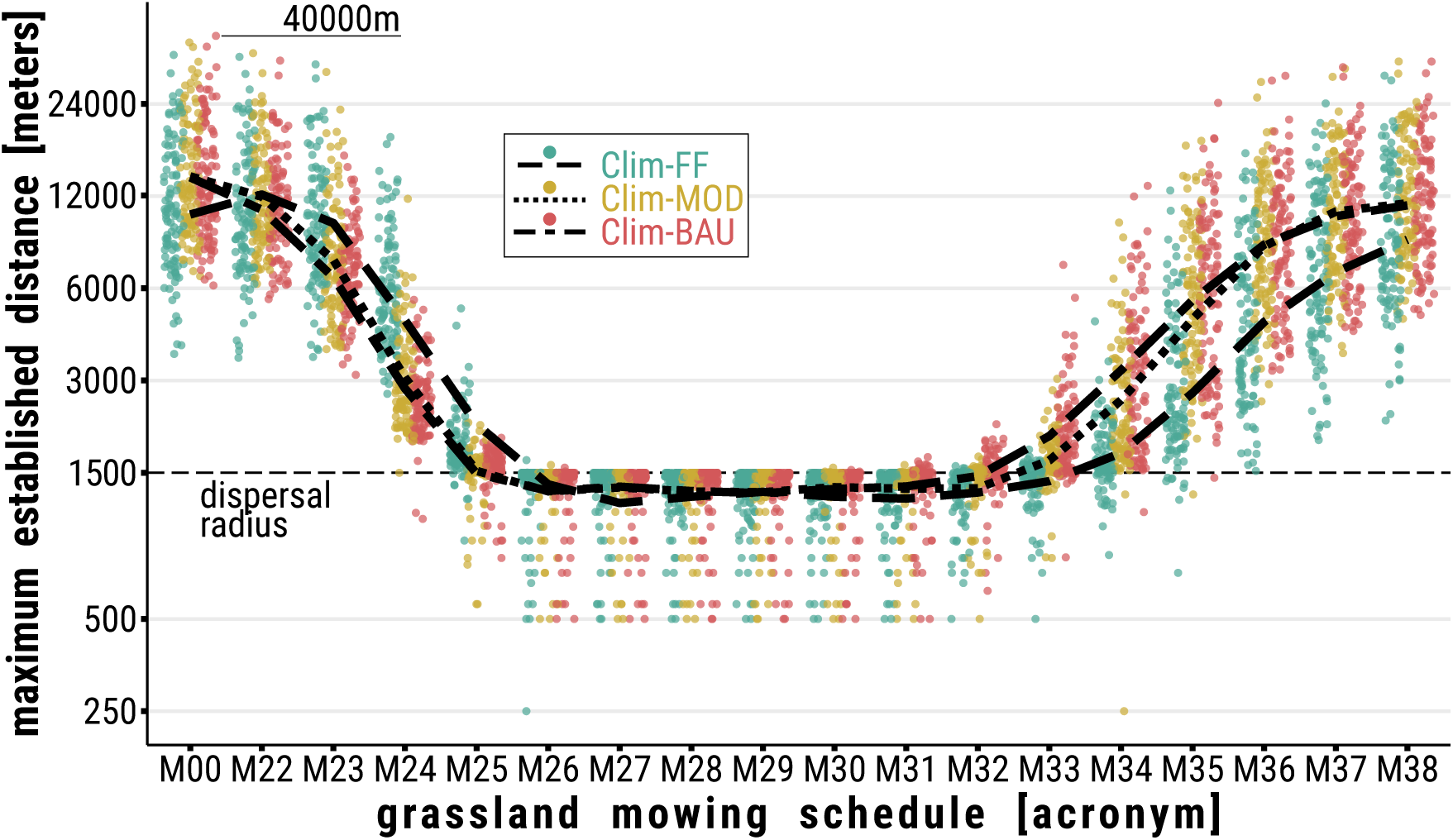
Distance in meters (y-axis) from a source habitat to the most distant established population in its neighborhood by the end of a simulation run in 2079 depending on mowing schedule (x-axis) and climate change scenario (CCS; green=Clim-FF, brown=Clim-MOD, pink=Clim-BAU). Each colored dot represents this distance value for either one of the 107 initial populations (or climate cells) averaged over ten replicates. The trend lines are distinguished by CCS (dashed=Clim-FF, dotted=Clim-MOD, dash dotted=Clim-BAU) and follow the mean for the distance value over the 107 cells in the study region depending on the applied mowing schedule. The horizontal black dashed line marks a distance of 1,500 m. Populations established directly through (LDD) from the source habitat were omitted in the calculating to avoid misleading maxima outside the dispersal radius

Figure 8A depicts the spatial distribution of results on the example of the *maximum established distance* by the end of simulation runs for scenario Clim-FF and a low-impact mowing schedule. We compared the spatial results of evaluation parameters *population size, population density* and *maximum established distance* for all three CCS and a low-impact mowing schedule to (a) the *population density* of simulations without dispersal and (b) the *grassland cover* within dispersal radius (1,500 m) of the initial populations. Table 3 provides the 18 corresponding correlation coefficients (ρ). Please refer to Supplement S2 for illustrations of spatial simulation results of the evaluation parameters used in Figure 8 and Table 3 depending on CCS, and to Supplement S5 for scatter plots depicting the parameter correlation resulting in the coefficients of Table 3.

**Table 3:**
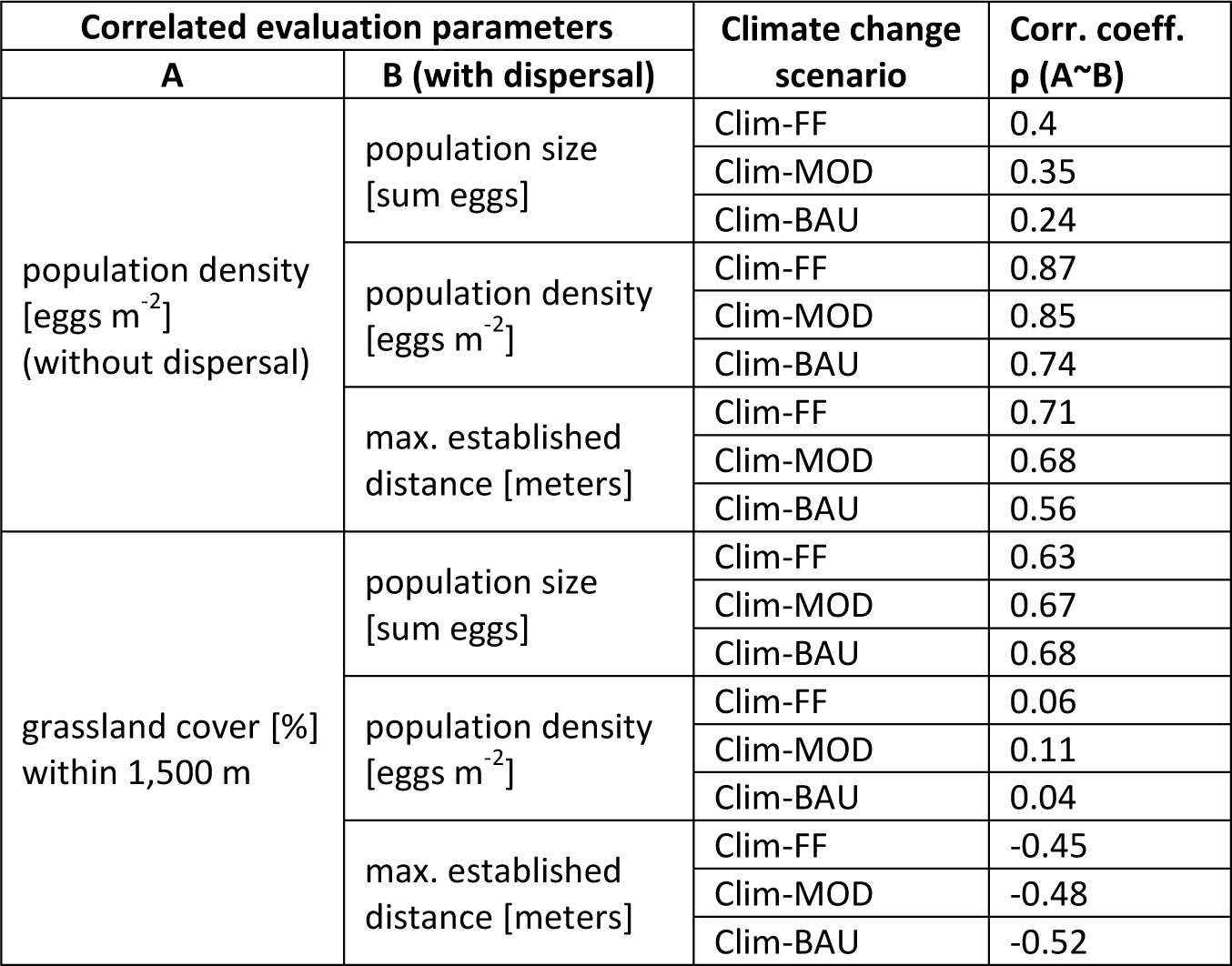
**Coefficients ρ (last column) from correlation of evaluation parameters A (first column) and B (second column) dependent on climate change scenario (third column). Correlated values stem from summarized results of 107 regionally different and independent simulations (with and without dispersal), each with 10 replicate runs (5 without dispersal), and data of grassland cover around the initial population within the respective region**.

**Figure 8:**
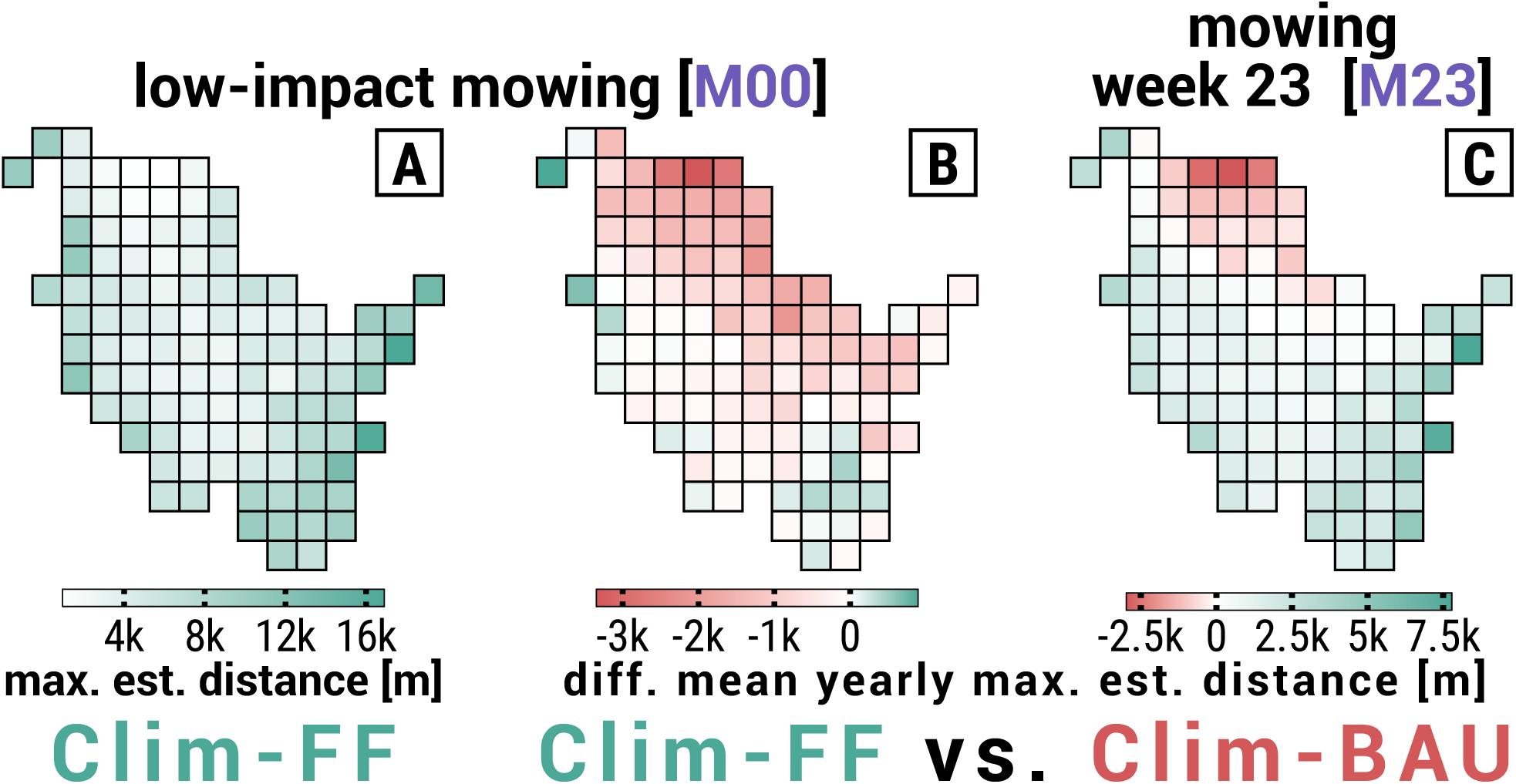
Spatial distribution of maximum established distance [meters] exemplary for low-impact mowing [M00] and climate change scenario (CCS) Clim-FF (subplot A), and difference (delta) in dispersal success in terms of mean yearly maximum establishment distance [meters] between CCS Clim-FF and Clim-BAU exemplary for (B) low-impact mowing [M00] and (C) mowing in week 23 [M23]. See Table 1 for detailed timing of mowing schedules. Each of the 107 cells per subplot represent INDEPENDENT simulation runs, or rather their mean over 10 replicate runs, and depict the results of the dispersal process from a SINGLE initial population in the center of a cell. In subplots B and C, values were determined per replicate by subtracting the yearly established distance of the CCS mentioned second from the CCS mentioned first in the header and then calculating the replicate mean of absolute delta. Further, the cells’ background colors highlight which of the respective CCS on average shows the higher differences during the 60 simulation years, where a LIGHTER color represents lower average difference. GREEN cells are in favor of Clim-FF and PINK cells in favor of Clim-BAU. Please refer to Supplements S2 and S3 for distribution maps of all CCS and mowing schedules.

We found that the *population density* resulting from simulations without dispersal is positively correlated with all evaluation parameters coming from simulations with dispersal, yet to different extents. There was a high correlation between the two *population densities* and the expected *maximum established distance* could be estimated reasonably well from simulations without dispersal, but *population size* could only be derived moderately. Generally speaking, the correlation was always higher for less severe CCS.

Regarding the effect of spatial patterns on the dispersal success, there were only correlations expectable within the domain of the HiLEG model: independent of the CCS and mowing schedule, *grassland cover* positively correlated with *population size* but did not correlate with *population density*; due to the LDD process, large *maximum established distances* were achieved in regions with low grassland cover, resulting in a moderate negative correlation.

Though the correlation with *grassland cover* was similar for all CCS, there are regional patterns in SH that only became apparent when looking at the difference (delta) of dispersal success between CCS depending on the considered mowing schedule. Figures 8B-C show the difference in *maximum established distance* averaged over the whole simulation run exemplary for the delta between Clim-FF and Clim-BAU in scenarios of low-impact mowing (Figure 8B) and mowing in week 23 (Figure 8C). The examples highlight, on the one hand, that there are in fact regions better suited under less severe climate change and others better suited under more severe climate change. On the other hand, they show that the suitability can spatially shift depending on the mowing schedule.

There are other regional patterns in SH that occurred repeatedly when comparing the delta of simulation results between climate scenarios (see Supplement S3 for comprehensive illustration). The state was divided into two parts by a virtual diagonal line running from the Northwest to the Southeast. Regions in the Northeast usually allowed a greater dispersal success for more severe CCS with occasional exceptions in the eastern part. Southwest regions tended to allow more successful dispersal for less severe CCS. Low-impact and late season mowing resulted in the Clim-MOD scenario allowing higher dispersal success than the Clim-FF scenario throughout all regions. Overall the largest deltas in dispersal success occurred in the upper Northeast, along most of the west coast and in the southeastern regions.

Please note again that the regional deltas depicted in Figures 8B-C and Supplement S3 represent the mean deltas over the full simulation runs and thereby do not capture variations in deltas during the simulations. Therefore, mean deltas can be low although there is in fact a clear difference between the CCS in many years. Since the same CCS does not always lead to larger dispersal success, these differences may, however, cancel each other out. Furthermore, the mean deltas can be different from the final deltas at the end of the simulation runs, but as time series of deltas are affected by many peculiarities, discussing them in detail would not easily lead to general insights. Overall, we found the mean deltas displayed in Figures 8B-C and Supplement S3 to be good indicators for identifying the CCS allowing for better dispersal success, and chose them deliberately to avoid focusing only on the final dispersal success.

## 4 Discussion

The implications of the findings described in Section 3 that we obtained by extending the HiLEG model by a dispersal process will be discussed below.

### 4.1 LMG is a fairly good but slow disperser

We found that in the simulations, the average distance for establishing new populations was roughly 467 m within two years, which is comparable to the dispersal distances of 400 m within two years reported by Marzelli (1994). The difference between model and empirical observation may have several reasons. In our simulations, the conditions were considered ideal other than in the natural environment of the field study. Also, the simulated grassland plots had a fixed size thus dispersal steps were restricted to a minimum distance of 250 m; excluding shorter distances certainly shifted the mean dispersal distance to larger values.

Regarding the applied dispersal radius, a ‘mark and recapture’ study by Malkus (1997) only found specimens in a maximum distance of 624 m. Other studies using genetic markers, however, estimated maximum dispersal distances of 3,000 m (Van Strien, 2013) or calculated a connection distance between two populations of up to 3,264 m (Keller, 2012). The value of 1,500 m we adapted from Griffioen (1996) thus appears to be plausibly middle ground, at least in an environment of high grassland cover.

In a more fragmented landscape, mobility of the LMG must be considered in a different light. Bönsel and Sonneck (2011) conducted a triannual ‘mark and recapture’ study for an isolated, yet stable habitat and found that none of the LMG specimens migrated to either one of the four suitable 1, 500 to 9,000 m distant study sites, concluding a low dispersal activity in a highly fragmented environment. Marzelli (1994) observed, on the other hand, that the LMG is able to cross unsuitable areas of 300 m, allowing dispersal at least in a slightly fragmented landscape. Regarding the LDD process there is no quantification of its parameters in the literature but evidence that it occurs at least occasionally: individuals of the LMG were found on an island 10 km offshore with the next known onshore population about 16 km away (Oppel, 2005) while flight was observed where individuals ascended more than 20 meters into the air and out of sight (Trautner and Hermann, 2008).

With this knowledge we implemented LDD using the values of regular dispersal, because it already includes parameters that reduce dispersal probability with distance between grassland plots. We restricted LDD to landscapes where the distance between grassland plots is larger than 1,500 m (see Section 2.5) to reflect the assumption that nearby grassland, if present, is prioritized for colonization or as stepping stone for farther dispersal. The effects on LMG dispersal by potentially unbridgeable barriers such as highways or forests and climate conditions such as wind direction were ignored in our simulations, as we focused on studying the interplay of climate change relevant parameters and mowing schedules. However, we increased dispersal mortality with decreasing grassland cover (Section 2.5) to account for unsuitable conditions in fragmented landscapes.

The fact that the dispersal success remained within reasonable bounds despite the applied simplifications and estimations provides the confidence to consider our simulation results of applied mowing schedules valid as well. More importantly, the rather short projected dispersal distances, especially in a disturbed environment, reinforce the choice for our approach to study the development of individual populations at regional or even local level.

### 4.2 Climate change facilitates the expansion in North SH state

The regional patterns of dispersal success in an undisturbed environment for each CCS separately are qualitatively very similar to each other. Some of the patterns even follow climatic conditions already largely found in simulations without dispersal that will be discussed later. Comparing the deltas between evaluation parameters of the three CCS pairs (Clim-FF vs. Clim-MOD, Clim-FF vs. Clim-BAU, Clim-MOD vs. Clim-BAU), however, revealed regional differences (Figures 8B-C, Supplement S3) with possible implications for climate dependent management strategies.

With some exceptions in the Southwest, the LMG widely benefits from climate change in SH. This again confirms our previous findings (Leins et al., 2021) as well as the results from similar studies (Poniatowski et al., 2018; Trautner and Hermann, 2008). A moderate climate change (Clim-MOD) would be beneficial for the LMG in the whole study region. In case of severe climate change (Clim-BAU) only the western coastal regions would be worse off but the conditions running from North to East of the study region would improve the most in this scenario. The latter is relevant for two reasons.

First, the northeastern interior of SH is the region where currently most of the inhabited LMG habitats are located (Figure 2B, orange circles). With climate change in mind, conditions would thus improve the most particularly for these existing populations. Second, the northern regions are currently the most difficult terrain for the LMG, where hardly any populations are found. Although it is going to remain the least suitable region climatically (Figure 8A, Supplement S2), it would improve the most in the more severe CCS (Figures 8B-C, Supplement S3) and as a result could facilitate LMG expansion to the North. Poniatowski et al. (2020) already found that many grasshopper species including the LMG expand their range due to global warming. Especially a climate change driven northern range shift is often discussed for – among other species (Van der Putten, 2012) – insects as well (Stange and Ayres, 2010) and was particularly shown for several grasshopper species (Poniatowski et al., 2018).

### 4.3 Higher grassland cover allows larger population size

The second region currently scarcely populated by the LMG is the west coast of SH and its interior. Only in the southern and central parts along the coast, a few populations are found. This is despite the fact of it having a high grassland cover (Figure 2B) and that our simulations showed suitable conditions of potentially high *population density* throughout the region (Supplement S2) even with mild climate change (Clim-FF). Especially on the central west coast with highest grassland cover (Figure 2B) that is notably correlated with high *population size* (Table 3, ρ between 0.63 and 0.68) there are currently no known LMG populations (Figure 2B, missing orange circles).

Even though it is reasonable that the higher availability of grassland allows a larger number of populations – and thus higher overall *population size*, the reason for the sparse presence of the LMG is apparently neither the climatic nor the biotic conditions but likely the fact that the northwestern region of SH has the state’s highest percentage of agricultural land, with more than 74% (Statistisches Amt für Hamburg und Schleswig-Holstein, 2021).

The negative correlation of grassland cover with *maximum established distance* (Table 3, ρ=-0.45 to -0.52), on the other hand, can be ignored within the domain of the model. It is due to the fact that especially in fragmented landscapes the LDD process applies, allowing for above-average dispersal distances.

The main problem for all of the currently (mostly) uninhabited regions, especially in the Northwest of the study region, is the relatively large distance to the closest established LMG populations (Figure 2B, orange circles). Measures to assist the LMG to migrate to these regions are likely to depend on local constraints and can only be partly derived from the results of the present study. We will nevertheless address potential management strategies later in the discussion.

### 4.4 Mowing slows down dispersal but still allows it up to a threshold

The key to all above considerations is the right timing of grassland mowing because it is one of the critical factors for the dispersal success (Figure 7) and survival of LMG populations. In our preceding study, it was unclear how to interpret the diminishing effect of mowing on the LMG’s lifetime during summer and early autumn (Leins et al., 2021, Figure 9). From the results of the present study we learn that, while population development might become increasingly restricted when mowing up to calendar week 25 and down to calendar week 32 (Figure 7), it still allows (slower) dispersal and establishment outside of a source habitat’s dispersal radius (Figure 6).

It is important to bear in mind that there is a spillover effect within the dispersal radius due to the unrealistically undisturbed source habitats and that the mowing dates should be interpreted in relative, not absolute terms (Section 2.5). Yet, the resulting dates provide valuable insight for potential management strategies in agricultural grasslands, because it means that there are ways of supporting LMG establishment and dispersal while allowing economically beneficial land use. Especially the early mowing weeks of late spring and early summer are of relevance here, because they produce the best yields for farmers (Gerling et al., 2022).

Furthermore, with a minor climate change (Clim-FF) mowing is even less problematic for a longer period of time before summer in most parts of SH than it would be with the more severe scenarios Clim-MOD and Clim-BAU (Figure 8C, Supplement S3). Such a longer period of unproblematic mowing with the Clim-FF scenario could be highly relevant when considering the implications of climate change for the species. Yet, this assessment is only valid under the assumption that beginning and duration of vegetation growth does not shift in the same way as the life cycle development of the LMG. In order to examine this hypothesis, dynamic models of grassland growth and management decisions would indeed be useful.

We discussed above that from climate change alone the LMG would benefit in all (Clim-MOD) or most (Clim-BAU) parts of the study region. However, SH is with an agricultural area of 68.5% (Statistisches Bundesamt, 2021) the most intensively farmed state in Germany (50.6%). In such an environment, the LMG would thus be better off in case of minor climate change or none at all. It would still require measures reducing intensive grassland use to allow the LMG to thrive and expand.

### 4.5 Spatially stationary simulations as indicator for suitable regions

As pointed out above, simulations without dispersal could already help identifying regions that in principle support LMG development and highlight the general implications of disturbances such as mowing on LMG populations. Depending on the evaluation parameter of simulations with dispersal, we found correlations of different extent with the *population density* stemming from the spatially stationary simulations (Table 3, top rows): Less surprisingly, the *population density* of established habitats within a region highly correlated (ρ=0.74-0.87), because it is mainly driven by regional climate conditions. Furthermore, there is only little correlation (ρ=0.24-0.4) with the *population size* as it depends more on grassland cover (see above).

Interestingly, however, there is a noticeable positive correlation (ρ=0.56-0.71) with the *established distance*, especially for the less severe CCS. Therefore, results from simulations of stationary populations could already be a good indicator for the development – and even the general ability to disperse – of species such as the LMG in a regional context. Within the domain of our model, high spatial resolution thus is not the key factor for broadly identifying (climatically) suitable regions. This is a useful insight, especially because simulations without dispersal require less information about a target species and have a much shorter runtime.

The actual development and distribution of a dispersing population could, however, change both qualitatively and quantitatively depending on the spatial patterns and climatic gradients within a region. Particularly in combination with disturbances, the introduction of the dispersal process delivered valuable information: First, mowing schedules that seemed highly problematic in spatially stationary simulations could still allow (reduced) dispersal success. Second, the grassland cover could change the implications of a region’s general suitability, because it might either hinder dispersal in fragmented landscapes of otherwise suitable conditions (Bönsel and Sonneck, 2011) or improve population establishment with high cover (Table 3, bottom rows) and thus a larger number of refuges.

The relevance of a dispersal process and spatial patterns might increase further if other factors are additionally considered. A mechanistic dispersal process (Vinatier et al., 2011) instead of the present statistical approach could, for instance, result in a more directed preference for neighboring habitats. This effect would especially apply, if (micro) climate was more heterogeneous or less gradually distributed in a study region. Similarly, a more realistic distribution of varying land use (timing) or other detrimental/beneficial environmental conditions could hinder/promote regional dispersal attempts. Furthermore, considering the effects of spatial patterns such as fragmentation on, for example, dispersal and mortality rates or extinction events might further change species distribution. Ways of including some of these mechanisms into the model to analyze the dispersal success in more detail are addressed at the end of the discussion.

### 4.6 Management decisions require expertise on a regional level

Overall our results showed that there is no universal formula for protecting and supporting LMG populations in cultivated grasslands of North Germany, just a tendency in the implications of (future) climate change and a coarse window of unsuitable mowing schedules. Though a broad approach of rather low-impact land use could be applied using our results, it would probably not be feasible on a large spatial scale, because such measures of poor spatial targeting have proven to be less effective (Meyer et al., 2015). At the same time, the uncertainty of climate change makes robust and cost-effective conservation policies necessary (Drechsler et al., 2021). Therefore, management decisions require expertise on a regional or even local level and should remain flexible, especially in grasslands (Joyce et al., 2016), to be able to react to the severity of climate change (Hulme, 2005). As mentioned above, our approach regarding management strategies is too broad to recommend specific local measures of LMG conservation, but we want to discuss below some suggestions that can nevertheless be derived from our results. We focus mainly on such suggestions, which could also be addressed with the HiLEG model using an adapted simulation setup in a follow-up study.

Heller and Zavaleta (2009) compiled a ranked list of recommendations for management strategies and conservation planning in the face of climate change. The authors recommended the integration of climate change monitoring into conservation planning, particularly in terms of management schedules. We can follow this recommendation because our results show, even for a small state like SH, that regional differences occur due to timing of mowing and severity of climate change (Figures 7 and 8). Coupling the mowing schedules to local climate conditions in some grassland plots could be one way to simulate such monitoring and help clarify its effect on the species dispersal success, especially if the coupled schedule falls outside the unsuitable time window. Additionally simulating aggregated grassland plots of suitable timing could help analyzing the effect of creating refuges of larger size, which is another recommendation stemming from the ranked list.

Our results and data further suggest that, despite the regional differences already discussed above, there are other factors worth considering for the allocation of conservation planning. Obviously, conservation measures should be focused on regions where the target species is already present, at least if the objective is not its reintroduction. In case of the LMG in SH (Figure 2B, orange circles), this does not necessarily match the regions suited best for the species, as discussed in Sections 4.2 and 4.3 regarding varying climate conditions and grassland cover. The spatial maps of dispersal success we compiled for this study (Figure 8, Supplements S2 and S3) give a first idea of both, potentially promising regions and others that would require precaution in management.

Along with the occurrence data, the spatial maps also highlight potential dilemmas for conservation planning, namely that the regions whose suitability would likely develop the most are sparsely inhabited, and that the most populated regions also have fragmented grasslands. While developing uninhabited, yet connected grasslands for species such as the LMG might be a long-term management objective, the species’ low dispersal speed could prove problematic for conservation planning in fragmented landscapes. Creating undisturbed satellite habitats in terms of metapopulation theory (Hanski, 1999; Levins, 1969) would in theory promote occasional (long distance) dispersal and compensate for local extinction, but the benefits of such metapopulation dynamics in fragmented landscapes for slowly dispersing species are controversial (Bönsel and Sonneck, 2011).

Altogether, simulating and exploring the prospects of different measures in different regions before actually implementing them is advisable. Such simulations could easily be conducted with only minor modifications to the HiLEG model and deliver valuable insights to conservation agencies for the protection of local LMG populations. With the right set of parameters, the model could additionally be adjusted for the life cycle of other species to achieve a broader picture of the implications for disturbed grassland communities in the face of climate change. However, as grasshopper species like the LMG are considered indicators for the quality of grassland biotopes (Báldi and Kisbenedek, 1997; Keßler et al., 2012; Sörens, 1996), the analysis of single species already gives a good idea of the implications for such a community.

## 5 Conclusion

The introduction of dispersal into the highly resolved, yet formerly non-spatially-explicit HiLEG model provided valuable insights regarding the implications of anthropogenic disturbances for the large marsh grasshopper (LMG) under different climate change scenarios. Our study reconfirmed that the LMG in principle benefits from a moderate climate change in temperate regions and was also helpful in unraveling the impact of grassland mowing schedules that were previously unclear. Namely that some of the schedules, despite inhibiting population development, could still allow species dispersal to some extent. It depends on the regional conditions and severity of climate change which mowing schedules this mainly involves.

A milder climate change permits a longer mowing period in the beginning of the season and is more beneficial in the southwestern parts of Schleswig-Holstein (SH). This is an important observation, because early mowing provides the highest yields for farmers. More severe climate change, on the other hand, allows for earlier resumption of mowing after summer, especially outside the western interior of the state. Grassland cover only plays a minor role in the development of the LMG, though a high cover facilitates population establishment within a region.

However, many of the regions that might either improve the most under climate change (North SH) or offer high grassland cover (West SH) are currently scarcely populated by the LMG. Assisting the grasshopper in migrating to those regions will require flexible management decisions on a local level, especially because the key factors hindering the LMG from thriving are anthropogenic (thus controllable) disturbances such as grassland mowing. Improving these practices might benefit other (insect) species as well, because of the LMG’s role as indicator for the quality of grasslands. However, this would need to be tested as the life cycles and their most sensitive phases can vary widely between species. HiLEG was designed to be adaptable for other grassland insect species as well (Leins et al. 2021).

In the above discussion, we identified four factors that we recommend to consider for such regional management decisions: (1) the development of climate conditions (when and in which region to apply measures); (2) the grassland cover (size, number and distribution of refuges); (3) the existence of LMG populations (habitats prioritized for protection); and (4) the use of simulation models (identifying suitable measures before implementing them).

The results from both the present and previous study, with and without consideration of dispersal, provided a number of key indicators for potential management strategies in cultivated landscapes. With their input alone, a reasonable protection of grassland (insect) species such as the LMG can be achieved. To further assist stakeholders on a regional level in their decision for viable management strategies, a more realistic or rather heterogeneous integration of disturbances could be of relevance. Such a follow-up study can easily be performed with only minor modifications to the HiLEG model along with the matching set of parameters – eligible for other target species as well.

## Supporting information

ODD model description

Illustration of dispersal success

Illustration of differences in dispersal success

Specs of climate and grassland cells

Illustration of correlation between evaluation parameters

## 6 Acknowledgements

This work has been carried out within the project Ecoclimb (www.b-tu.de/en/ecoclimb), funded by the German Federal Ministry of Education and Research (grant no. 01LA1803B). We especially thank Björn Schulz from the *Stiftung Naturschutz Schleswig-Holstein* for providing further expertise and literature about the dispersal behavior of the large marsh grasshopper. Furthermore, we thank two anonymous reviewers for their helpful comments and suggestions.

## 7 Competing Interests Statement

There is no known competing interest that could have influenced the present work.

## 8 Author Contributions section

JL: concept, design and implementation of the model; acquisition and processing of input data; analysis and interpretation of the simulation output; drafting and visualization of the article. MD: model concept; data interpretation; review and editing of the article. VG: data interpretation; drafting and editing of the review.

## 9 Data Accessibility Statement

Open access model code, executables and required input data are available via GitLab (git.ufz.de/leins/hileg). Output data generated by the HiLEG simulation runs for this study can be found at ufz.de/record/dmp/archive/11898/en/ and the aggregated data used for analysis and illustration at ufz.de/record/dmp/archive/11896/en/ (with dispersal) andufz.de/record/dmp/archive/11899 (without dispersal). A representation of the data used for mapping and weighing climate and grassland cells for bilinear interpolation can be found at ufz.de/record/dmp/archive/11900. The model release version used to run the simulations for this study is located at git.ufz.de/leins/hileg/-/tree/v1.4 along with detailed descriptions of the input data and simulation parameters.

## Footnotes

1 Consortium for Small-scale Modeling in Climate Mode

2 Irish Centre for High-End Computing

3 provided by *Landesamt für Landwirtschaft, Umwelt und ländliche Räume* via our project partner *Stiftung Naturschutz Schleswig-Holstein*

4 HiLEG GitLab repository: git.ufz.de/leins/hileg

## Notes

https://www.ufz.de/record/dmp/archive/11896/en/

https://git.ufz.de/leins/hileg

https://www.ufz.de/record/dmp/archive/11898/en/

https://www.ufz.de/record/dmp/archive/11899/en/

https://www.ufz.de/record/dmp/archive/11900/en/

### Competing Interest Statement

The authors have declared no competing interest.

