## Supplementary material for "Large scale PVA modelling of insects in cultivated grasslands: the role of dispersal in mitigating the effects of management schedules under climate change": ODD model description

The model description below follows the ODD (Overview, Design concepts, Details) protocol (Grimm et al., 2020, 2006). It includes the original model description (M1 and black text) used to study the effects of land use and climate change on spatially stationary populations of the large marsh grasshopper (LMG) in Northwest Germany (Leins et al., 2021). Furthermore, it describes the extensions made to the original model (M2 and green text) for the present study to explore the additional effects on the species when adding dispersal in an environment of higher spatial resolution.

### 1 Purpose and patterns

M1: The *PURPOSE* of the model HiLEG (*Hi*gh resolution *L*arge *E*nvironmental *G*radient) is to answer the following questions regarding populations of the large marsh grasshopper (LMG) *Stethophyma grossum* (Linné 1758) in Northwest Germany: (1) How do population density and viability shift regionally, given different climate change scenarios? (2) Which mowing schedule has the least negative impact on the overall population density and viability in the study region? (3) Does the mowing impact severity depend on the spatial location (with its specific climate)?

The empirical patterns used to ensure that the model is realistic enough for its purpose are observed features of the life cycle and their sensitivity to environmental conditions, which were taken from literature. These patterns were used for the model's design. Model output in terms of population structure, densities and persistence were not compared to data, as such data are sparse. Therefore, all model predictions are relative, not absolute. The model was implemented in C++. The source code of the model implementation and the input files used for the simulations runs are available via a GitLab repository[1].

M2: The purpose of the model extension is to study the additional effect of dispersal on the LMG in a North German environment of realistic grassland distribution with higher spatial resolution to answer the following questions: (1) Are there (regional) differences in dispersal success depending on climate change scenario? (2) Is the success of dispersal additionally affected by spatial patterns such as grassland cover? (3) Can dispersal compensate for otherwise detrimental grassland mowing?

The species' dispersal metrics were taken from literature and known LMG habitats (main manuscript, Figure 2B, orange circles) adapted from survey data[2] gathered in the years 2000 to 2016 were used to analyze some implications of regional effects. Other measures of dispersal success are relative, not absolute, due to a lack of relevant data.

### 2 Entities, state variables and scales

M1: The model has the following entities: *Grid Cells* (defining environmental conditions) and *Population* per grid cell comprised of *Life Stages* which are comprised of age-distinguished *Cohorts*. *Flows* are auxiliary entities that manage the density transfer between *Life Stages* or their loss through mortality.

---

[1] HiLEG GitLab repository: https://git.ufz.de/leins/hileg

[2] Provided by *Landesamt für Landwirtschaft, Umwelt und ländliche Räume* via our project partner *Stiftung Naturschutz Schleswig-Holstein*

M2: Instead of *Grid Cells* the model extension has two separate entities *Climate Cells* (defining large scale climate conditions in a 12 x 12 $km^2$ region) and *Grassland Cells* (defining environmental conditions, e.g. interpolated climate values, on a scale of 250 x 250 $m^2$). A *Grassland Cell* contains an initially empty *Population* entity and is considered *inhabited* if the *Population* has a non-zero *density*. Otherwise it is considered *uninhabited*. The *Flow* entities additionally connect *Grassland Cells* and handle the density transfer between their *Populations* during dispersal.

M1: The LMG develops through three main *Life Stages* during a year (cf. main manuscript, section 2.2). Following Ingrisch (1983) and Wingerden et al. (1991), we divided the egg/embryo stage into pre-diapause, diapause and post-diapause development (called embryo hereafter) to account for the clutch's different susceptibility to climate conditions in autumn, winter and spring.

This subdivision yields five *Life Stages*: 1 pre-diapause, 2 diapause, 3 embryo, 4 larva, 5 imago. Stages 1 to 3 occur below ground, stages 4 and 5 above ground. Furthermore, stages 2 and 4 can have multiple *Cohorts*, to allow survival over several years in case of conditions during winter that unsuitable for development, and to account for different temperature-driven development speed depending on hatching date.

M2: Dispersal only occurs between the *Populations'* imago *Life Stages* of *Grassland Cells* within a defined neighborhood.

M1: Table S1-1 provides an overview of the model's entities and their state variables. The *Population* consists of several *Life Stages* and is characterized by the *coordinates* of its grid cell and a *minimum density* [individuals $m^{-2}$]. The *Population*'s *density* and *aboveground density* [both in individuals $m^{-2}$] are calculated from its *Life Stages*' *densities*.

A *Life Stage* has a *name*, consists of one or more *Cohorts* and supplies a *maximum age* [days] for each of those *Cohorts*. *Cohorts* exceeding the *maximum age* or falling below the *minimum density* are considered extinct. Furthermore, a *Life Stage* has a *flag* informing whether it occurs below or above ground. The *Life Stages*' *density* [individuals $m^{-2}$] is calculated from its *Cohorts*' densities and the amount of *gain* [individuals $m^{-2}$] from the incoming transfer *Flow* of its preceding *Life Stage*.

*Cohorts* are distinguished by an *ID* and have an *age* [days] and a *density* [individuals $m^{-2}$]. They have a *development progress* given as a ratio $\in [0,1]$ that defines if and how density is transferred to subsequent *Life stages*. The *development progress* is used in two different ways depending on the external *Influence* (section 7.1) associated with the *transfer Flow*: (1) it is directly affected by external *Influences* and contributes to a *Flow*'s density transfer to the subsequent *Life Stage* if it reaches a value of 1 (section 7.4); or (2) it is used within stochastic *Influences* to determine the probability and extent of contribution to the *flow rate* (section 7.1, *Binominal Climate,* M2: renamed from *Factor*; Pseudocode S1-2). Without external *Influences*, the *development progress* defaults to a value of 1.

**Table S1-1: Overview of the model's entities (first column) and their state variables (second column). The text in parentheses of columns one and two represents the entity's or state variable's symbol when used, e.g., in equations. The third column gives the (initial) value(s) of the state variables, the fourth column gives their units (if any). In the fifth column, a brief description of the state variable is provided. The indexes for location and time step that distinguish entities and their dynamically changing states are implied and not explicitly specified in the identifier. The parameter $A_{hab}$=62,500 m$^2$ is the habitat size modeled for a population.**

| Entity (symbol) | State variable (symbol) | Value(s) | Unit | Description |
|---|---|---|---|---|
| Population ($P$) | coordinate ($coord_{x,y}$) | $x, y \in [1,36]$ | | Index of rotated pole grid coordinates (see main manuscript) |
| | set of life stages ($P^{stages}$) | $\{S^{pre}, S^{dia}, S^{emb}, S^{lar}, S^{ima}\}$ | | Distinguished life stages of the target species |
| | set of flows ($P^{flows}_{type}$) | | | A set of all flows associated with this Population distinguished by their flow type (see below) |
| | density ($dens^P$) | $\sum\{dens^{pre}, dens^{dia}, dens^{emb}, dens^{lar}, dens^{ima}\}$ | ind. m$^{-2}$ | Summed life stage densities |
| | aboveground density ($dens^P_{above}$) | $\sum\{dens^{lar}, dens^{ima}\}$ | ind. m$^{-2}$ | Summed density of the aboveground life stages |
| | minimum density ($dens_{min}$) | $1/A_{hab}$ | ind. m$^{-2}$ | Minimum of one individual per habit |
| Life Stage ($S^{name}$) | name | $\in \{pre, dia, emb, lar, ima\}$ | | Name of the distinguished life stages |
| | set of cohorts ($S^{name}_{cohorts}$) | $\{C^m, .., C^n\}, \{m, n\} \in \mathbb{N}$ | | Distinguished cohorts associated with the life stage |
| | density ($dens^{name}$) | $\sum\{dens^m, .., dens^n\}, \{m, n\} \in \mathbb{N}$ | ind. m$^{-2}$ | Summed densities of the associated cohorts |
| | gain ($gain^{name}$) | | ind. m$^{-2}$ | Summed total flow amount of the incoming flows (sections 7.3, 7.4) |
| | aboveground flag ($above^{name}$) | $\in \{TRUE, FALSE\}$ | | Boolean flag defining whether the stage occurs above ground |
| | maximum age ($age^{name}_{max}$) | $\in \{210,1700,120,90,120\}$ | days | Maximum age of the associated cohorts |
| Cohort ($C^{ID}$) | ID | $\in \mathbb{N}$ | | Unique cohort identifier |
| | density ($dens^{ID}$) | | ind. m$^{-2}$ | The cohort's individual density |
| | age ($age^{ID}$) | | days | Age in days since cohort creation |
| | development progress ($prog^{name}_{ID}$) | $\in [0,1]$ | | The ratio of development compared to full development |
| Flow ($F^{name}_{type}$) | type | $\in \{trans, repr, mort, disp\}$ | | Defines how the flow is processed |
| | life stage of origin (orig) | $\in \{S^{pre}, S^{dia}, S^{emb}, S^{lar}, S^{ima}\}$ | | Life stage used to determine the amount of flow |
| | life stage of destination (dest) | $\begin{cases} mort & \text{if } type = mort \\ \in \{S^{pre}, S^{dia}, S^{emb}, S^{lar}, S^{ima}\} & \text{otherwise} \end{cases}$ | | Life stage receiving the amount of flow, or the amount of mortality |
| | set of influences ($I^{name}_{type}$) | | | Environmental drivers associated with this flow |
| | base flow rate ($rate^{name}_{type}$) | | day$^{-1}$ | Daily per capita base flow rate |
| | dynamic flow rate ($dyn^{ID}_{type}$) | | day$^{-1}$ | Daily per capita flow rate per cohort in the life stage of origin |
| | current flow amount ($amount^{ID}_{type}$) | $dens^{ID} \times dyn^{ID}_{type} \times day$ | ind. m$^{-2}$ | Amount of density flow over one day |
| | total flow amount ($amount^{name}_{type}$) | $\sum\{amount^m_{type}, .., amount^n_{type}\}, \{m, n\} \in \mathbb{N}$ | ind. m$^{-2}$ | Summed amount (per cohort) of density flow over one day |
| Climate Cell ($\Omega$) | ID | $\in [1,107]$ | | Unique identifier of Climate Cells |
| | $center_{x,y}$ | $x, y \in \mathbb{N}$ | | Geometric center in coordinate system of Grassland Cells |
| | temperature ($\omega_{ts}$) | | °C | Local surface temperature |
| | humidity ($\omega_{rhug}$) | | % | Local relative humidity in the upper 2 cm of the ground |
| | contact water ($\omega_{cw}$) | | kg m$^{-2}$ | Amount of water in the upper 2 cm of the ground |
| Grassland Cell ($G$) | coordinate ($coord_{x,y}$) | $x, y \in \mathbb{N}$ | | Index in Cartesian coordinate system |
| | carrying capacity ($cap_{above}$) | 25 | ind. m$^{-2}$ | Maximum aboveground density |
| | climate value(s) ($\omega_{clim}$) | | | Calculated by weighing resp. values of adjacent Climate Cells |
| | mowing schedule ($T_{mow}$) | See Table S1-3 | days | Occurrence days of mowing events |

**Abbreviations: above=aboveground, C=Cohort, cap=capacity, coord=coordinate, cw=contact water, dens=density, dest=destination, dev=development, dia=diapause, disp=dispersal, dyn=dynamic, emb=embryo, F=Flow, G=Grassland Cell, hab=habitat, ID=identifier, ima=imago, ind=individuals, kg=kilogram, lar=larva, m=meter, max=maximum, min=minimum, mort=mortality, mow=mowing, orig=origin, P=Population, pre=pre-diapause, prog=progress, repr=reproduction, rhug=relative humidity upper ground, S=Life Stage, t=time step, ts=surface temperature, trans=transfer**

M1: The auxiliary entity *Flow* was introduced to ease the implementation of the model. Each *Flow* is characterized by the *life stage of origin* and *life stage of destination* (empty for mortality flow), a static per capita *base flow rate* [day$^{-1}$] and a *total flow amount* [individuals m$^{-2}$]. For each *Cohort* in its *life stage of origin* it has a per capita *dynamic flow rate* [day$^{-1}$] and a current *flow amount* [individuals m$^{-2}$]. These *flow rates* are calculated using external *Influences* (section 7.2). The different *types* of *Flows* define the different flow processes: *trans* (transferring density to a subsequent stage: 1->2, 2->3, 3->4, 4->5); *repr* (reproduction from imago (no loss) to pre-diapause stage; 5->1). Additionally, all five *Life Stages* and their *Cohorts* lose density through mortality (*Flow type mort*).

M2: A fourth *Flow type disp* (*dispersal*) is introduced defining the density transfer from a *Life Stage* to the same *Life Stage* (here, imago only) of a neighboring *Population*. Potential density loss during dispersal of the imago *Life Stage* is handled by an additional mortality *Flow*.

M1: The *Grid Cells* comprising the environment are characterized by their coordinate (cell indexes), *carrying capacity* (maximum number of aboveground population that can be sustained), daily climate conditions (*temperature*, *humidity*, *contact water*[3]) and land use schedule (*mowing day*).

M2: *Grid Cells* are replaced by the two entities *Climate Cell* and *Grassland Cell* with the indexes of the former and the geometric center of the latter belonging to the same Cartesian coordinate systems. In this coordinate system, x-coordinates increase from West to East and y-coordinates from North to South. *Climate Cells* have a unique ID and contain the daily climate conditions in the original spatial resolution. *Grassland Cells* take the *carrying capacity* and a *mowing schedule* while adapting the climate conditions of up to four adjacent *Climate Cells* to calculate the local climate conditions using bilinear interpolation. Both the *mowing schedules* and the bilinear interpolation will be described in more detail below.

M1: The model uses daily time steps for updating the model's states and process. This time scale also reflects the sampling of the climate data. However, the single possible mowing event per year is considered on a weekly basis. To account for this weekly frequency, a year has 364 days by definition, resulting in exactly 52 full calendar weeks year$^{-1}$. Input data (i.e., climate time series) is either cropped or expanded accordingly. Simulations were run for 20 years (7280 time steps) or stopped earlier in case all *Cohorts* of the *Population* became extinct.

M2: For each year of the climate data, February 29$^{th}$ (if exists) and December 31$^{st}$ are omitted to achieve 364 days. A simulation run takes 21840 time steps (60 years) starting on January 1st 2020

[3] i.e., such that LMG eggs are covered with water or lie in moist soil (Ingrisch, 1983)

and ending on December 30th 2079. In the case of premature extinction of all Populations, simulations stop earlier.

M1: The environment, or study region, comprises 1296 (36 x 36) cells, each having an area of 144 $km^2$ (12 km x 12 km), which corresponds to the resolution of the climate input data (section 6). The integer grid cell indexes (1-36) are mapped to rotated pole grid coordinates (main manuscript, Figure 1). 968 grid cells are terrestrial and therefore belong to the model domain. Within a grid cell a single habitat is simulated and represents a squared virtual grassland plot with the size of 6.25 ha (250 x 250 $m^2$). Grid cells and hence habitats are not connected, i.e., there is no exchange of individuals. If populations become extinct, there is no recolonization.

M2: The study region is the German federal state of Schleswig-Holstein consisting of a 107 terrestrial *Climate Cell* subset of the original data and 72,969 *Grassland Cells* representing the state's actual grassland area that were retrieved using the software DSS-Ecopay (Mewes et al., 2012; Sturm et al., 2018). Each *Grassland Cell* by definition has a size of 6.25 ha (250 x 250 $m^2$) that reflects the spatial resolution of the data quite well. *Grassland Cells* within a radius of 1,500 m are connected by the dispersal process of the imago *Life Stage*. In some cases, there are additional connections outside this radius representing long distance dispersal (LDD). The connections and dispersal processes will be described below.

### 3 Process overview and scheduling

M1: In each time step and for every grid cell (M2: *inhabited Grassland Cell*), four main blocks of PROCESSES are executed: 'Update environmental drivers', 'Flow update', 'Life Stage update', and 'Cohort update'. The first three blocks are SCHEDULED one after the other, while 'Cohort update' is executed as a submodel of 'Life Stage update'. Pseudocode S1-1 gives an overview of this top level scheduling. During 'Flow update' the *flow rates* and *amounts* are calculated using the *Flow*'s *life stage of origin* while depending on its *type* and specified external influences (submodel 'Update environmental drivers').

M2: The PROCESS 'Bilinear climate interpolation' is executed each time step as a submodel of 'Update environmental drivers'. If the *life stage of destination* of a non-zero (in terms of *flow amount) dispersal Flow* belongs to the *Population* of an *uninhabited Grassland Cell*, the *Population* is not empty anymore and the cell thus rendered *inhabited*. In this case, the submodel 'Dispersal setup' is executed to find all *Grassland Cells* considered neighbors of the *Population* and establish a dispersal connection to each of them by creating a respective *Flow* of type *disp*.

M1: The 'Life Stage update' first handles creation (input from *Flows*) and lastly deletion (density falling below *minimum density*) of its *Cohorts*. In between *Cohort* creation and deletion, the submodel 'Cohort update' is executed.

**Pseudocode S1-1: Main process overview. Processes executed at each inhabited Grassland Cell and for each time step. The processes 'Bilinear climate interpolation' and 'Cohort Update' are executed as submodel of 'Update environmental drivers' and 'Life Stage update', respectively.**

```
FOR each inhabited Grassland Cell / Population
   EXECUTE submodel 'Update environmental drivers'
   FOR each originating Flow
      EXECUTE submodel 'Flow update'
      IF (Flow type = DISPERSAL
          AND Flow amount > 0
          AND target Population density = 0)
      THEN
         DEFINE target Grassland Cell as inhabited
      ENDIF
   ENDFOR
ENDFOR
FOR each inhabited Grassland Cell / Population
   SET Population density := 0
   FOR each Life Stage
      EXECUTE submodel 'Life Stage update'
      ADD Life Stage density to Population density
   ENDFOR
   IF Population density < minimum density THEN
      DEFINE Grassland Cell as uninhabited
   ENDIF
ENDFOR
```

### 4 Design concepts

*Basic Principles:*

M1: The model uses viability, i.e., the ability of small populations to persist, as a comparative metric to identify grassland mowing schedules that are compatible with the projected climate conditions in a certain region. Development and mortality of the different life stages are driven by environmental factors. For this, relationships were imposed that reflect the available empirical knowledge and data.

M2: The extension uses the same comparative metrics and environmental drivers as the original study. Additionally, it applies density-independent dispersal using a fat-tailed dispersal kernel that changes depending on regional grassland cover. Relevant dispersal parameters were adapted from empirical studies of the target species.

*Emergence:*

M1: The relationships describing the life cycle are imposed and not emergent from first principles, such as energy budgets or adaptive decision making.

M2: The dispersal metrics are imposed by the parameter definitions such as dispersal radius.

*Interaction:*

M1: The model does not include direct interaction within or among different life stages. Indirect interactions are included by assuming density dependence of the mortality of larvae and imagines.

M2: Neighboring *Populations* of inhabited *Grassland Cells* indirectly interact through a dispersal process that depends on the grassland cover surrounding the originating cell.

*Stochasticity:*

M1: Transfer from embryo to larval ($F_{trans}^{emb}$, Table S1-1) and larval to imago ($F_{trans}^{lar}$) *Life Stage* is stochastically drawn from a binominal distribution (section 7.1). The probability is influenced by *temperature*, *Cohort density* and *development progress*.

M2: Dispersal mortality as well as the actual dispersal from a *Population's* imago *Life Stage* to the imago *Life Stages* of its connected *Grassland Cells* is determined stochastically using a density dependent binomial distribution. The dispersal probability for each connection is calculated using a predefined base dispersal rate, a distance dependent dispersal preference, a probability of finding the connected neighbor and a survival probability, where the latter two depend on grassland cover and distance between both *Grassland Cells*. The probability of dispersal mortality is derived from the summed dispersal probabilities.

### 5 Initialization

M1: The model is initialized with a *starting date*, *duration*, *mowing day* and *climate change scenario* (CCS). Additionally, each *Population* receives an initial density per *Life Stage*. The *carrying capacity* [individuals m$^{-2}$] for the aboveground population is assumed to be the same for all *Grid Cells* to reduce the number of confounding factors in the model analysis. As the climate data only offers a single projection per RCP scenario (cf. main manuscript, section 2.3) and location, we resampled the time series per replicate run (seed): for a simulation period of 20 years, we used a time frame of 30 years (simulation period ±5 years) and reordered the years by sampling with replacement. This resampling is feasible since there are no significant trends in the relevant climate parameters within such a period (K. Keuler, *pers. comm.)*. Simulation period 2060-79 has just 26 years, because climate data was only available up to the year 2080. A list of the sampled years per seed and simulation period is supplied in Supplement S2. The initial settings of the model are listed in Table S1-2.

**Table S1-2: List of parameters used to initialize a simulation run. First column: parameter name used in text. Second column: parameter symbol when used in model equations. Third column: initial value(s) resp. value options used for simulation runs. Fourth column: brief description of parameter. Parameters below double line can be varied in principle, but are constant in the presented work. M2: last five rows contain relevant parameters for dispersal process**

| Parameter Name | Symbol | Value(s) | Description |
|---|---|---|---|
| starting date | $t_{init}$ | 01 January 2020 | The date of initial time steps translated to climate data index |
| mowing day | $t_{mow}$ | none or day 134, 141,..,274 | The timing of mowing per year |
| mowing schedule | $T_{mow}$ | | A set of mowing days per year (see Table S1-3) |
| Climate change scenario | $CCS$ | FF, MOD or BAU | Representative Concentration Pathways of $CO_2$ model |
| duration | $t_{\Delta}$ | 21,840 days | Runtime in days resp. time steps |
| climate cell size | $size_{clim}$ | 12,000 m | Width and height of square climate cells |
| habitat size | $size_{hab}$ | 250 m | Width and height of square habitats / grassland cells |
| habitat area | $A_{hab}$ | 250 x 250 m$^2$ | Area of grassland plot inside a grid cell |
| initial density | $dens_{init}$ | {0,0.725,0,0,0} ind. m$^{-2}$ | The initial population density per life stage in individuals m$^{-2}$ |
| carrying capacity | $cap_{above}$ | 25 individuals m$^{-2}$ | Maximum aboveground density per square meter |
| dispersal radius | $rad_{disp}$ | 1,500 m | Maximum distance covered by an individual (Griffioen, 1996) |
| base dispersal rate | $rate_{disp}^{ima}$ | 0.00595 day$^{-1}$ | Daily rate of furthest dispersing imagos (Malkus, 1997) |
| dispersal preference | $pref^{near}$ | 1 | Preference of selecting a neighbor during the dispersal process. Higher values result in selection of closer neighbors. |
| sight | $sight_{disp}$ | 0.5 | Ability to find a selected neighbor during a dispersal process |
| decay rate | $dec_{disp}$ | 0.04 | Distance dependent probability of surviving dispersal |

**(FF=full force, MOD=moderate, BAU, business as usual, CCS=climate change scenario, above=aboveground, cap=capacity, clim=climate, dec=decay, dens=density, disp=dispersal, hab=habitat, ima=imago, init=initial, mow=mowing, pref=preference, rad=radius, scen=scenario, t=time step)**

M2: Every simulation starts on 01 January 2020 and runs for 60 years (21,840 time steps). A run is initialized with one of three CCS, a single starting *Population* in the *Grassland Cell* closest to the geometric center of either one of the 107 *Climate Cells* and one out of 18 mowing schedules (Table S1-3). Ignoring replicates, this results in a total of 5,778 distinct simulations runs. In the *Grassland Cell* of the starting *Population* the base mowing schedule named M20+00+44 always applies. Here, the first number of the schedule's name stands for early mowing calendar week 20 (day 133) and the last number for late mowing week 44 (day 301). The middle number defines the (additional) mowing weeks 22-38 of more intensive grassland management schedules (acronyms: M22-M38). All other cells receive the initially defined schedule. In that way, the starting location works as a rather undisturbed habitat of low-impact grassland mowing rendering it a fixed point for analyzing the dispersal process. Early mowing (day 133) for schedules M22-25 is omitted, because cuts should be at least six weeks apart, and late mowing (day 301) for schedules M35-38, because it us unnecessary due to slowed grassland growth and being economically unbeneficial for farmers (Gerling et al., 2022). Per replicate run, each simulation year was resampled randomly using ±10 years, e.g. the simulation year 2053 was determined using the set $[2043,2044,..2063]$. If the set would exceed the available data (e.g. for years > 2080), it is reduced accordingly.

**Table S1-3: Yearly grassland mowing schedules as applied in the simulation runs. First column gives the names of the 18 mowing schedules that encode the calendar weeks of yearly mowing occurrence divided by a plus (+) symbol. An acronym of the schedule name is provided in the second column, encoding the relevant mowing week in its name. The last three columns give the actual yearly mowing days (first day of respective calendar week) per mowing schedule. Schedules that include cells containing a dash (encoded by '00' in the respective name) only have two mowing occurrences per year, all others have three. The first mowing schedule M20+00+44 represents low-impact mowing, while more intensive mowing schedules follow in the rows below the double line.**

| Schedule name | Acronym | Mowing days | | |
|---|---|---|---|---|
| *M20+00+44* | *M00* | 133 | - | 301 |
| *M00+22+44* | *M22* | - | 147 | 301 |
| *M00+23+44* | *M23* | - | 154 | 301 |
| *M00+24+44* | *M24* | - | 161 | 301 |
| *M00+25+44* | *M25* | - | 168 | 301 |
| *M20+26+44* | *M26* | 133 | 175 | 301 |
| *M20+27+44* | *M27* | 133 | 182 | 301 |
| *M20+28+44* | *M28* | 133 | 189 | 301 |
| *M20+29+44* | *M29* | 133 | 196 | 301 |
| *M20+30+44* | *M30* | 133 | 203 | 301 |
| *M20+31+44* | *M31* | 133 | 210 | 301 |
| *M20+32+44* | *M32* | 133 | 217 | 301 |
| *M20+33+44* | *M33* | 133 | 224 | 301 |
| *M20+34+44* | *M34* | 133 | 231 | 301 |
| *M20+35+00* | *M35* | 133 | 238 | - |
| *M20+36+00* | *M36* | 133 | 245 | - |
| *M20+37+00* | *M37* | 133 | 252 | - |
| *M20+38+00* | *M38* | 133 | 259 | - |

### 6 Input data

M1: As input source, time series of climate data for each grid cell are used. Each climate parameter (*ts, pr, mrso, smt* – cf. main manuscript, section 2.3 and Supplement S1) is read from a single NetCDF data file that stores one data point per day and coordinate. Files provided for this work contain time series from 01 January 1995 to 31 December 2080 including leap years. All data points on 29 February and 31 December are removed to achieve 364 day long years (section 2).

M2: Climate parameters of a *Grassland Cell* are determined using up to four terrestrial *Climate Cells* in direct squared neighborhood that in terms of their geometrical center are closest to the coordinate of the *Grassland Cell*. The parameter values are then calculated using bilinear interpolation (Section 7.6) resulting in a two-dimensionally gradual climate data resampling of higher spatial resolution (107 to 72,969 spatial data points).

### 7 Submodels

M1: Table S1-4 gives an overview of the model processes and dynamics rates. The underlying full equations are provided in Table S1-5 and the corresponding parameters in Table S1-6.

M2: Sections 7.1.7, 7.5 and 7.6 describe the dispersal process between *Populations*, the setup of a dispersal network within a neighborhood of *Grassland Cells* and the calculation of climate values for a *Grassland Cell* using bilinear interpolation of values stemming from coarse-scale parameters of adjacent *Climate Cells*.

**Table S1-4: Overview of the model processes and their daily rates and equations. Processes (second column) are distinguished by *Life Stage* (first column) and referenced by their symbol (third column) as used in the model equations. The fourth column defines the equation and environmental drivers (*Influences*, f-symbols, cf. Table S1-5) used for calculating a cohort-specific dynamic rate. Equation segments marked with *s are simplifications of the iterative process of updating the flow rate described in Pseudocode S1-4. Superscript letters reference the sources used to parameterize the processes and equations for the LMG (coefficients in Table S1-6): [a] Ingrisch (1983), [b] B. Schulz (pers. comm.), [c] Wingerden et al. (1991), [d] Ingrisch and Köhler (1998), [e] Helfert and Sänger (1975), [f] Helfert (1980), [g] Kriegbaum (1988), [h] Waloff (1950), [i] Griffioen (1996), [j] Malkus (1997).**

| Life stage (symbol) | Process | Process symbol | Dynamic rate (daily) | Description |
|---|---|---|---|---|
| pre-diapause ($S^{pre}$) | mortality | $F^{pre}_{mort}$ | $dyn^{ID}_{mort} = rate^{pre}_{mort} + (1 - rate^{pre}_{mort})^* \times [f^A_{sig} \times f^B_{thd} + f^C_{mow}]$ | The base mortality rate [a] increases in two cases: (1) humidity-driven ($f^A_{sig}$) [a] if contact water ($f^B_{thd}$) [a] is missing; (2) if mowing is scheduled ($f^C_{mow}$) [b] |
| | development | $prog^{pre}_{ID}$ | $prog^{pre}_{ID} = \begin{cases} prog^{pre}_{ID} + f^D_{thd} & if f^D_{thd} > 0 \\ 0 & otherwise \end{cases}$ | Needs three days below 10 °C in a row ($f^D_{thd}$) [a] to fully develop |
| | transfer | $F^{pre}_{trans}$ | $dyn^{ID}_{trans} = \begin{cases} 1 & if prog^{pre}_{ID} = 1 \\ 0 & otherwise \end{cases}$ | Immediately transfers to diapause stage if fully developed |
| diapause ($S^{dia}$) | mortality | $F^{dia}_{mort}$ | $dyn^{ID}_{mort} = rate^{dia}_{mort} + \left(1 - rate^{dia}_{mort}\right) \times f^E_{mow}$ | The base mortality rate increases if mowing is scheduled ($f^E_{mow}$) [b] |
| | development | $prog^{dia}_{ID}$ | $prog^{dia}_{ID} = \begin{cases} prog^{dia}_{ID} + f^F_{thd}/2 & if prog^{dia}_{ID} < 0.5 \\ prog^{dia}_{ID} + f^G_{thd}/2 & if prog^{dia}_{ID} \geq 0.5 \wedge f^G_{thd} \neq 0 \\ 0.5 & otherwise \end{cases}$ | Needs 61 days below 5 °C to break diapause ($f^F_{thd}$) [a,c] and afterwards three days above 10 °C in a row ($f^G_{thd}$) [a] to fully develop. Temperatures > 5 °C before diapause is broken reverse development |
| | transfer | $F^{pre}_{trans}$ | $dyn^{ID}_{trans} = \begin{cases} 1 & if prog^{dia}_{ID} = 1 \\ 0 & otherwise \end{cases}$ | Immediately transfers to embryo stage if fully developed |
| embryo ($S^{emb}$) | mortality | $F^{emb}_{mort}$ | $dyn^{ID}_{mort} = rate^{emb}_{mort} + (1 - rate^{emb}_{mort})^* \times [f^H_{exp} + f^I_{sig} \times f^J_{thd} + f^K_{mow}]$ | The base mortality is defined by the temperature ($f^H_{exp}$) [c]. It increases in two cases: (1) humidity-driven ($f^I_{sig}$) [a] if contact water ($f^J_{thd}$) [a] is missing and/or (2) if mowing is scheduled ($f^K_{mow}$) [b] |
| | transfer | $F^{emb}_{trans}$ | $dyn^{ID}_{trans} = rate^{emb}_{trans} \times f^L_{bin}\left(f^M_{exp}, f^N_{exp}, dens^{ID}, prog^{name}_{ID}\right)$ | The base transfer rate is multiplied by a temperature- ($f^M_{exp}$, $f^N_{exp}$) [c], progress- and the density-driven hatching probability stemming from a binomial distribution ($f^L_{bin}$) |
| larva ($S^{lar}$) | mortality | $F^{lar}_{mort}$ | $dyn^{ID}_{mort} = rate^{lar}_{mort} + (1 - rate^{lar}_{mort})^* \times [f^O_{cap} + f^P_{lin} + f^Q_{mow}]$ | The base mortality rate [d] increases with density ($f^O_{cap}$) [b] and in two additional cases: (1) the temperature ($f^P_{lin}$) is below 10 °C and/or (2) if mowing is scheduled ($f^Q_{mow}$) [b] |
| | transfer | $F^{lar}_{trans}$ | $dyn^{ID}_{trans} = rate^{lar}_{trans} \times f^R_{bin}(f^S_{lin}, f^T_{lin}, dens^{ID}, prog^{lar}_{ID})$ | The base transfer rate is multiplied by a temperature- ($f^S_{lin}$, $f^T_{lin}$) [e,f], progress- and density-driven maturation probability stemming from a binomial distribution ($f^R_{bin}$) |
| imago ($S^{ima}$) | mortality | $F^{ima}_{mort}$ | $dyn^{ID}_{mort} = rate^{ima}_{mort} + \left(1 - rate^{ima}_{mort}\right)^* \times [f^U_{cap} + f^V_{lin} + f^W_{mow} + f^X_{bif}]$ | The base mortality rate [g] increases with density ($f^U_{cap}$) [b] and in three additional cases: (1) the temperature ($f^V_{lin}$) is below 10 °C, (2) if mowing is scheduled ($f^W_{mow}$) [b] and/or (3) loss during dispersal ($f^X_{bif}$) |
| | reproduction | $F^{ima}_{repr}$ | $dyn^{ID}_{repr} = rate^{ima}_{repr}$ | Fixed reproduction rate [d,h] |
| | dispersal | $F^{ima}_{disp}(a, b)$ | $dyn^{ID}_{disp} = f^Y_{bif}$ | Dispersal rate from a to b depends on grassland cover around a, and is stochastically determined by a binomial distribution ($f^Y_{bif}$) [i,j] |

**Abbreviations: bif=binomial factor, bin=binominal climate, cap=capacity, dens=density, dia=diapause, disp=dispersal, emb=embryo, exp=exponential, f=symbol of Influence function, F=Flow, ID=Cohort identifier, ima=imago, lar=larva, lin=linear, mort=mortality, mow=mowing, pre=pre-diapause, prog=progress, repr=reproduction, sig=sigmoid, t=time step, thd=threshold, trans=transfer**

#### *7.1 Update environmental drivers*

M2: The environmental drivers are updated every time step for each *Grassland Cell*. This means here that the climate values are recalculated by bilinear interpolation (Section 7.6) and it is checked whether a mowing schedule takes effect.

M1: Both *flow rates* and a *Cohort*'s *development progress* can change depending on the current value of environmental drivers or static conditions. Our model includes functions or equations – called *Influences* – that may be applied to achieve dynamic changes.

Generally speaking, each *Influence* provides a *factor* that can mediate the effect of environmental conditions on the variables *dynamic flow rate* and *development progress* of a *Flow* or *Cohort*, respectively. This *factor* may be restricted to a *minimum* and *maximum* value ($f_{min} = 0$ and $f_{max} = 1$, if not specified otherwise) regardless of those conditions and contributes either in a *multiplicative* or *additive* way to the update of a rate or progress. The contribution to an *Influences*' current *factor* $f_{inf}$ by a *multiplicative factor* $f_{mult}$ is – as the name suggests – simply $f_{inf} \times f_{mult}$; while an *additive factor* $f_{add}$ contributes via the equation

$$f_{inf} + (1 - f_{inf}) \times f_{add} \tag{S1-1}$$

A *factor* is calculated during the update of the associated *Flow* / *Cohort*. The specific *Influences* are described below and their equations summarized in Table S1-5. *Influences* may be combined for the use in a process. The empirical basis for these equations is given below.

**Table S1-5: List of equations / functions applied for environmental drivers including numbering (first column), name as used in the text (second), symbol with subscript of name abbreviation (third) and brief description (last)**

| Eqn# | Name | Equation / Function (symbol) | Description |
|---|---|---|---|
| S1-2 | Capacity Influence | $\mathbf{f_{cap}} = \left(1.0 + e^{\beta^1_{cap}\left(\frac{dens^P_{above}}{A_{hab}} - \beta^2_{cap} cap_{above}\right)}\right)^{-1}$ | Applying carrying capacity for the aboveground population |
| S1-3 | Linear Climate Influence | $\mathbf{f_{lin}} = \beta^1_{lin} + \beta^2_{lin}\omega_{clim}$ | Linear correlation with a climate parameter |
| S1-4 | Exponential Climate Influence | $\mathbf{f_{exp}} = \beta^1_{exp} + \beta^2_{exp} e^{\beta^3_{exp}\omega_{clim}}$ | Exponential correlation with a climate parameter |
| S1-5 | Sigmoid Climate Influence | $\mathbf{f_{sig}} = \beta^1_{sig}\left\lvert 1 - \left(1 + e^{-\beta^2_{sig}\left[\omega_{clim} - \beta^3_{sig}\right]}\right)^{-1}\right\rvert$ | Sigmoid correlation with a climate parameter |
| S1-6 | Factor Threshold | $\mathbf{f_{thd}} = \begin{cases} \beta^1_{thd} & \text{if } \omega_{clim} \leq thd_{clim} \\ \beta^2_{thd} & \text{otherwise} \end{cases}$ | Varying factor depending on threshold exceedance |
| S1-7 | Binominal Climate | $\mathbf{f_{bin}}(\mu_{\Delta t}, \sigma_{\Delta t}, dens^{ID}, prog^{name}_{ID})$ | This factor is a climate-, density- and progress-driven probability stemming from a binomial distribution. See Pseudocode S1-2 for details |
| S1-8 | Land Use Influence | $\mathbf{f_{mow}} = \begin{cases} mort_{mow} & \text{if } t_{mow} = t \\ 0 & \text{otherwise} \end{cases}$ | Increased mortality if mowing occurs |
| S1-9 | Binominal Factor | $\mathbf{f_{bif}}(\beta^1_{bif}, dens^{ID})$ | Density-driven probability stemming from a binomial distribution of a predefined mean (factor). See Pseudocode S1-3 for details |

**(above=aboveground, bif=binominal factor, bin=binominal climate, cap=capacity, clim=climate, dens=density, exp=exponential, hab=habitat, ID=Cohort identifier, lin=linear, mort=mortality, mow=mowing, prog=progress, sig=sigmoid, t=time step, thd=treshold)**

M1: The *Capacity Influence* is used in a logistic function that increases its *factor* with increasing population density. It restricts life stages to a maximum size by raising the mortality for large populations (Eqn. S1-2).

A *Linear Climate Influence* provides a *factor* that is linearly correlated with a specified climate value either in a positive or negative manner (Eqn. S1-3).

Similar to the linear correlation an *Exponential Climate Influence* can be used to provide a *factor* that is exponentially correlated to a specified climate value (Eqn. S1-4).

The *Sigmoid Climate Influence* is the third option to provide a *factor* by correlating with a climate value (Eqn. S1-5). It allows bounding a *factor* to an upper and lower limit without clipping it.

A *Factor Threshold* is used with either of the above *Influences*. It allows applying two different values depending on whether a defined climate threshold is exceeded (Eqn. S1-6). It can be used, for instance, to either enable another *Influence* (threshold exceeded, *factor* = 1) or disable it (threshold not exceeded, *factor* = 0). An iterative use of this *Influence* is possible to apply thresholds for multiple climate values.

The *Binomial Climate* (M2: renamed from *Factor*) is an Influence that depends on climate conditions and a *Cohort*'s *density* and *development progress*. It stochastically determines its *factor* by drawing from a binomial distribution (Eqn. S1-7, Pseudocode S1-2). In other words, it defines which share of a *Cohort* population (density) is affected by an event (e.g. death). The binomial distribution is approximated by the cumulative distribution of a standard normal distribution. Their mean and standard deviation are calculated using one of the above *Climate Influences*. In that way, population statistics stemming from climate-driven stochastic processes can be translated to a dynamic *factor* like mortality rate.

**Pseudocode S1-2: 'Binominal Climate' (M2: renamed from Factor) function to calculate a flow rate using a stochastically determined flow amount taken from a binominal distribution that is emulated by a normal distribution. The flow probability used to calculate the flow amount is taken from cumulative normal standard deviations using climate-dependent mean and standard deviation of flow duration. The resulting absolute flow rate is the ratio from flow amount to Cohort density. Finally, the applied relative flow rate takes development progress – or rather previously determined flow amount – into account to ensure flow of full density after maximum flow duration / full progress is reached.**

```
BEGIN Influence 'Binominal Climate'
   GET climate-dependent mean flow duration μ_Δt and standard deviation σ_Δt
   GET Cohort's density and (development) progress
   DEFINE cumulative standard normal distribution: CDF(x)
   DEFINE normal distribution: NORM(mean, SD)
   SET climate-dependent maximum flow duration: max_Δt := μ_Δt + 3 * σ_Δt
   SET climate-dependent relative age of Cohort: age := max_Δt * progress
   SET today's age diff to mean duration: diff_tod := (age - μ_Δt) / σ_Δt
   SET tomorrow's age diff to mean duration: diff_tom := (age + 1 - μ_Δt) / σ_Δt
   SET flow probability: p_trans := CDF(diff_tom) - CDF(diff_tod)
   SET mean flow amount: μ_amount := density * p_trans
   SET standard deviation of flow amount: σ_amount := SQRT(μ_amount * (1 - p_trans))
   DRAW from normal distribution to determine amount: n_amount := NORM(μ_amount, σ_amount)
   SET absolute flow rate: r_abs := n_amount / density
   SET relative flow rate: r_rel := 0
   IF progress < 1
      SET r_rel := r_abs / (1 - progress)
   ENDIF
   IF r_rel > 1
      SET r_rel := 1
   ENDIF
   SET progress := progress + r_rel
   RETURN r_rel
END Influence 'Binominal Climate'
```

M1: *Land Use Influence* is a timed event that increases the mortality of a *Cohort* differently depending on its *Life Stage*. In our model, land use is defined as a mowing event that occurs once a year (Eqn. S1-8). M2: The model extension adds the option to provide a mowing schedule that defines a set of days per year on which land use occurs.

M2: The *Binomial Factor* works similarly to the *Binomial Climate Influence* described above but deviates in two aspects. First, mean and standard deviation are calculated using a predefined probability *factor* only. Second, it just uses a *Cohort's density* for the calculation ignoring the *development progress* (Eqn. S1-9, Pseudocode S1-3).

**Pseudocode S1-3: The 'Binominal Factor' function calculates a flow rate using a stochastically determined flow amount taken from a binominal distribution that is emulated by a normal distribution. The predefined probability factor is used directly as mean value of the distribution and indirectly to calculate the standard deviation. The resulting flow rate is the ratio of flow amount compared to Cohort density.**

```
BEGIN Influence 'Binominal Factor'
   GET predefined probability factor p_flow
   GET Cohort's density
   DEFINE normal distribution: NORM(mean, SD)
   SET mean flow amount: μ_amount := density * p_flow
   SET standard deviation of flow amount: σ_amount := SQRT(μ_amount * (1 - p_flow))
   DRAW from normal distribution to determine amount: n_amount := NORM(μ_amount, σ_amount)
   SET flow rate: r_flow := n_amount / density
   IF r_flow > 1
      SET r_flow := 1
   ENDIF
   RETURN r_flow
END Influence 'Binominal Factor'
```

M1: In the following we describe the life cycle processes of the LMG separately for each life stage and define the environmental drivers influencing them. The parametrization for the belowground life stages is mainly adapted from temperature experiments conducted by Wingerden et al. (1991) and soil moisture and contact water experiments by Ingrisch (1983). For the aboveground population, data from different sources was used (sections 7.1.4 and 7.1.5) – not all of them including LMG explicitly. The parameter values for the impact of grassland were established following personal communication with B. Schulz[4] (section 7.1.6).

M2: Information incorporated to estimate and validate the dispersal process (Section 7.1.7) of the LMG are an empirical 'mark and recapture' study by Malkus (1997), field studies by Griffioen (1996) and Marzelli (1994), as well as two experiments using genetic markers to measure distances between populations (Keller et al., 2012; Van Strien, 2013)

M1: The values for base rates and the coefficients used to parameterize the *Influences* as applied for the processes of the target species are listed in Table S1-6. A verbal description on their usage for the target species is given in the following subsections.

[4] Conservation agency "Stiftung Naturschutz Schleswig-Holstein"

**Table S1-6: List of coefficients (third column) used to parameterize the Influences equation (second column, Table S1-5) as used to modify the processes (first column, Table S1-4) of the target species.**

| Process symbol | Influences | Base rate of process and/or coefficients of influences | Sources |
|---|---|---|---|
| $F_{mort}^{pre}$ | $f_{sig}^{A}, f_{thd}^{B}, f_{mow}^{C}$ | $(rate_{mort}^{pre} = 8.164147 \times 10^{-4})$[a], $(\beta_{sig}^{A1} = 0.357, \beta_{sig}^{A2} = 17.183, \beta_{sig}^{A3} = 0.703, \omega_{clim}^{A} = \omega_{rhug})$[a], $(\beta_{thd}^{B1} = 1, \beta_{thd}^{B2} = 0, \omega_{clim}^{B} = \omega_{cw}, thd_{clim}^{B} = 0\ kg\ m^{-2})$[a],$(mort_{mow}^{C} = 0.05)$[b] | [a] Ingrisch (1983)<br>[b] B. Schulz (pers. comm.)<br>[c] Wingerden et al. (1991)<br>[d] Ingrisch & Köhler (1998)<br>[e] Helfter & Sänger (1975)<br>[f] Helfert (1980)<br>[g] Kriegbaum (1988)<br>[h] Waloff (1950)<br>[i] Griffioen (1996)<br>[j] Malkus (1997) |
| $prog_{ID}^{pre}$ | $f_{thd}^{D}$ | $(\beta_{thd}^{D1} = {}^{1}/_{3}, \beta_{thd}^{D2} = 0, \omega_{clim}^{D} = \omega_{ts}, thd_{clim}^{D} = 10°C)$[a] | |
| $F_{mort}^{dia}$ | $f_{mow}^{E}$ | $(rate_{mort}^{pre} = 8.164147 \times 10^{-4})$[a], $(mort_{mow}^{E} = 0.05)$[b] | |
| $prog_{ID}^{dia}$ | $f_{thd}^{F}, f_{thd}^{G}$ | $(\beta_{thd}^{F1} = {}^{1}/_{61}, \beta_{thd}^{F2} = -{}^{1}/_{182}, \omega_{clim}^{F} = \omega_{ts}, thd_{clim}^{F} = 5°C)$[a,c], $(\beta_{thd}^{G1} = 0, \beta_{thd}^{G2} = {}^{1}/_{3}, \omega_{clim}^{G} = \omega_{ts}, thd_{clim}^{G} = 10°C)$[a] | |
| $F_{mort}^{emb}$ | $f_{exp}^{H}, f_{sig}^{I}, f_{thd}^{J}, f_{mow}^{K}$ | $(\beta_{exp}^{H1} = 6.949 \times 10^{-3}, \beta_{exp}^{H2} = 5.45 \times 10^{-8}, \beta_{exp}^{H3} = 0.4743, \omega_{clim}^{H} = \omega_{ts})$[c], $(\beta_{sig}^{I1} = 0.351, \beta_{sig}^{I2} = 40, \beta_{sig}^{I3} = 0.975, \omega_{clim}^{I} = \omega_{rhug})$[a], $(\beta_{thd}^{J1} = 1, \beta_{thd}^{J2} = 0, \omega_{clim}^{J} = \omega_{cw}, thd_{clim}^{J} = 0\ kg\ m^{-2})$[c], $(mort_{mow}^{K} = 0.05)$[b] | |
| $F_{trans}^{emb}$ | $f_{exp}^{M}, f_{exp}^{N}$ | $(\beta_{exp}^{M1} = 7.991, \beta_{exp}^{M2} = 1069.98, \beta_{exp}^{M3} = -0.2248, \omega_{clim}^{M} = \omega_{ts})$[a], $(\beta_{exp}^{N1} = 0.9251, \beta_{exp}^{N2} = 126.3933, \beta_{exp}^{N3} = -0.1978, \omega_{clim}^{N} = \omega_{ts})$[a] | |
| $F_{mort}^{lar}$ | $f_{cap}^{O}, f_{lin}^{P}, f_{mow}^{Q}$ | $(rate_{mort}^{lar} = 0.0358)$[d], $(\beta_{cap}^{O1} = -1.5, \beta_{cap}^{O2} = 0.85)$, $(f_{max}^{P} = 0.95, \beta_{lin}^{P1} = 1.9, \beta_{lin}^{P2} = -0.19, \omega_{clim}^{P} = \omega_{ts})$, $(mort_{mow}^{Q} = 0.95)$[b] | |
| $F_{trans}^{lar}$ | $f_{lin}^{S}, f_{lin}^{T}$ | $(\beta_{lin}^{S1} = 83, \beta_{lin}^{S2} = -1.679, \omega_{clim}^{S} = \omega_{ts})$[e,f], $(\beta_{lin}^{T1} = -2.2188, \beta_{exp}^{T2} = 0.2188, \omega_{clim}^{T} = \omega_{ts})$[e,f] | |
| $F_{mort}^{ima}$ | $f_{cap}^{U}, f_{lin}^{V}, f_{mow}^{W}, f_{bif}^{X}$ | $(rate_{mort}^{ima} = 0.0475)$[g], $(\beta_{cap}^{U1} = -1.5, \beta_{cap}^{U2} = 0.85)$, $(f_{max}^{V} = 0.95, \beta_{lin}^{V1} = 1.9, \beta_{lin}^{V2} = -0.19, \omega_{clim}^{V} = \omega_{ts})$, $(mort_{mow}^{W} = 0.95)$[b], $(\beta_{bif}^{X1} = mort_{disp}^{ima}$, see Eqn. S1-16) | |
| $F_{repr}^{ima}$ | | $(rate_{repr}^{ima} = 1.3)$[d,h] | |
| $F_{disp}^{ima}$ | $f_{bif}^{Y}$ | $(\beta_{bif}^{Y1} = rate_{disp}^{ima}(a, b)$, see Eqn. S1-11)[l,j] | |

**(bif=binominal factor, dia=diapause, disp=dispersal, emb=embryo, exp=exponential, f=symbol of Influence function, F=Flow, ID=Cohort identifier, ima=imago, lar=larva, lin=linear, mort=mortality, mow=mowing, pre=pre-diapause, prog=progress, repr=reproduction, sig=sigmoid, thd=threshold, trans=transfer)**

##### 7.1.1 Pre-diapause life stage

M1: The first sub-stage of the belowground population represents the LMG's clutch after oviposition and before diapause. Following the study of Ingrisch (1983), we define that it requires three consecutive days below a temperature of 10 °C to start diapause thus develop into the next life stage (*Factor Threshold,* Eqn. S1-6). The intrinsic mortality rate rapidly increases if the eggs experience pre-winter drought stress caused by missing contact water (Ingrisch, 1983) (*Factor Threshold,* Eqn. S1-6). The increased mortality is calculated using the *Sigmoid Climate Influence* (Eqn. S1-5) driven by humidity.

##### 7.1.2 Diapause life stage

M1: The diapause life stage occurs mainly during winter and is mostly unaffected by climate conditions in terms of our model. Therefore, the intrinsic mortality rate remains constant. We defined that it needs to experience a total of 61 days below 5 °C to break diapause (*Factor Threshold,* Eqn. S1-6). By that, we followed the cold treatments in the studies by Ingrisch (1983) and Wingerden et al. (1991), though with a shorter period of time to avoid stagnation in slightly warmer winters. Additionally, a too early indication of spring is prevented by defining that the life stage requires three consecutive days of at least 10 °C to develop into the next life stage (*Factor Threshold,* Eqn. S1-6). The diapause life stage can have multiple *Cohorts*, i.e., one for each consecutive year of unsuitable conditions (diapause not broken).

##### 7.1.3 Embryo life stage

M1: This life stage is the most complex in our model. Mortality is influenced by three climate parameters while the temperature-driven hatching process (transfer to larval stage) is stochastically determined by the stage's density and development progress. Following Wingerden et al. (1991) we established the mortality rate using the fact that a temperature of 22.2 °C yields the highest hatching success rate (~82%) while the rate drops with both lower and higher temperatures (*Exponential Climate Influence,* Eqn. S1-4). At the same time higher temperatures decrease the mean hatching time from ~45 days at 15.0 °C to ~8 days at 37.5 °C (*Exponential Climate Influence,* Eqn. S1-4). In our model, the base mortality rate is in fact mainly determined by temperature. Similar to the pre-diapause life stage it can further increase in the case of post-winter drought stress caused by missing contact water (Ingrisch, 1983) (*Factor Threshold,* Eqn. S1-6). This additional mortality is calculated using the *Sigmoid Climate Influence* (Eqn. S1-5) driven by humidity.

As mentioned above, the timing and amount of hatching eggs is stochastically determined (*Binomial Climate* [M2: renamed from *Factor*]*,* Eqn. S1-7). The hatching probability increases with development progress and higher temperatures. Mean and standard deviation defining the binomial distribution to draw the probability from are factors that are correlated with the temperature (*Exponential Climate Influence,* Eqn. S1-4).

##### 7.1.4 Larva life stage

M1: The larval life stage has multiple *Cohorts*, i.e., one for each hatching day per year of the previous embryo stage. These larva *Cohorts* are processed independently. High temperatures increase the development speed of larvae. We used the development traits found in three related locust species (Helfert, 1980; Helfert and Sänger, 1975) to calculate temperature-driven mean and standard deviation (*Linear Climate Influence,* Eqn. S1-3). Both are then applied to stochastically determine the density- and progress-dependent transfer (or rather development) of a larva *Cohort* to the imago life stage (*Binomial Climate* [M2: renamed from Factor]*,* Eqn. S1-7).

The larva base mortality rate was adopted by averaging the parameters of two related locust species (Ingrisch and Köhler, 1998, Tab. 16). It increases with aboveground population density

(*Capacity Influence,* Eqn. S1-2) allowing a maximum of 25 individuals m$^{-2}$ (B. Schulz, *pers. comm.*) and by definition with temperatures below 10 °C (*Linear Climate Influence,* Eqn. S1-3).

##### 7.1.5 Imago life stage

M1: Similar to the larval stage, the base mortality rate for the imago life stage was adopted using the daily survival rates of four related locust species determined by Kriegbaum (1988). It increases with aboveground population density (*Capacity Influence,* Eqn. S1-2) allowing a maximum of 25 individuals m$^{-2}$ (B. Schulz, *pers. comm.*) and by definition with temperatures below 10 °C (*Linear Climate Influence,* Eqn. S1-3). The daily transfer or rather oviposition rate of 1.3 was defined including several considerations. First of all, an LMG egg pod contains 11-14 eggs (Waloff, 1950) and reproduction is delayed up to two weeks after maturation (Ingrisch and Köhler, 1998). Furthermore, we assumed that half of the LMG population is female and that a female lays up to three egg pods during a lifetime of 60 days. In terms of our model, oviposition is not influenced by any external drivers.

##### 7.1.6 Land use

M1: As mentioned above, land use in our model is a representation of grassland mowing that occurs once per year on the first day of the same calendar week. It affects the belowground population (life stages 1-3) less severely than the aboveground population (4-5) by increasing the stage's mortality rate *additively* by 0.05 (below ground) and 0.95 (above ground), respectively (B. Schulz, *pers. comm.*).

M2: Disturbance through land use occurs on 2-3 days per simulation year depending on initially defined mowing schedule (Table S1-3).

##### 7.1.7 Dispersal

M2: In terms of the model extension, the dispersal rate from the imago *Life Stage* of *Population* $P_a$ to the same stage of each neighboring *Population* $P_n \in N_a$ is stochastically determined (*Binomial Factor,* Eqn. S1-9) every time step using a *base dispersal rate* ($rate_{disp}^{ima} = 0.00595\ day^{-1}$) and a pre-calculated *dispersal probability* (Eqn. S1-11). We determined the *base dispersal rate* by defining those individuals as dispersers that travelled the largest distance during one day (1 out of 168) in a 'mark and recapture' study by Malkus (1997). The dispersal mortality rate is calculated using the inverse of the dispersal rate (Eqn. S1-16). Following the maximum covered distance of LMG individuals described by Griffioen (1996), dispersal is defined to remain within a radius ($rad_{disp}$) of 1,500 m, in principle. In regions of low grassland cover, however, dispersal outside this radius (LDD) can occur to account for the LMG's flight ability (Sörens, 1996). Section 7.5 describes in detail, how the neighborhood (both inside and outside of the dispersal radius) of each *Population* is defined and the values of the dispersal process are determined.

#### 7.2 Flow update

During 'Flow update', the state variables *total flow amount, current flow amount* and *dynamic flow rate* (Table S1-4) are iteratively recalculated using the *Cohorts* associated with the *life stage of origin* (Pseudocode S1-4). If there are any external *Influences* (section 7.1) associated with this *Flow*, they change the *dynamic flow rate* depending on their *type*.

**Pseudocode S1-4: Submodel 'Flow Update'. All associated Flows of the defined Population are iteratively updated by this submodel. Sequence of update is irrelevant. Pseudocode describes the update of a single Flow. Equations of influence functions are listed in Table S1-5.**

```
BEGIN submodel 'Flow update'
   SET total flow amount and amount / rate per Cohort to 0.0
   FOR each Cohort in life stage of origin
      IF (Flow type = MORTALITY
         OR Cohort's development progress ≥ 1.0)
      THEN
         CALCULATE age of Cohort from creation time
         IF Flow type = MORTALITY AND age ≥ Life Stage's maximum age THEN
            SET Cohort's flow rate to 1.0
         ELSE
            SET Cohort's flow rate to Flow's base rate
            FOR each Influence in Flow's Influences
               CALCULATE factor by EXECUTING associated Influence function
               CASE OF Influence type
                  ADDITIVE: rate := rate + factor * (1.0 - rate)
                  MULTIPLICATIVE: rate := rate * factor
               ENDCASE
            ENDFOR
         ENDIF
      ENDIF
      SET Cohort's flow amount := rate * density
      SET total flow amount := total flow amount + Cohort's flow amount
   ENDFOR
END submodel 'Flow update'
```

#### 7.3 Life Stage update

The submodel 'Life stage update' (Pseudocode S1-5) calculates a *Life Stage*'s total *density* from the *density* of its subordinate *Cohorts* and its *gain* from the incoming transfer *Flow* of its preceding *Life Stage*. The *gain* is then used to determine whether to create a new *Cohort* from it or add it to an existing *Cohort*. Furthermore, overaged *Cohorts* or such of *density* below the minimum are erased.

**Pseudocode S1-5: Submodel 'Life Stage update'. All associated Life Stages of a Population are iteratively updated using this submodel. The pseudocode describes the update of a single Life Stage.**

```
BEGIN submodel 'Life Stage update'
   SET total density and gain to 0.0
   SET Cohort creation flag := FALSE
   FOR each Flow in input Flows
      ADD Flow's total amount to gain
      IF gain ≥ Life Stage's minimum density THEN
         IF multi cohort Life Stage OR list of Cohorts empty THEN
            CREATE a new Cohort and DEFINE as gaining Cohort
            SET Cohort creation flag := TRUE
            ADD gain to new Cohort's density
         ELSE
            ADD gain to most recent Cohort's density
         ENDIF
      ENDIF
   ENDFOR
   FOR each Cohort in list of Cohorts
      EXECUTE submodule 'Cohort update'
      IF Cohort density < Life Stage's minimum density
         REMOVE Cohort from list of Cohorts
      ELSE
         ADD Cohort's density to total density
      ENDIF
   ENDFOR
   IF Cohort creation flag = TRUE
      ADD new Cohort to list of Cohorts
   ENDIF
END submodel 'Life Stage update'
```

#### *7.4 Cohort update*

During 'Cohort update', *density* and *development progress* of a *Cohort* a recalculated (Pseudocode S1-6). First, loss of *density* is calculated using the outgoing transfer *Flow* of the *Cohort*'s parent *Life Stage*. Then, *gain* (if any) is added to the *density* and reset to zero afterwards. Finally, the *development progress* is increased using external influences, if there were defined any. Otherwise it is set to a value of 1.

**Pseudocode S1-6: Submodel 'Cohort update'. All associated Cohorts of the defined Life Stage are iteratively updated by this submodel. Sequence of update is irrelevant. Pseudocode describes the update of a single Cohort. Equations of influence functions are listed in Table S1-5.**

```
BEGIN submodel 'Cohort update'
   SET loss := 0.0
   FOR each Flow in parent Life Stage's output Flows
      IF Flow type ≠ REPRODUCTION THEN
         DETERMINE Cohort's flow amount from Flow
         ADD Cohort's flow amount to loss
      ENDIF
   ENDFOR
   SET density := density + (gain - loss)
   SET gain := 0.0
   SET temp progress := 1.0
   FOR each Influence in development Influences
      CALCULATE factor by EXECUTING associated Influence function
      SET temp progress := factor * temp progress
   ENDFOR
   ADD temp progress to development progress
   COMPUTE development progress to stay between 0.0 and 1.0
END submodel 'Cohort update'
```

#### 7.5 Dispersal setup

The submodel 'Dispersal setup' is called every time the *life stage of destination* of an empty *Population* is subject in a non-zero *dispersal Flow* (in terms of *flow amount*), in other words, if dispersal is directed to an *uninhabited Grassland Cell* (for the first time). In this case, the submodel locates all *Populations* $P_b \in N_a \subset R$ in the study region $R$ belonging to the sub region or neighborhood $N_a$ of *Population* $P_a$ in the now *inhabited* cell $G_a$ and establishes a connection to each of them by creating a *dispersal Flow* between the imago *Life Stages* of the formerly empty *Population* (*life stage of origin*) and the respective neighboring *Population* (*life stage of destination*). Populations considered a neighbor $P_b \in N_a$ are determined in two ways: (1) all *Populations* inside a predefined dispersal radius (home range); and (2) the nearest *Population* in either one of eight cardinal directions [North (N), Northeast (NE), East (E), Southeast (SE), South (S), Southwest (SW), West (W), Northwest (NW)], in case there are no cells found inside the home range in this direction (LDD). Identifying *Populations* inside the home range is straight forward using the Euclidean distance $dist_{a,b}$ [in meters] between two *Grassland Cells* $G_a$ and $G_b$:

$$dist_{a,b} = \sqrt{\left(coord_x^a - coord_x^b\right)^2 + \left(coord_y^a - coord_y^b\right)^2} \cdot size_{hab} \qquad \text{(S1-10)}$$

If $dist_{a,b} \leq rad_{disp}$ (Table S1-2), the cells belong to the same neighborhood.

Finding potential neighbors for LDD into one of the cardinal directions $DIR = \{N, NE, E, SE, S, SW, W, NW\}$ requires additional information about the surroundings of the source *Grassland Cell* $G_a$. If none of the *Populations* $P_c \in N_a^{dir}$ fulfills the home range constraint $dist_{a,c} \leq rad_{disp}$, the nearest cell $P_{LDD} \in N_{a,LDD}^{dir} \subset N_a^{dir}$ is selected as long distance neighbor in direction $dir$. Here, $N_a^{dir}$ represents a set of all *Populations* located within a 90-degree angle in direction $dir \in DIR$ of cell $G_a$ and $N_{a,LDD}^{dir}$ a subset of slightly narrowed angle outside the dispersal selected neighbor during the dispersal process (Eqn. S1-14) and a probability to survive the dispersal (Eqn. S1-15):

$$pref_{a,b} = \frac{1/\left[dist_{a,b}\right]^{pref^{near}}}{\sum_{P_n \in \mathrm{N}_a}\left[dist_{a,n}\right]^{pref^{near}}} \tag{S1-13}$$

$$p_{a,b}^{find} = \left[rGrass_{a,b}^{at}\right]^{(1-sight_{disp})} \tag{S1-14}$$

$$p_{a,b}^{surv} = \exp^{-decay_{disp}\cdot\left[1-rGrass_{a,b}^{to}\right]\cdot dist_{a,b}} \tag{S1-15}$$

Table S1-2 contains the parameter values used to define the dispersal process of the LMG and their usage is described in the following. The size of the parameter $pref^{near} \geq 0$ in Eqn. S1-13 defines to what extent nearby cells are preferred over more distant ones, where a value of zero leads to equal preference independent of the distance. Finding a selected neighbor (Eqn. S1-14) depends on the relative grassland cover $rGrass_{a,b}^{at}$ in the same distance as the destined cell (i.e., the actual number of grassland cells relative to the potential number), and the ability of the species to locate suitable neighbors given by the parameter $sight_{disp} \in [0,1]$, where a value of zero defines a success rate equal to the grassland ratio and a value of one a 100 % success rate. The probability to survive the dispersal process (Eqn. S1-15) depends on the relative grassland cover $rGrass_{a,b}^{to}$ in the perimeter of a radius lower than the distance to the destined cell and a distance dependent decay rate $decay_{disp} \in [0,1]$, where a value of zero equals 100 % and a value of one minimum survival probability. Furthermore, the dispersers are subject to dispersal mortality which is the difference of the sum of all the dispersal probabilities multiplied by the *base dispersal rate*:

$$mort_{disp}^{ima} = \left(1 - \sum_{P_n \in \mathrm{N}_a} p_{a,n}^{disp}\right) \cdot rate_{disp}^{ima} \tag{S1-16}$$

Finally, the eight sub regions $N_{a,LDD}^{dir} \subset R$ representing all cells found outside the dispersal radius in the direction $dir \in DIR$ of an originating *Grassland Cell* $G_A$ are determined as follows:

$$N_a^N = \left\{\forall G_n \in R \backslash \{G_a\} \mid dist_{a,n}^x < dist_{a,n}^y \wedge coord_y^n < coord_y^a \wedge dist_{a,n} > rad_{disp}\right\} \tag{S1-17}$$

$$N_a^{NE} = \left\{\forall G_n \in R \backslash \{G_a\} \mid coord_x^n > coord_x^a \wedge coord_y^n < coord_y^a \wedge dist_{a,n} > rad_{disp}\right\} \tag{S1-18}$$

$$N_a^E = \left\{\forall G_n \in R \backslash \{G_a\} \mid coord_x^n > coord_x^a \wedge dist_{a,n}^y < dist_{a,n}^x \wedge dist_{a,n} > rad_{disp}\right\} \tag{S1-19}$$

$$N_a^{SE} = \left\{\forall G_n \in R \backslash \{G_a\} \mid coord_x^n > coord_x^A \wedge coord_y^n > coord_y^a \wedge dist_{a,n} > rad_{disp}\right\} \tag{S1-20}$$

$$N_a^S = \left\{\forall G_n \in R \backslash \{G_a\} \mid dist_{a,n}^x < dist_{a,n}^y \wedge coord_y^a > coord_y^a \wedge dist_{a,n} > rad_{disp}\right\} \tag{S1-21}$$

$$N_a^{SW} = \left\{\forall G_n \in R \backslash \{G_a\} \mid coord_x^n < coord_x^a \wedge coord_y^n > coord_y^a \wedge dist_{a,n} > rad_{disp}\right\} \tag{S1-22}$$

$$N_a^W = \left\{\forall G_n \in R \backslash \{G_a\} \mid coord_x^n < coord_x^a \wedge dist_{a,n}^y < dist_{a,n}^x \wedge dist_{a,n} > rad_{disp}\right\} \tag{S1-23}$$

$$N_a^{NW} = \left\{\forall G_n \in R \backslash \{G_a\} \mid coord_x^n < coord_x^a \wedge coord_y^n < coord_y^a \wedge dist_{a,n} > rad_{disp}\right\} \tag{S1-24}$$

We defined the sub regions in such a way that each of them overlaps with both neighbors in clockwise and counterclockwise direction to minimize LDD to cases where the area in direction of interest is largely empty within the home range. Figure 5 (main manuscript) visualizes the areas belonging to either of the sub regions.

#### 7.6 Bilinear climate interpolation

M2: This submodel is called every time step to achieve heterogeneous, gradual values at the location of each *Grassland Cell* using bilinear interpolation of the climate data from the closest adjacent neighbors. This is done by weighing the distances from a *Grassland Cell* $G_a$ to the center of the (up to) four *Climate Cells* $\{\Omega_{a,NE}, \Omega_{a,SE}, \Omega_{a,SW}, \Omega_{a,NW}\}$ into secondary cardinal directions of $G_a$. The resulting bilinear weights $w_{bilin}^{a,dir}$ are multiplied with their respective climate values $\omega_{clim}^{a,dir}$ and then summed to achieve the value at $G_a$. The interpolated climate value $\omega_{clim}^{a}$ for cell $G_a$ is then calculated as follows.

$$\omega_{clim}^{a} = \sum_{dir \in DIR_{sec}} w_{bilin}^{a,dir} \cdot \omega_{clim}^{a,dir} \tag{S1-25}$$

$$w_{bilin}^{a,dir} = 1 - \frac{\left(size_{clim} - dist_{a,dir}^{x}\right) \cdot \left(size_{clim} - dist_{a,dir}^{y}\right)}{[size_{clim}]^2} \tag{S1-26}$$

$$dist_{a,dir}^{xy} = \left|coord_{xy}^{a} - center_{xy}^{a,dir}\right| \cdot size_{hab} \tag{S1-27}$$

Here, $DIR_{sec} \subset DIR$ (see Section 7.5) are the secondary cardinal directions {NE, SE, SW, NW} and $\omega_{clim}^{a,dir}$ is the projected value in the *Climate Cell* $\Omega_{a,dir}$ into direction $dir$ of $G_a$. Parameters $size_{clim}$ of a *Climate Cell* and $size_{hab}$ of a *Grassland Cell* were introduced in Table S1-2. The value $dist_{a,dir}^{xy}$ for the distances in x- or y-direction are calculated using the respective coordinate of the geometric center of a *Climate Cell* ($center_{x,y}^{a,dir}$ , Table S1-2). Figure 4 (main manuscript) illustrates the calculation of the weights for a single grassland cell using a simplified geometric example. Please refer to Supplement S4 for a description of the mapping between climate and grassland cells, and a reference to the calculated weights.
