## Supplementary material for "Large scale PVA modelling of insects in cultivated grasslands: the role of dispersal in mitigating the effects of management schedules under climate change": Illustration of dispersal success

The document contains illustrations for dispersal success of the large marsh grasshopper (LMG) per climate change scenarios (CCS) depending on source habitat and mowing schedule in terms of three evaluation parameters: (Section 1) the maximum *established distance* in meters from a source habitat to a habitat with imago density ≥ 0.002 individuals $m^{-2}$ during a year, (Section 2) the *population size* in total number of eggs in all established habitats by the end of a simulation year, and (Section 3) the *population density* in eggs $m^{-2}$ for all established habitats by the end of a simulation year. The title of each Figure indicates, which evaluation parameter, CCS (Clim-FF, Clim-MOD or Clim-BAU) and mowing schedule (see main manuscript Table 1) applies. Each of the 107 subplots in every Figure represents INDEPENDENT simulation runs, or rather their mean over 10 replicate runs, and depicts the results of the dispersal process from a SINGLE initial population in the center of a cell. Inside each subplot, dots are the log-scaled yearly means of the respective value, solid black lines represents their smoothed trends using a generalized additive model and dashed horizontal grey lines mark the mean of the yearly values during the 60 simulation years. The background color of the subplots (FF=green, MOD=brown, BAU=pink) highlights the latter mean value in comparison to other subplots, where a LIGHTER color represents a lower mean, and reflects in the bottom legend.

#### Section 1: illustration of maximum *established distance*

### Clim-FF
### mean yearly max. established distance

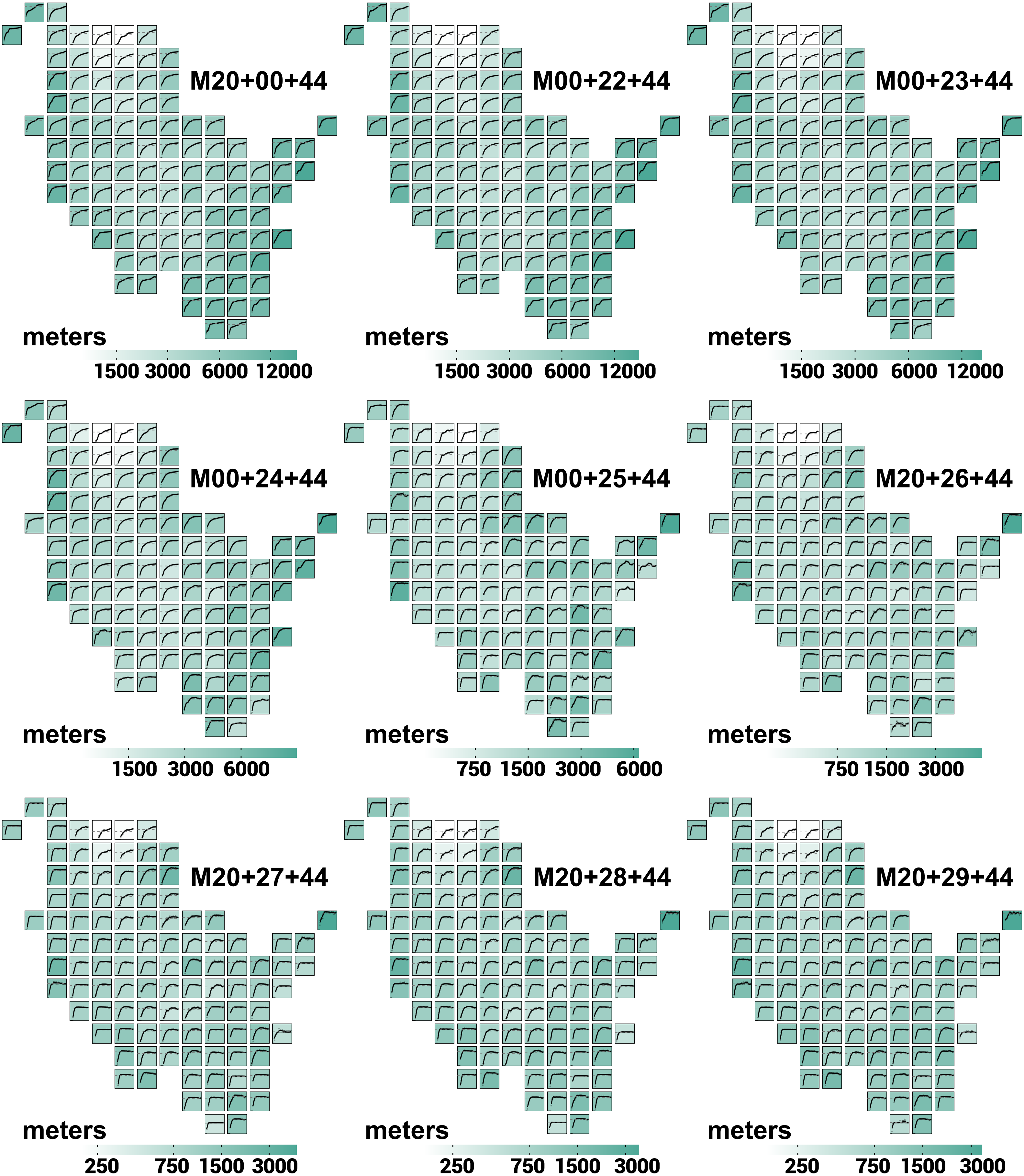

### Clim-FF
### mean yearly max. established distance

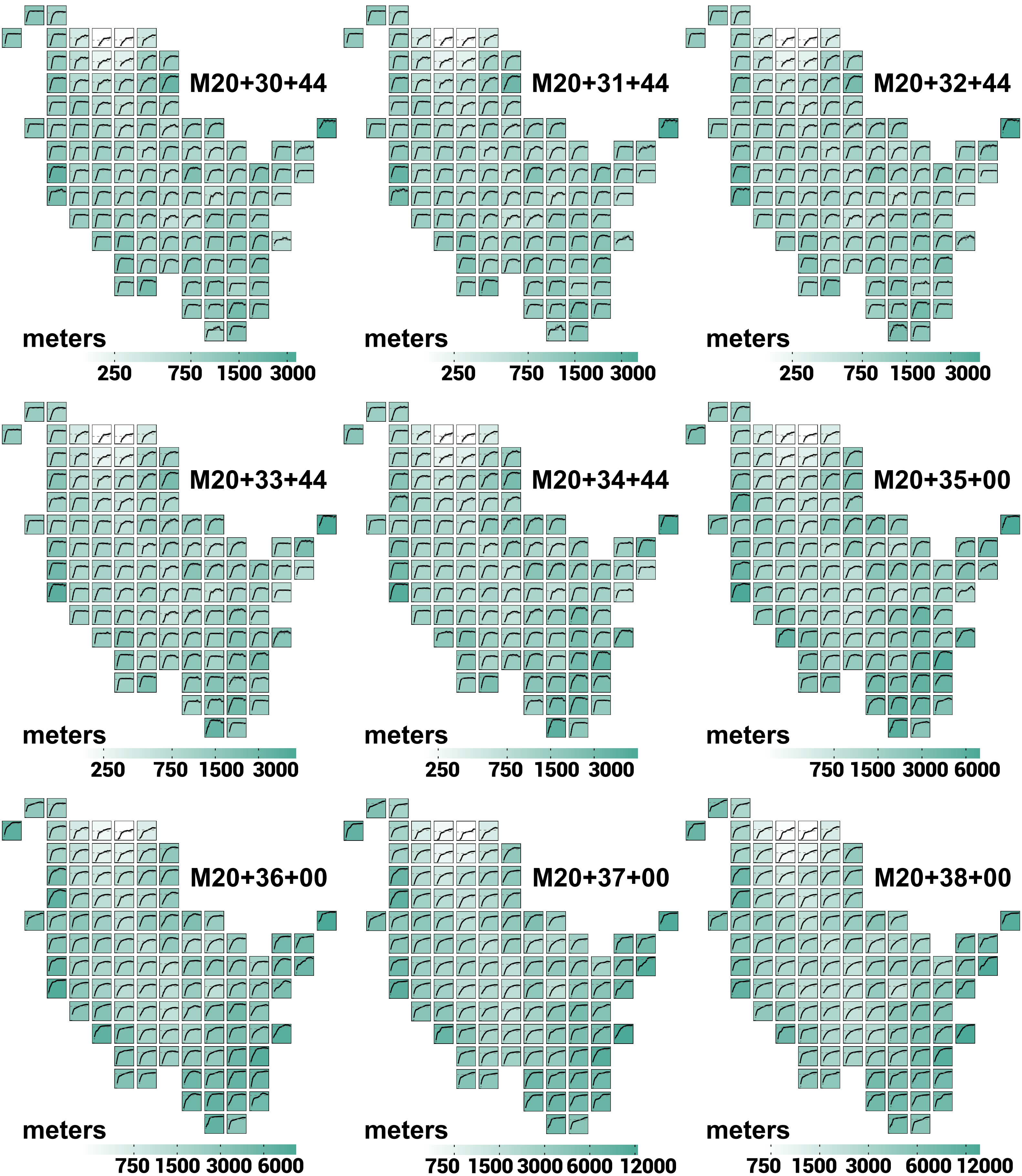

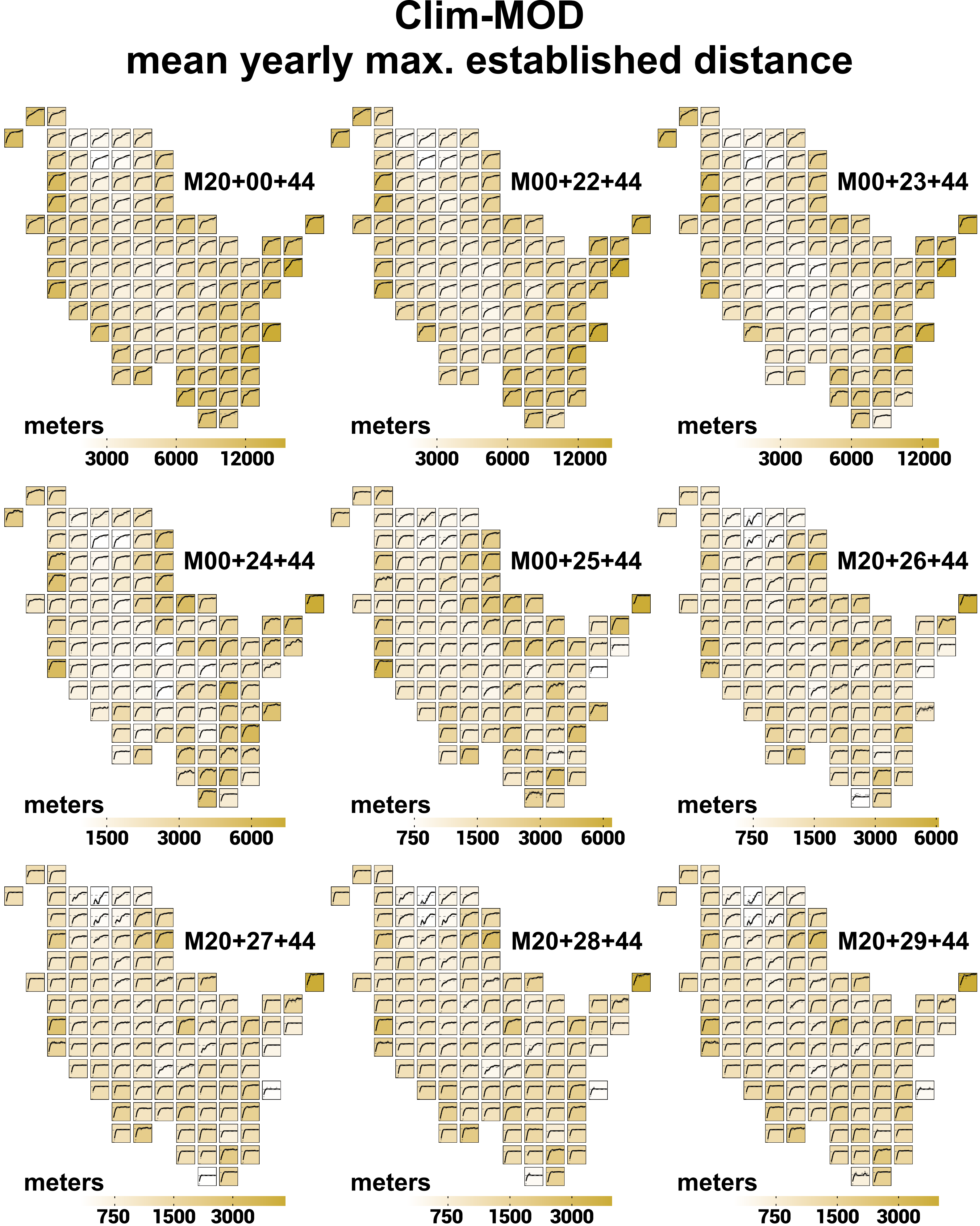

Clim-MOD
mean yearly max. established distance
M20+00+44
M00+22+44
M00+23+44
M00+24+44
M00+25+44
M20+26+44
M20+27+44
M20+28+44
M20+29+44
meters
3000
6000
12000
1500
750

### Clim-MOD
### mean yearly max. established distance

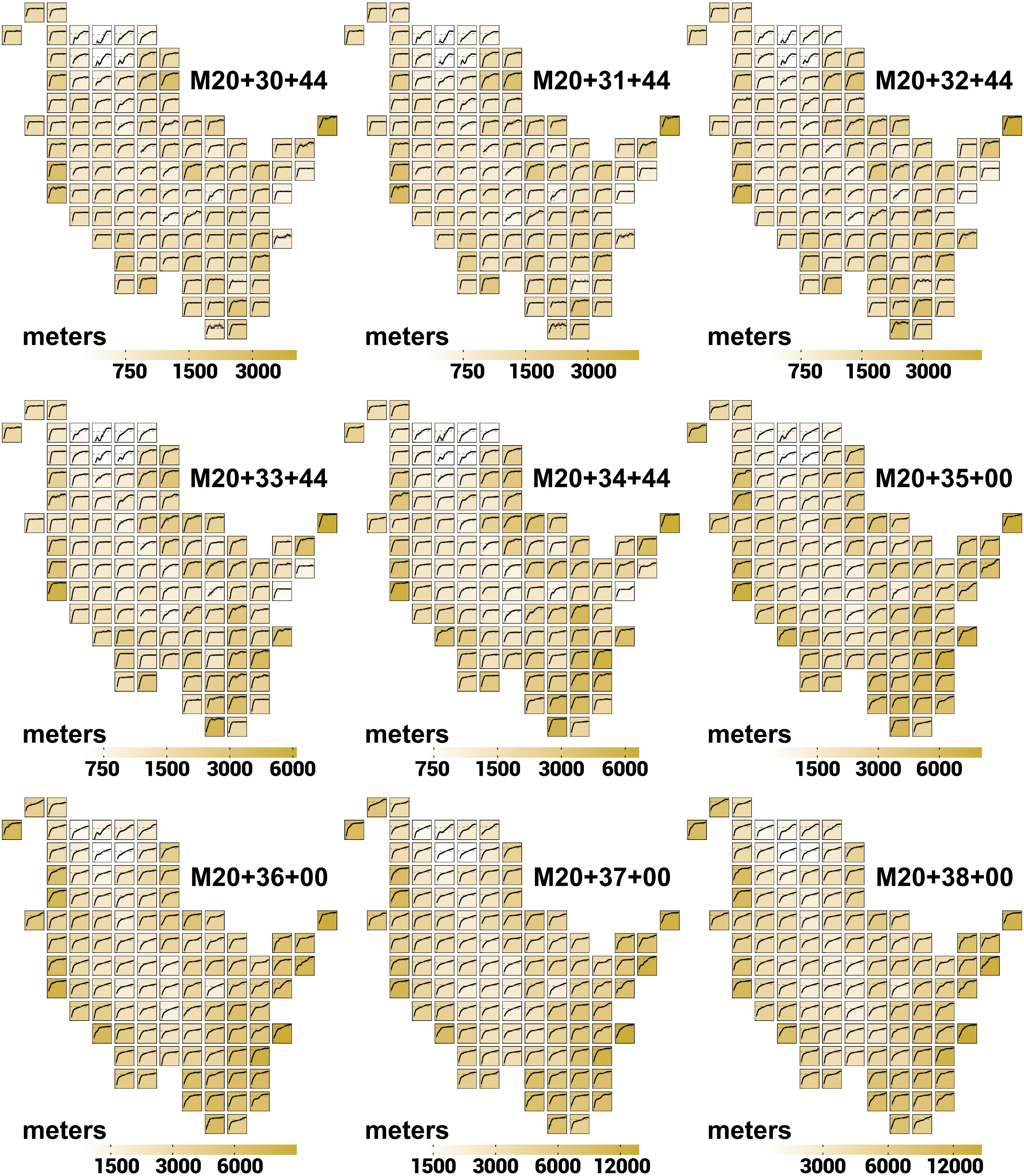

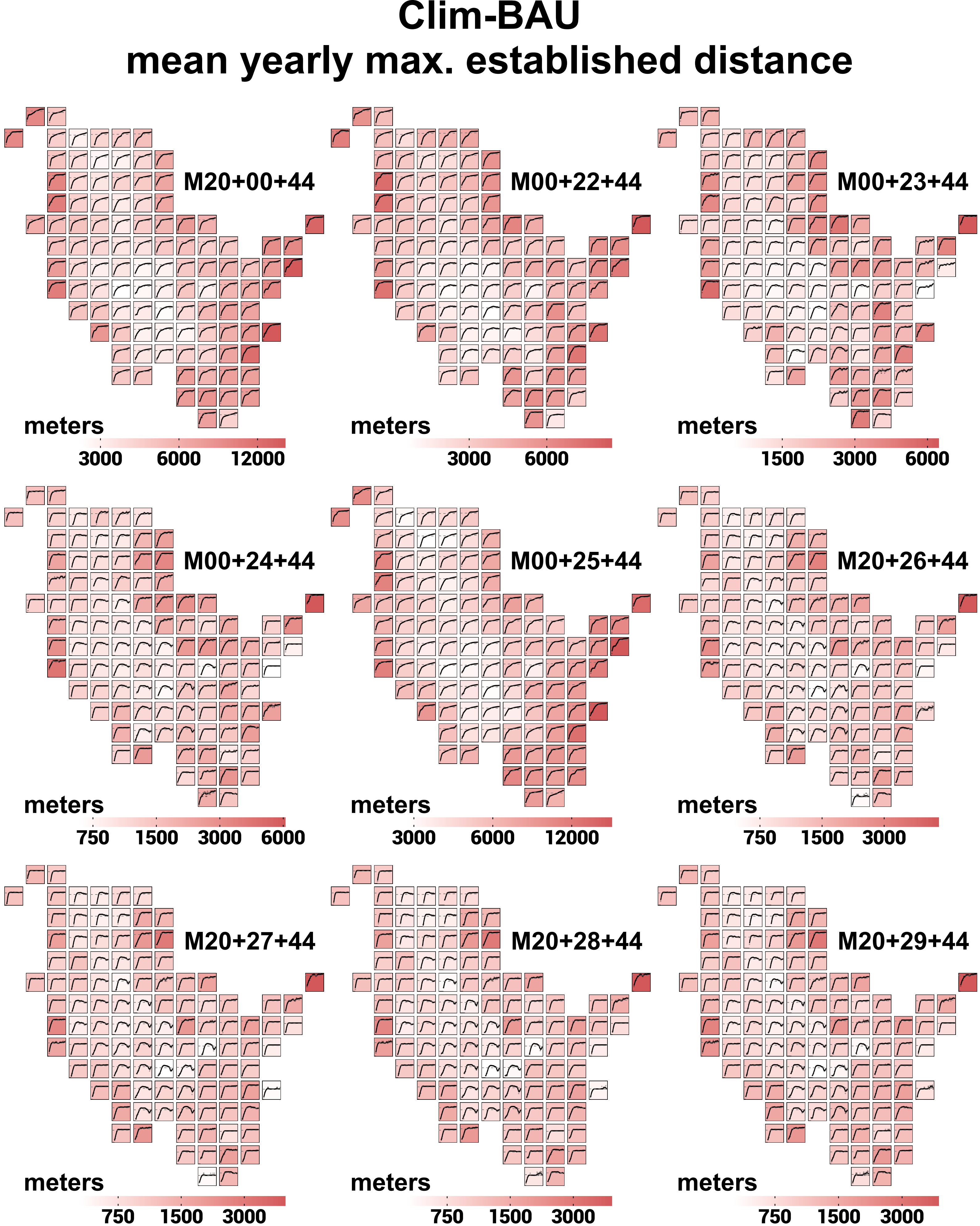
Clim-BAU
mean yearly max. established distance
M20+00+44
meters
3000
6000
12000
M00+22+44
meters
3000
6000
M00+23+44
meters
1500
3000
6000
M00+24+44
meters
750
1500
3000
6000
M00+25+44
meters
3000
6000
12000
M20+26+44
meters
750
1500
3000
M20+27+44
meters
750
1500
3000
M20+28+44
meters
750
1500
3000
M20+29+44
meters
750
1500
3000

### Clim-BAU
### mean yearly max. established distance

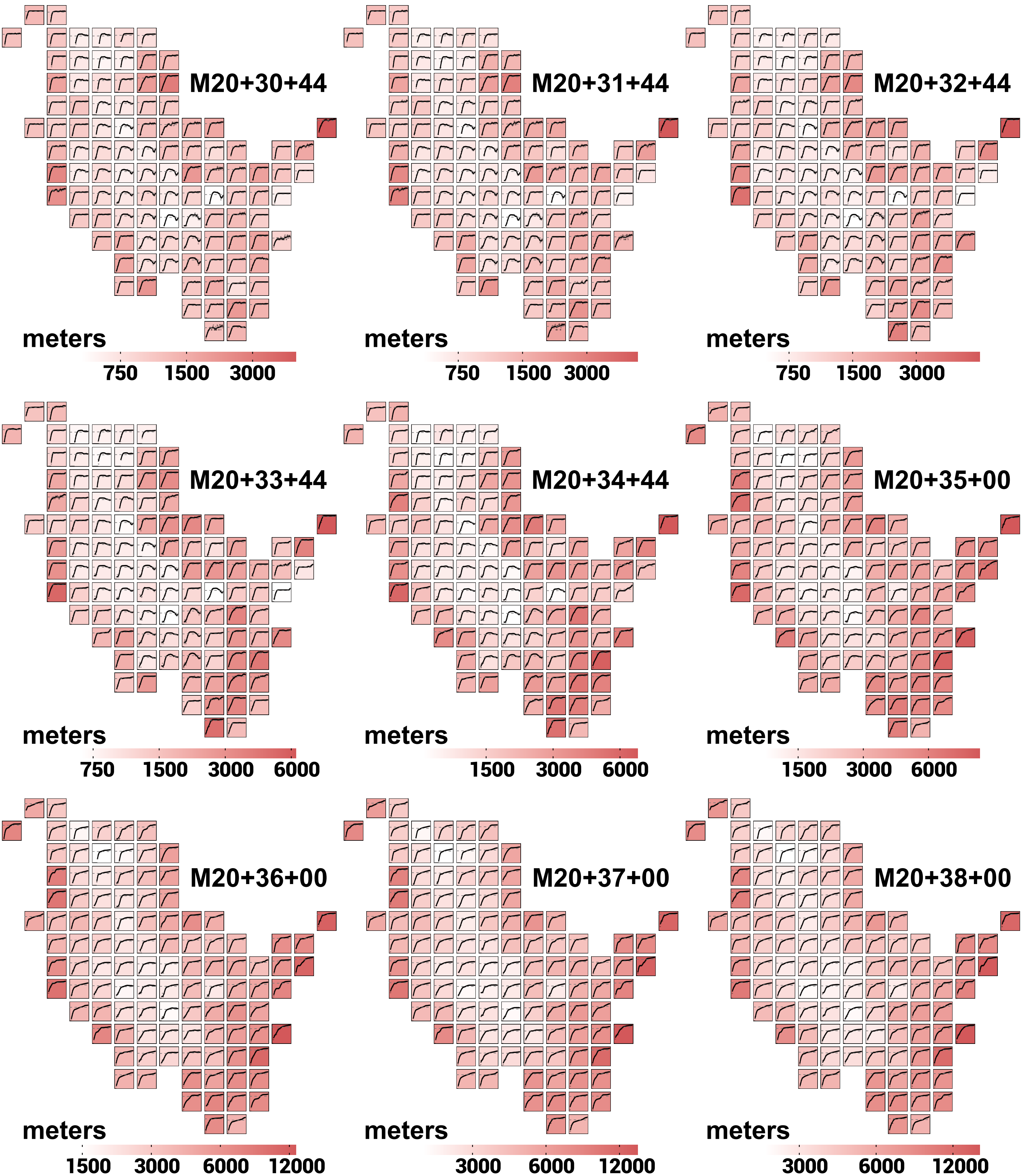

#### Section 2: illustration of *population size*

### Clim-FF
### mean yearly population size

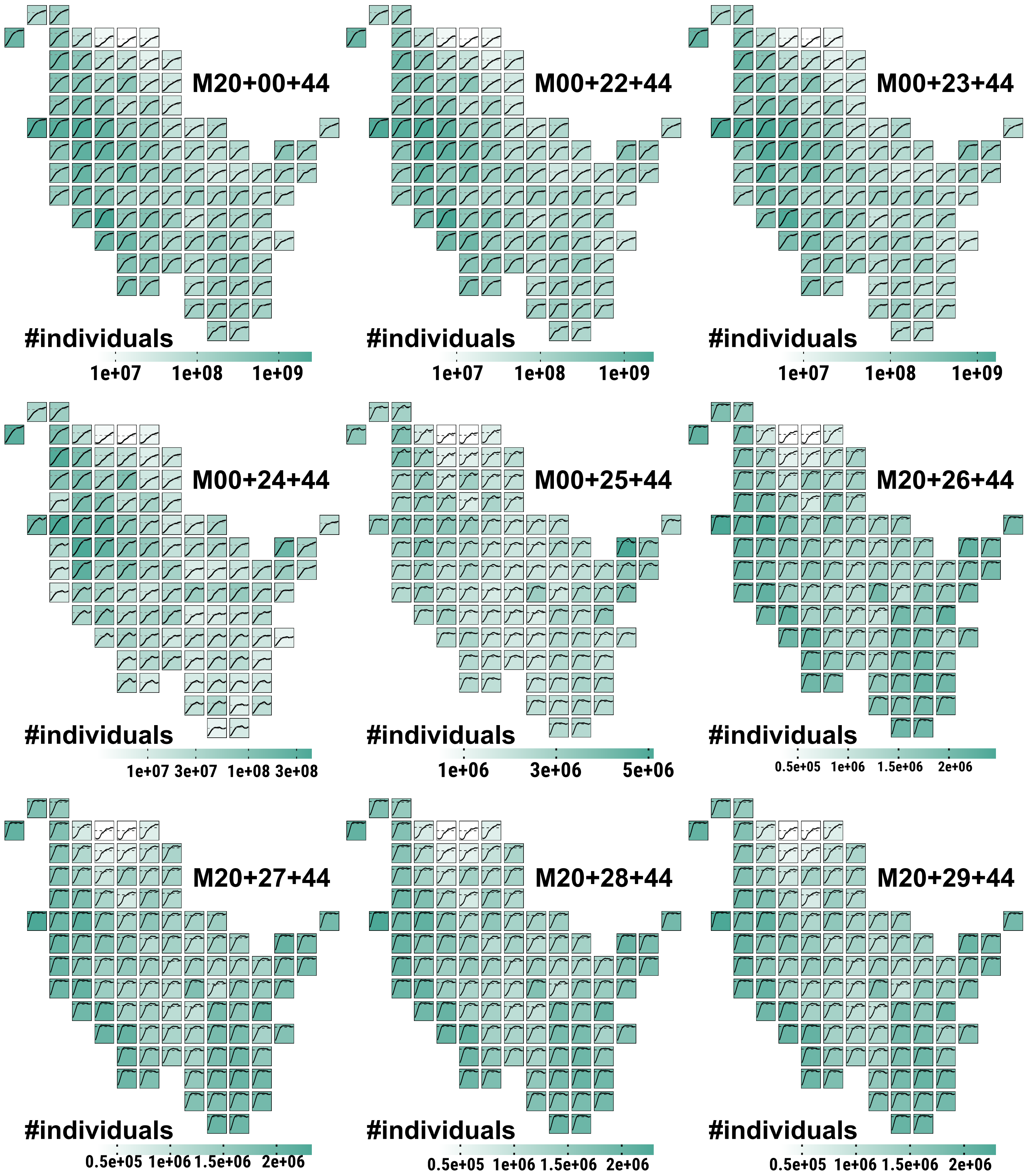

### Clim-FF
### mean yearly population size

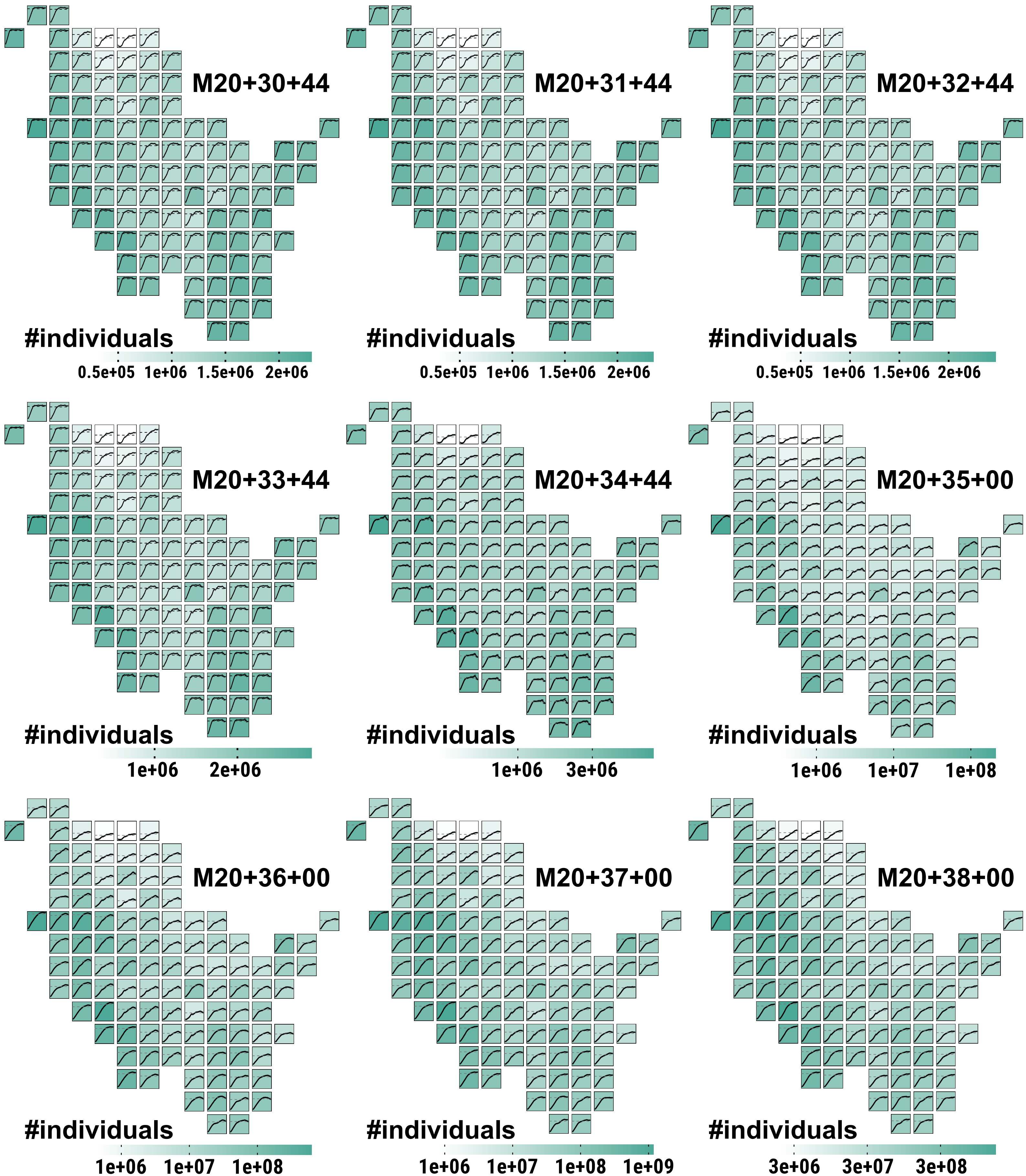

### Clim-MOD
### mean yearly population size

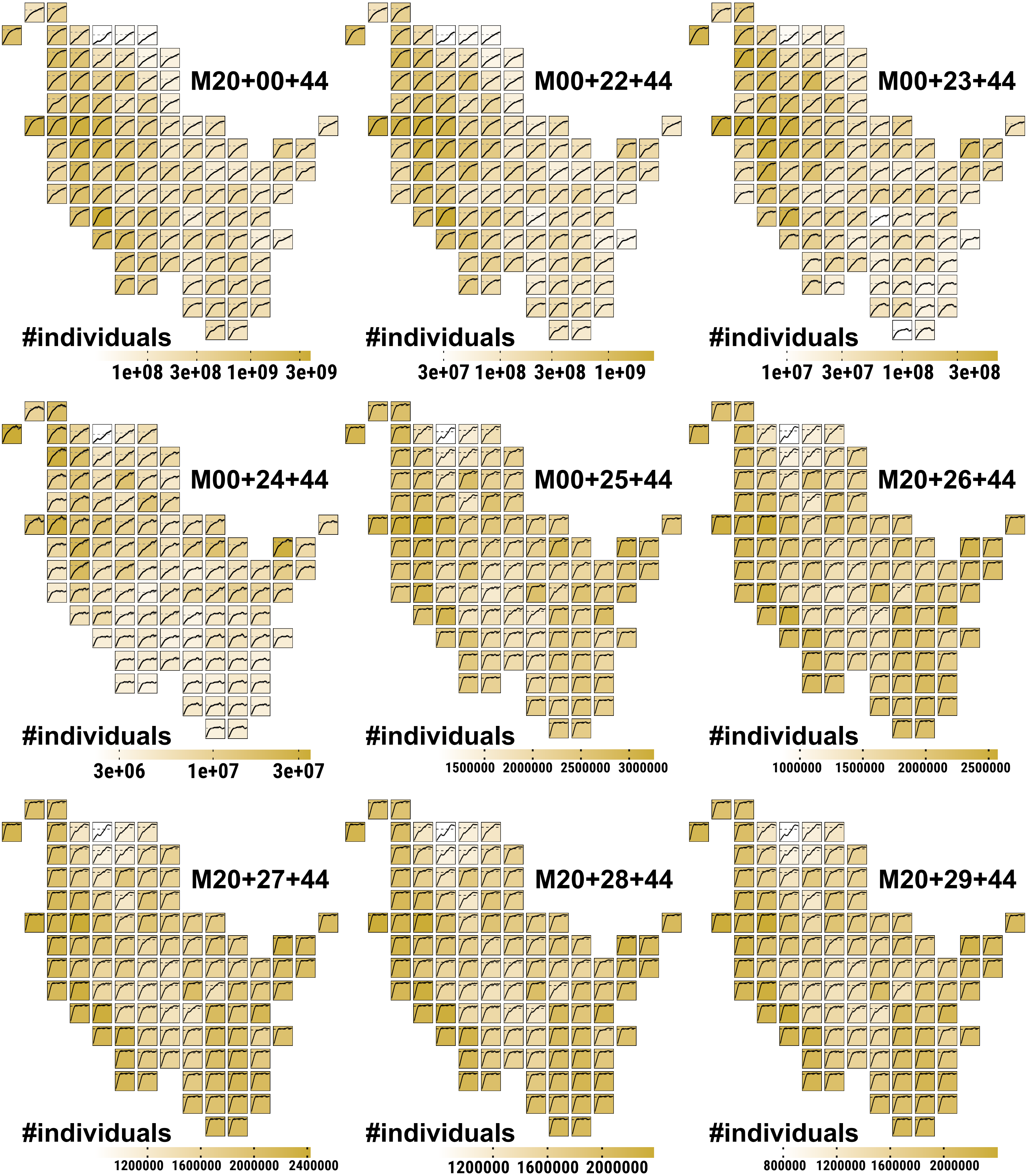

### Clim-MOD
### mean yearly population size

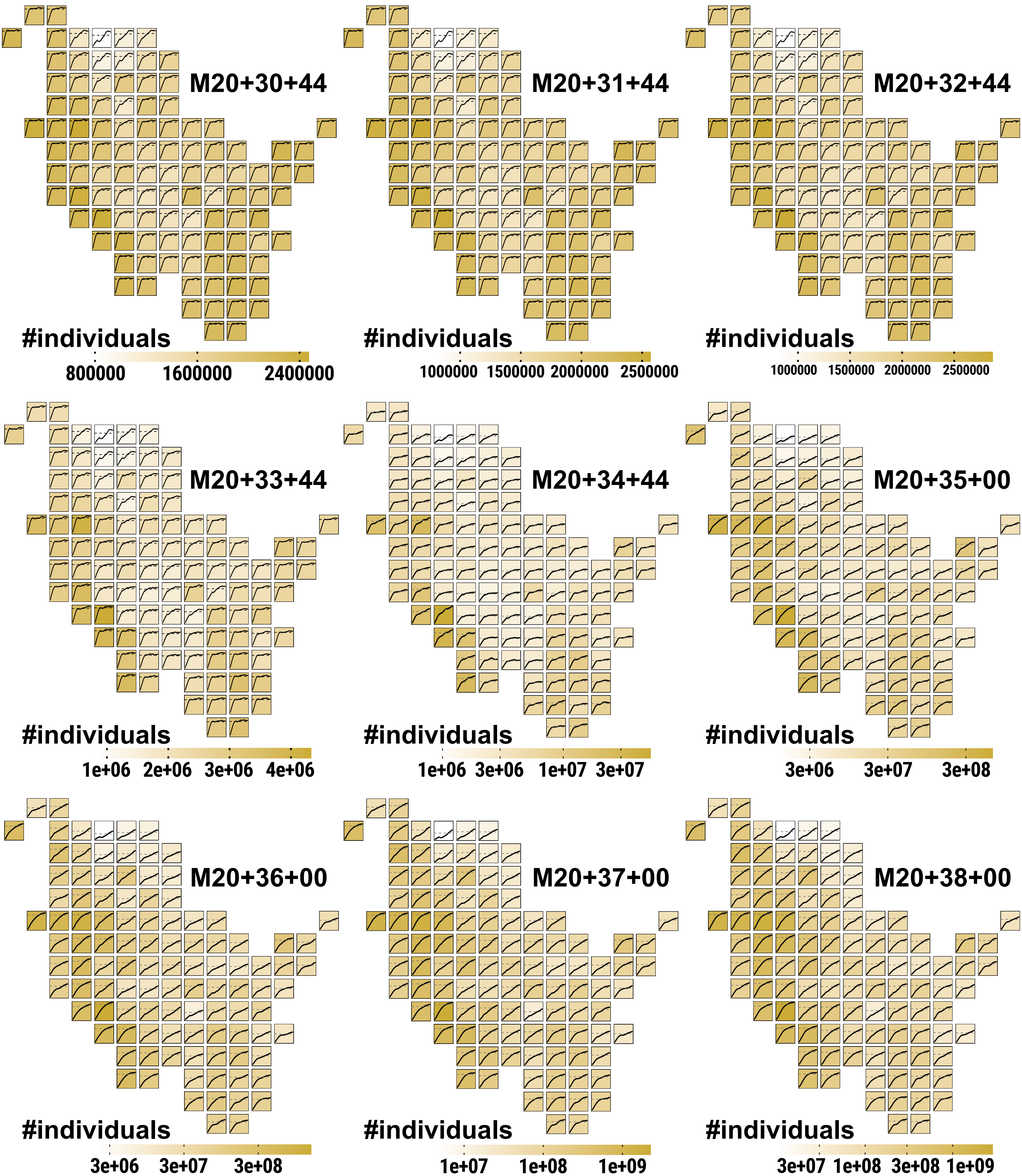

### Clim-BAU
### mean yearly population size

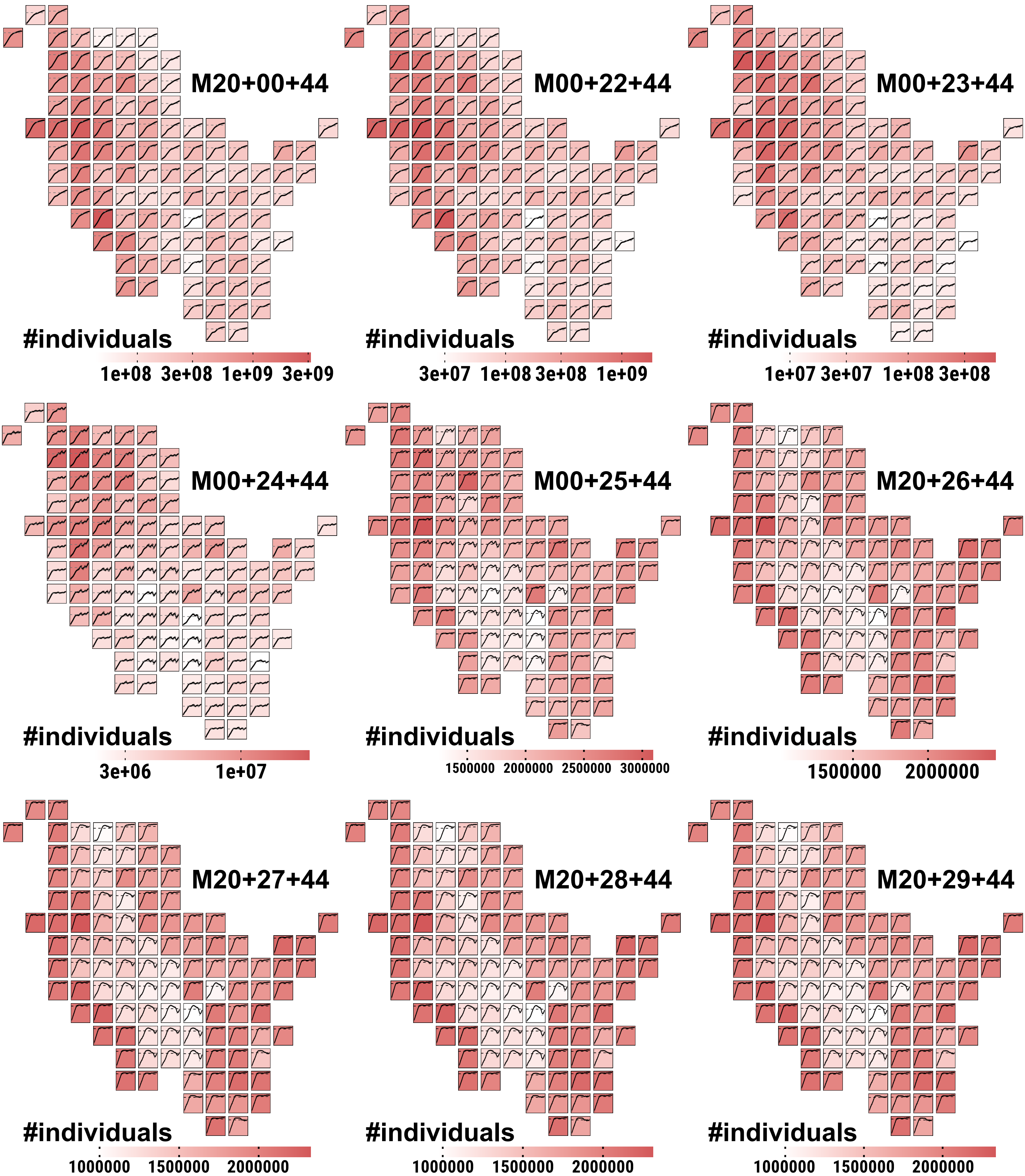

### Clim-BAU
### mean yearly population size

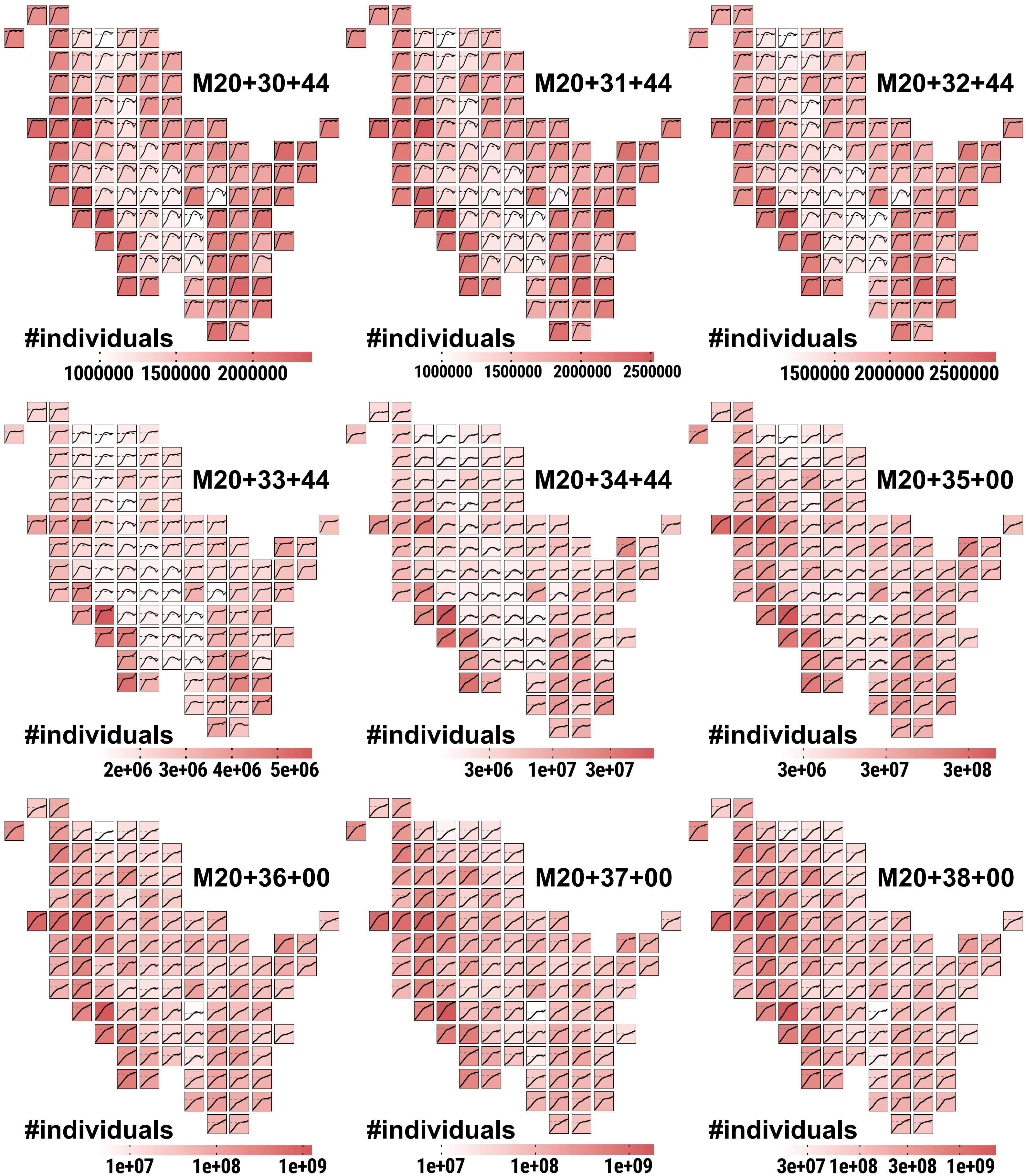

#### Section 3: illustration of *population density*

### Clim-FF
### mean yearly population density

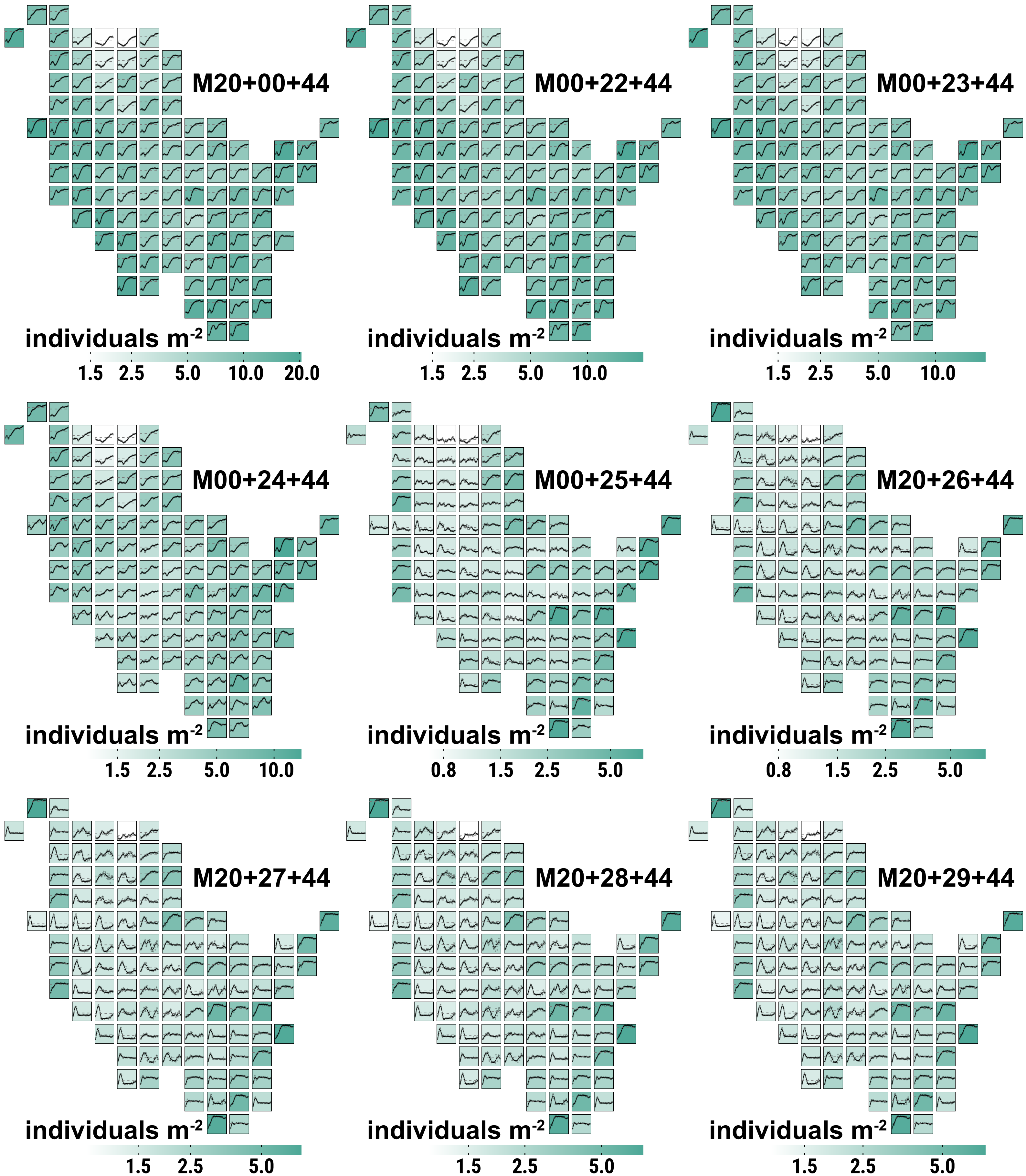

### Clim-FF
### mean yearly population density

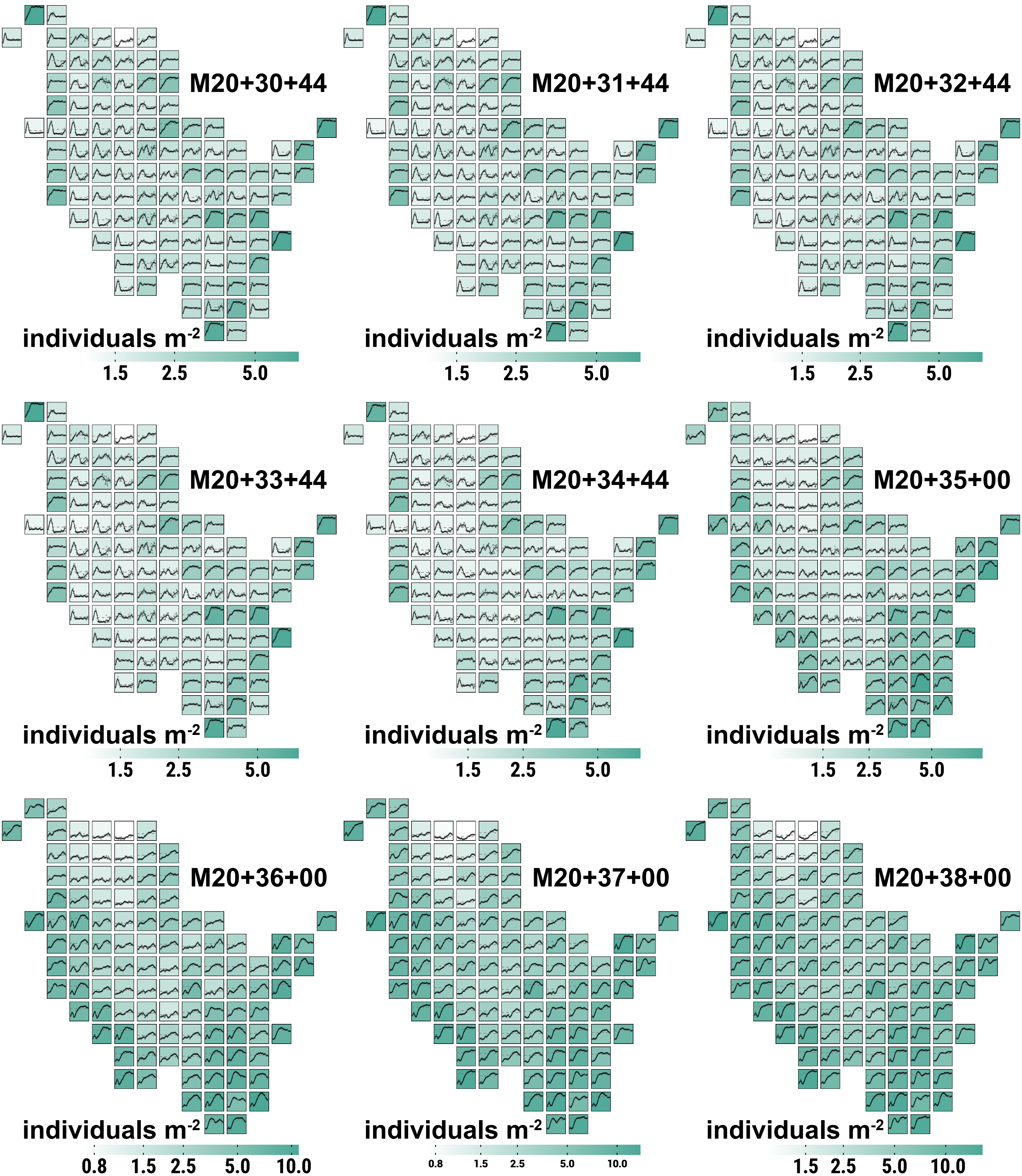

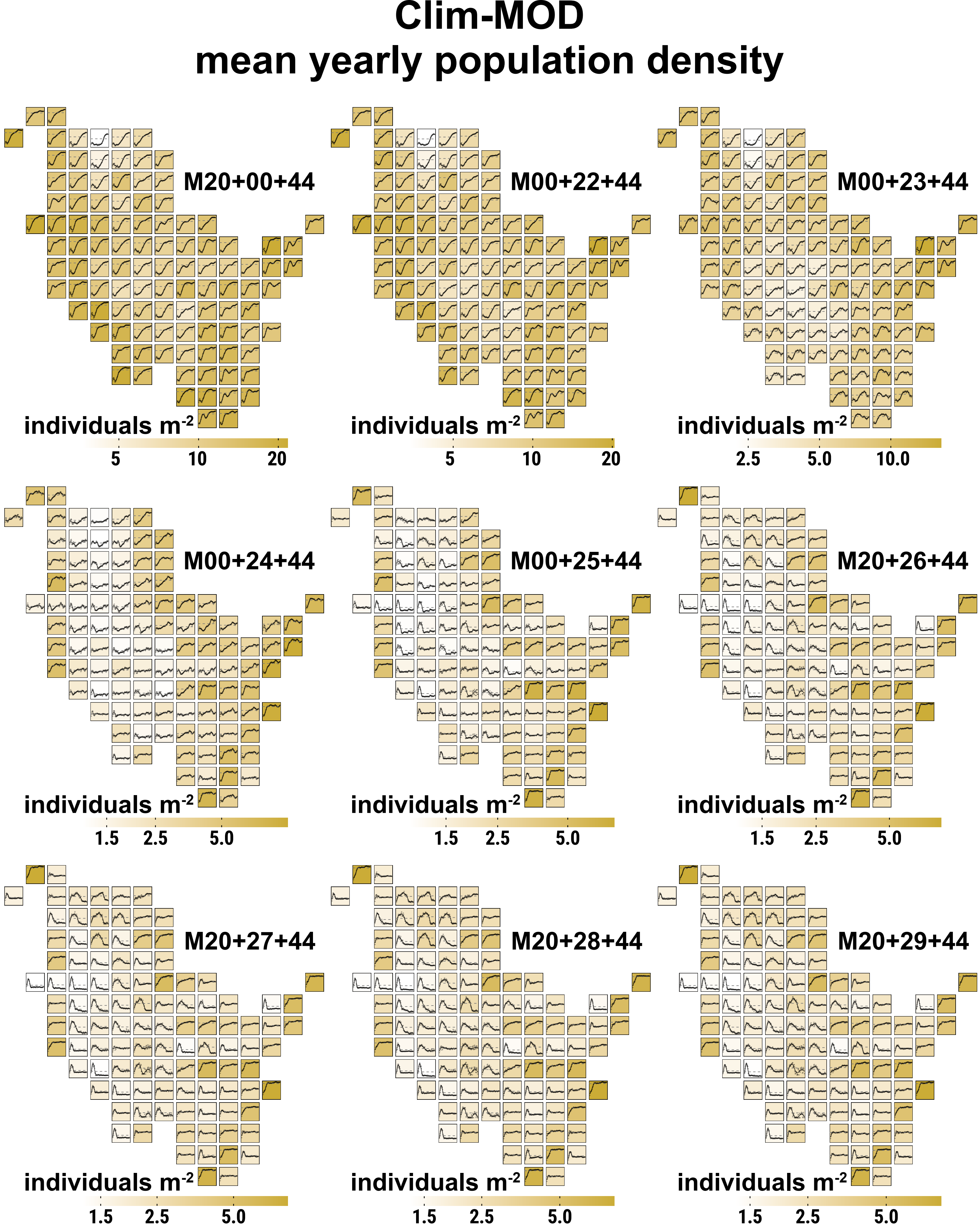

Clim-MOD
mean yearly population density
M20+00+44
M00+22+44
M00+23+44
M00+24+44
M00+25+44
M20+26+44
M20+27+44
M20+28+44
M20+29+44
individuals m-2
5
10
20
2.5
5.0
10.0
1.5

### Clim-MOD
### mean yearly population density

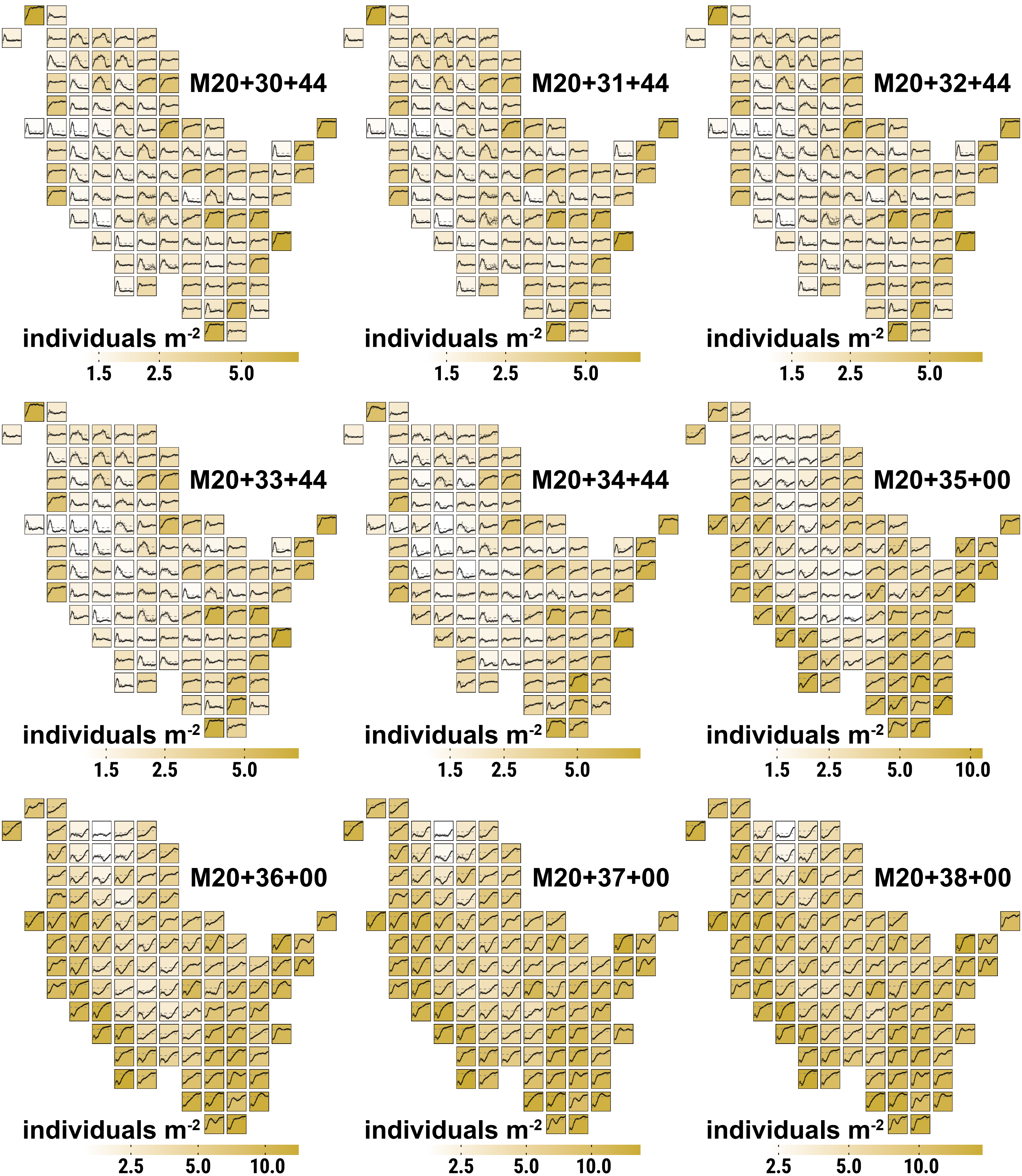

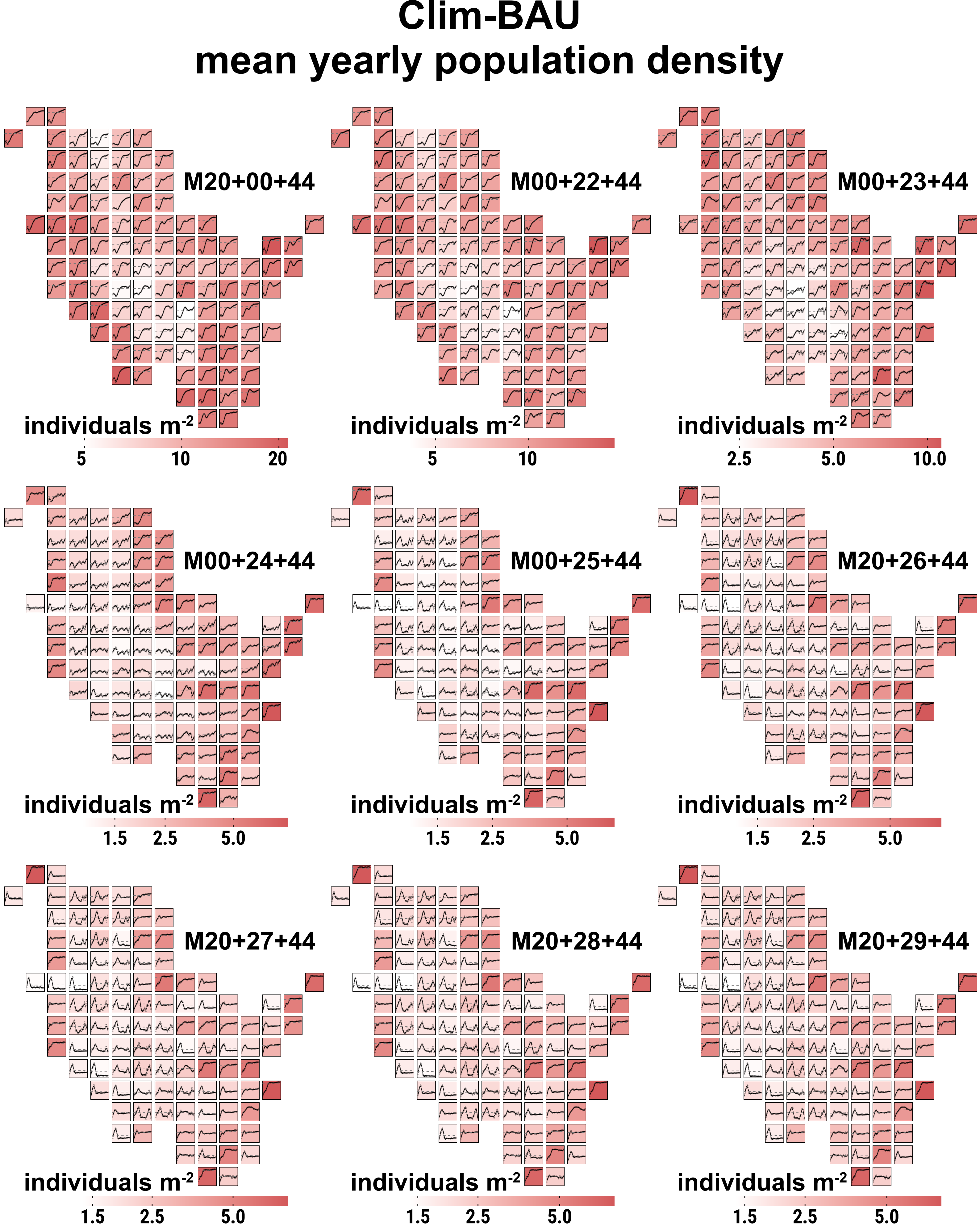
Clim-BAU
mean yearly population density
M20+00+44
individuals m-2
5
10
20
M00+22+44
individuals m-2
5
10
M00+23+44
individuals m-2
2.5
5.0
10.0
M00+24+44
individuals m-2
1.5
2.5
5.0
M00+25+44
individuals m-2
1.5
2.5
5.0
M20+26+44
individuals m-2
1.5
2.5
5.0
M20+27+44
individuals m-2
1.5
2.5
5.0
M20+28+44
individuals m-2
1.5
2.5
5.0
M20+29+44
individuals m-2
1.5
2.5
5.0

### Clim-BAU
### mean yearly population density

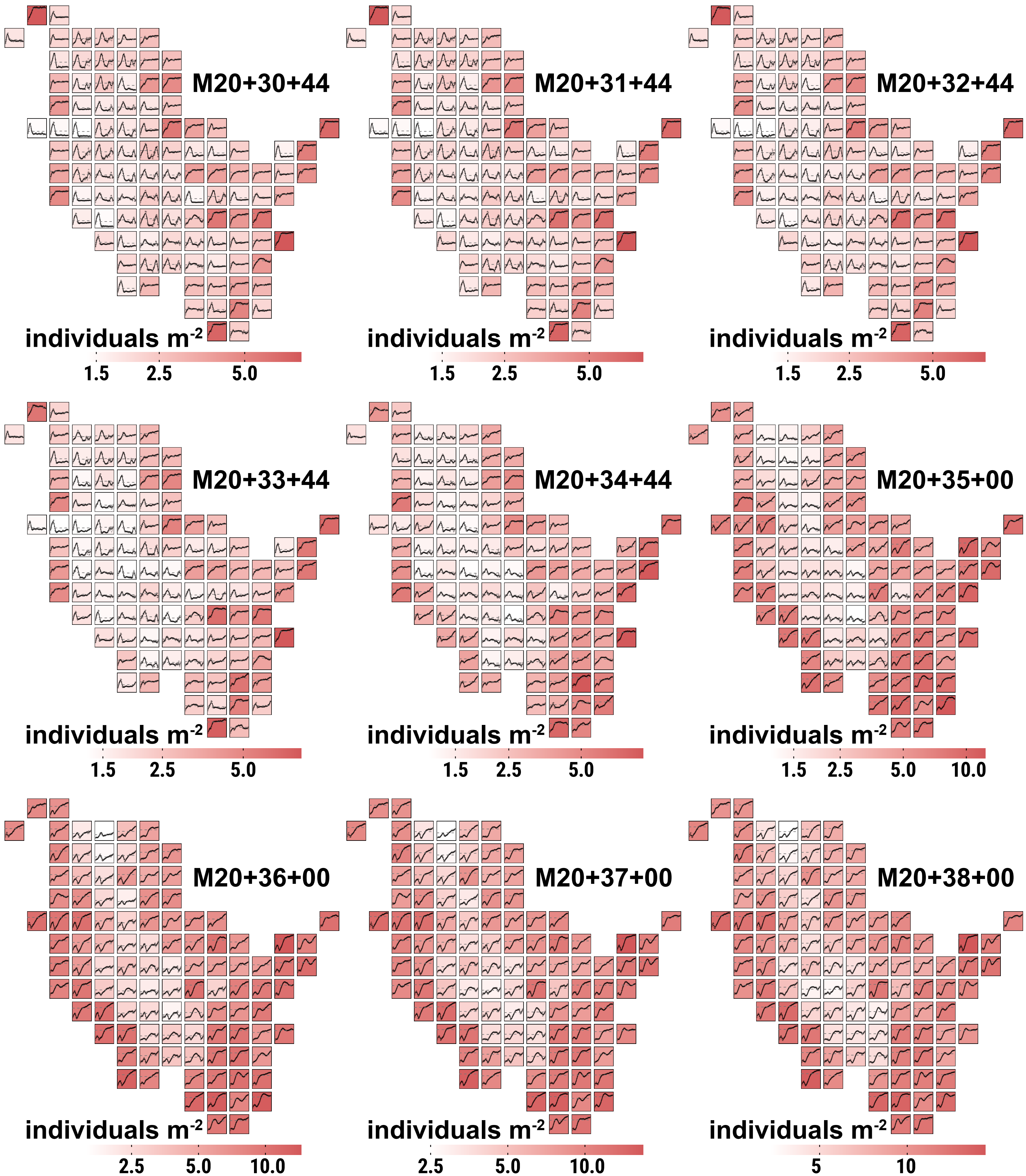
