## Supplementary material for "Large scale PVA modelling of insects in cultivated grasslands: the role of dispersal in mitigating the effects of management schedules under climate change": Illustration of differences in dispersal success

The document contains illustrations of the difference (delta) in dispersal success of the large marsh grasshopper (LMG) between climate change scenarios (CCS) depending on source habitat and mowing schedule in terms of four evaluation parameters: (Section 1) the maximum *established distance* in meters from a source habitat to a habitat with imago density ≥ 0.002 individuals $m^{-2}$ during a year, (Section 2) the *population size* in total number of eggs in all established habitats at the end of a simulation year, and (Section 3) the *population density* in eggs $m^{-2}$ for all established habitats at the end of a simulation year. The title of each Figure indicates, which evaluation parameter, CCS delta (Clim-FF vs. Clim-MOD, Clim-FF vs. Clim-BAU or Clim-MOD vs. Clim-BAU) and mowing schedule (see main manuscript Table 1) applies. Values were determined per replicate by subtracting the yearly parameter values of the CCS mentioned second from the CCS mentioned first in the title and then calculating the replicate mean of the delta. Each of the 107 subplots in every Figure represents INDEPENDENT simulation runs, or rather their mean over 10 replicate runs, and depicts the results of the dispersal process from a SINGLE initial population in the center of a cell. Inside each subplot, dots are the log-scaled yearly means of the differences, solid black lines represents their smoothed trends using a generalized additive model, solid horizontal grey lines mark zero and dashed horizontal grey lines mark the mean of the yearly values during the 60 simulation years. The background color of the subplots highlight which of the respective CCS on average shows the higher differences during the 60 simulation years, where a LIGHTER color represents lower average difference. GREEN cells are in favor of Clim-FF, BROWN cells in favor of Clim-MOD and PINK cells in favor of Clim-BAU. These colors reflect in the bottom legend.

#### Section 1: difference in maximum *established distance*

### Clim-FF vs. Clim-MOD
### difference in mean yearly max. established distance

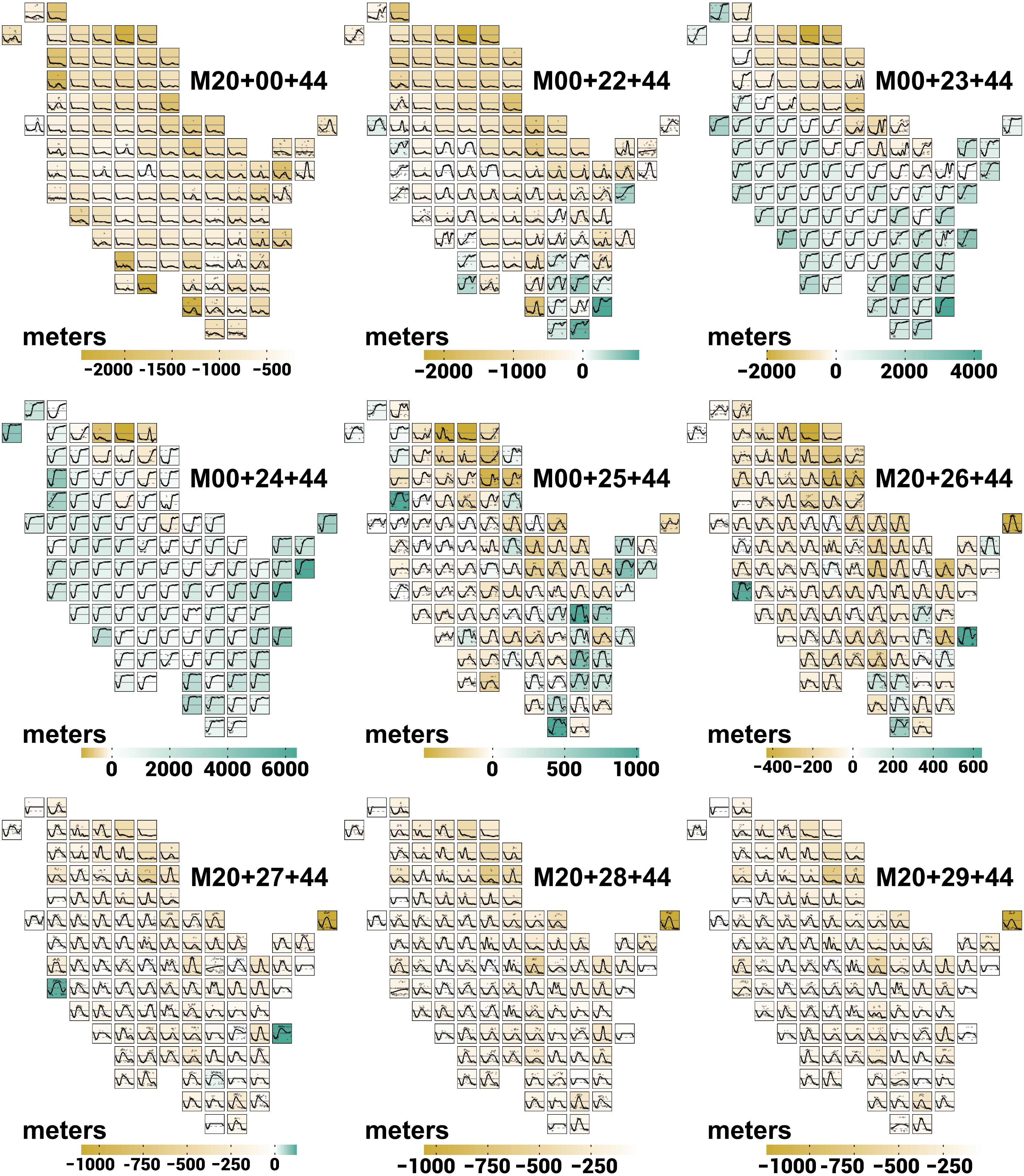

### Clim-FF vs. Clim-MOD
### difference in mean yearly max. established distance

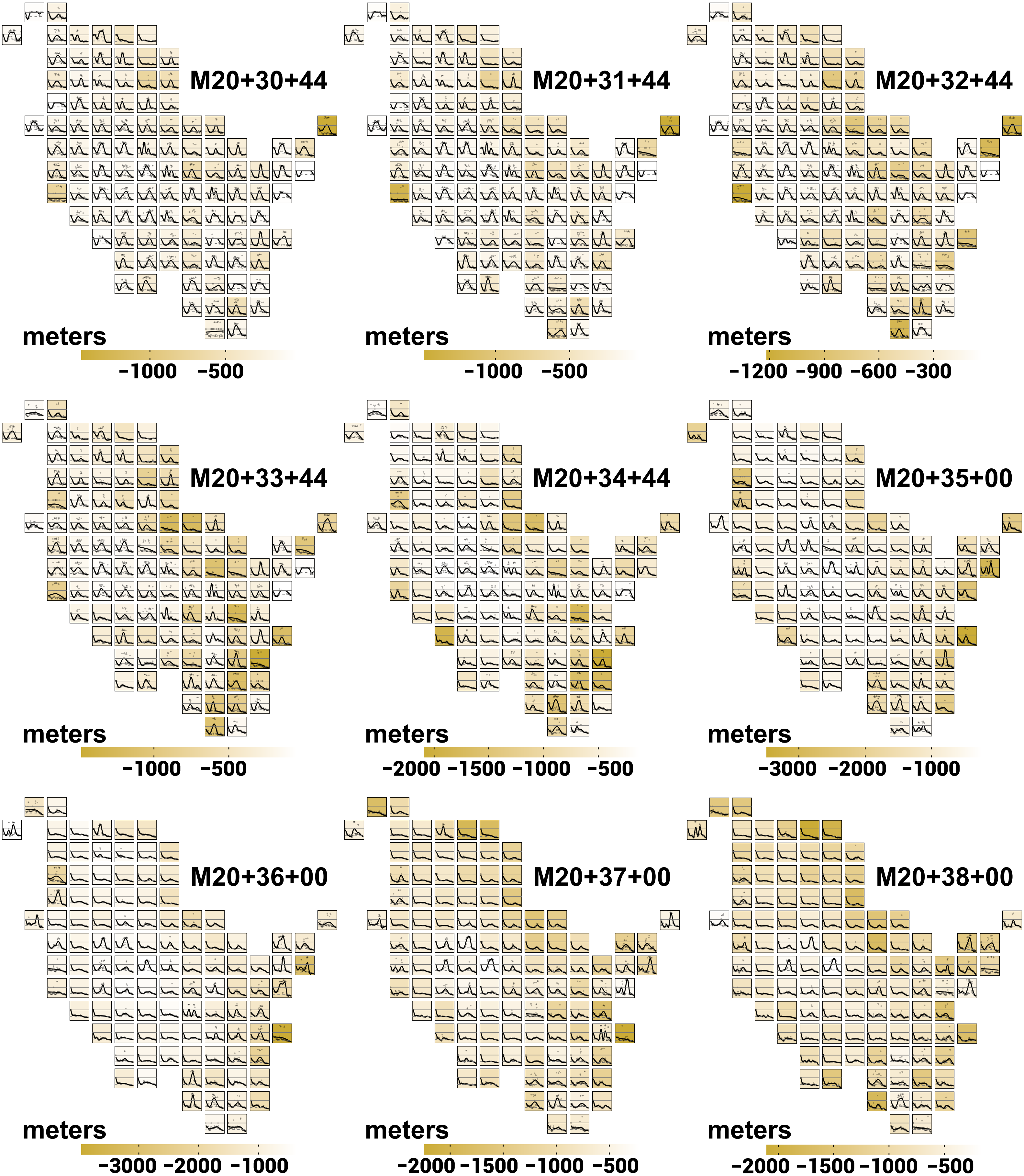

### Clim-FF vs. Clim-BAU
### difference in mean yearly max. established distance

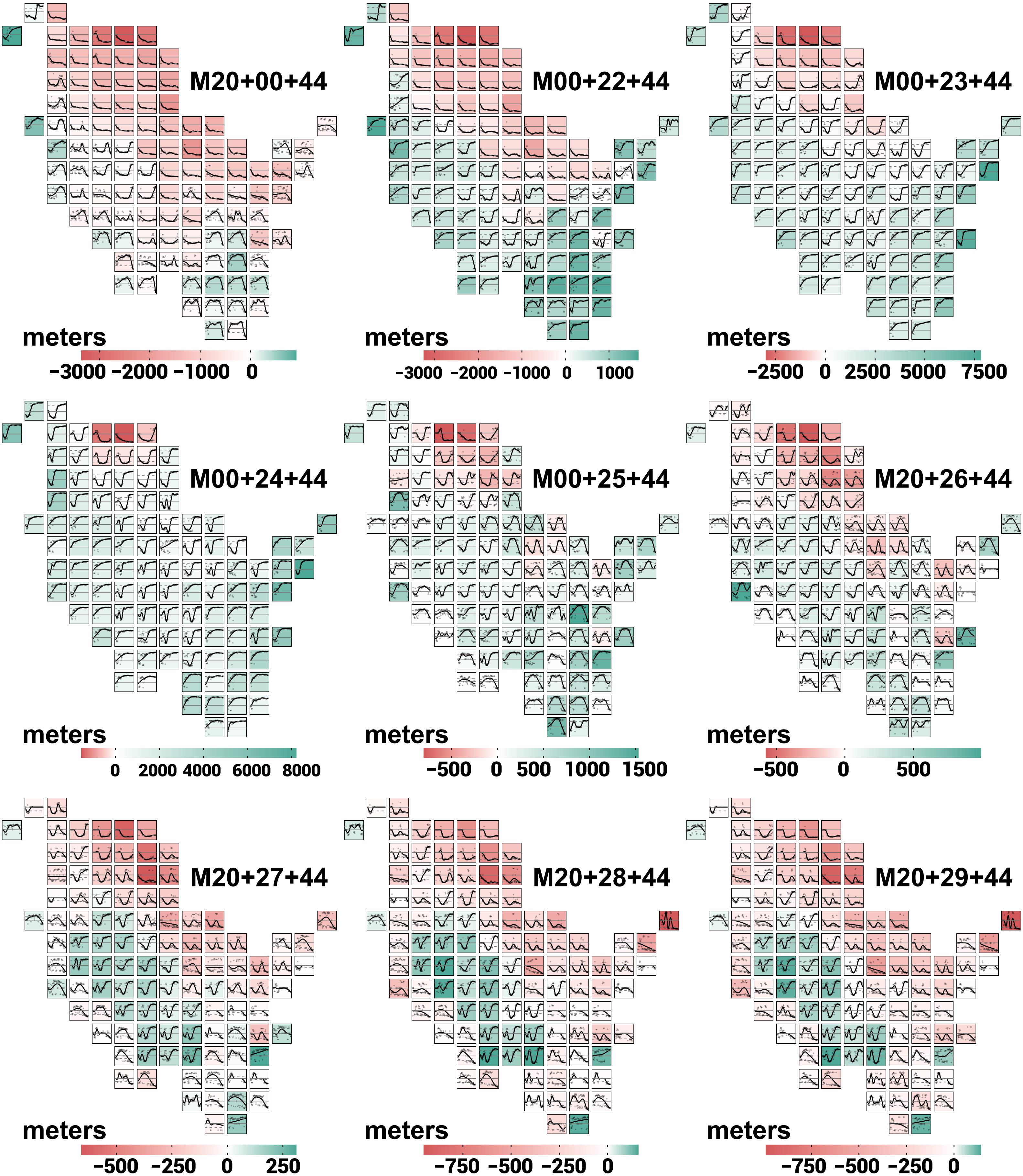

### Clim-FF vs. Clim-BAU
### difference in mean yearly max. established distance

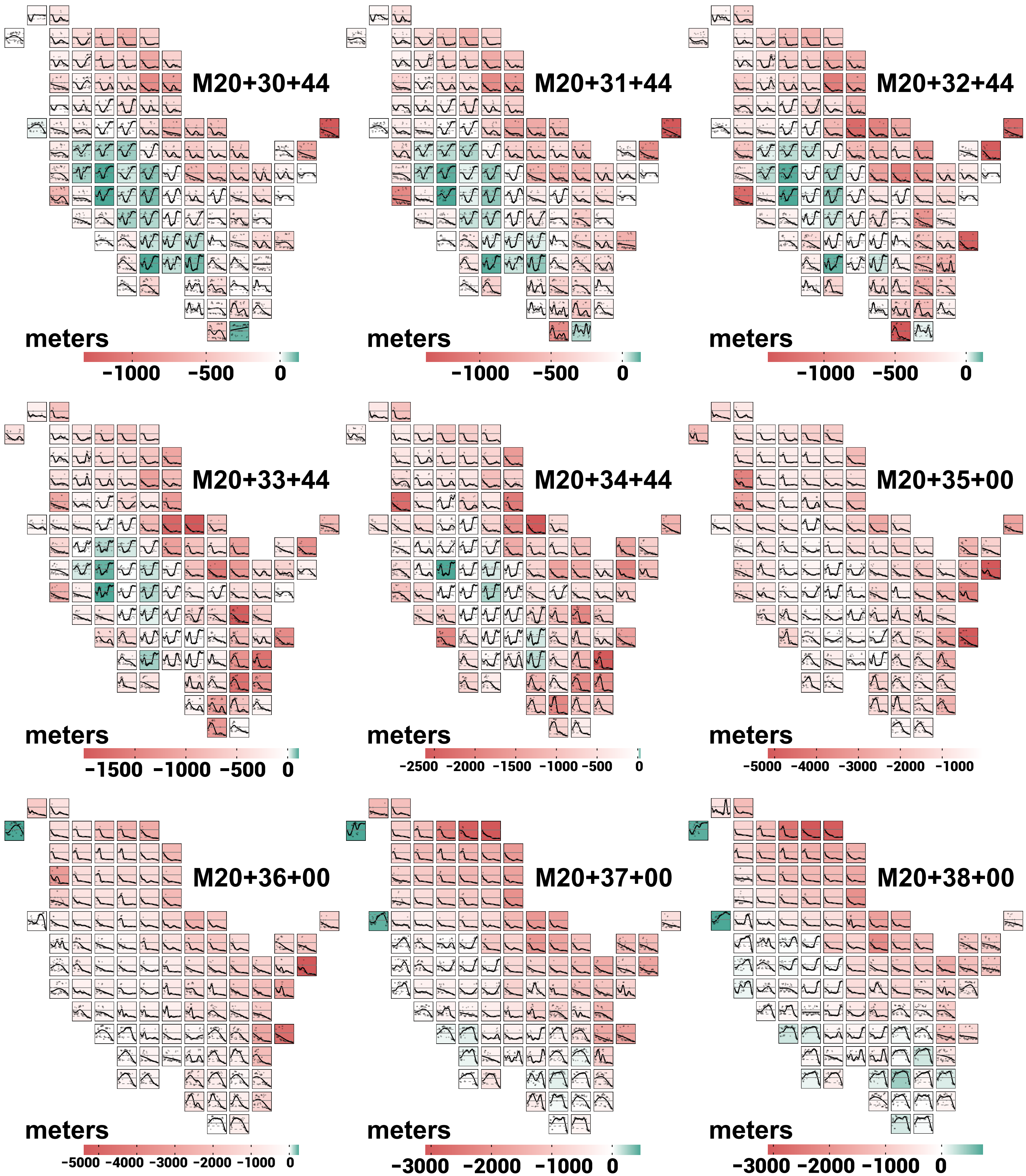

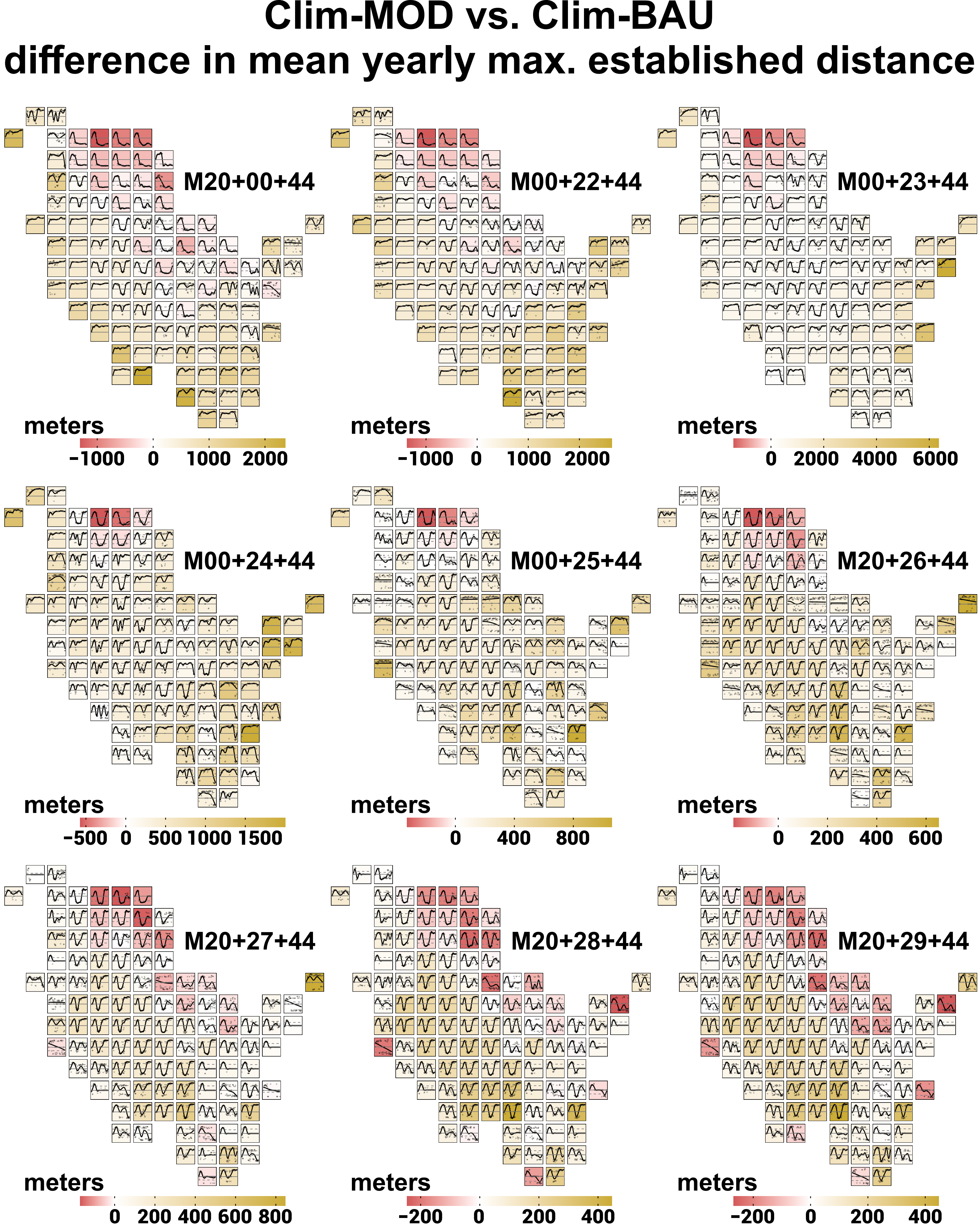

Clim-MOD vs. Clim-BAU
difference in mean yearly max. established distance
M20+00+44
meters
−1000 0 1000 2000
M00+22+44
meters
−1000 0 1000 2000
M00+23+44
meters
0 2000 4000 6000
M00+24+44
meters
−500 0 500 1000 1500
M00+25+44
meters
0 400 800
M20+26+44
meters
0 200 400 600
M20+27+44
meters
0 200 400 600 800
M20+28+44
meters
−200 0 200 400
M20+29+44
meters
−200 0 200 400

### Clim-MOD vs. Clim-BAU
### difference in mean yearly max. established distance

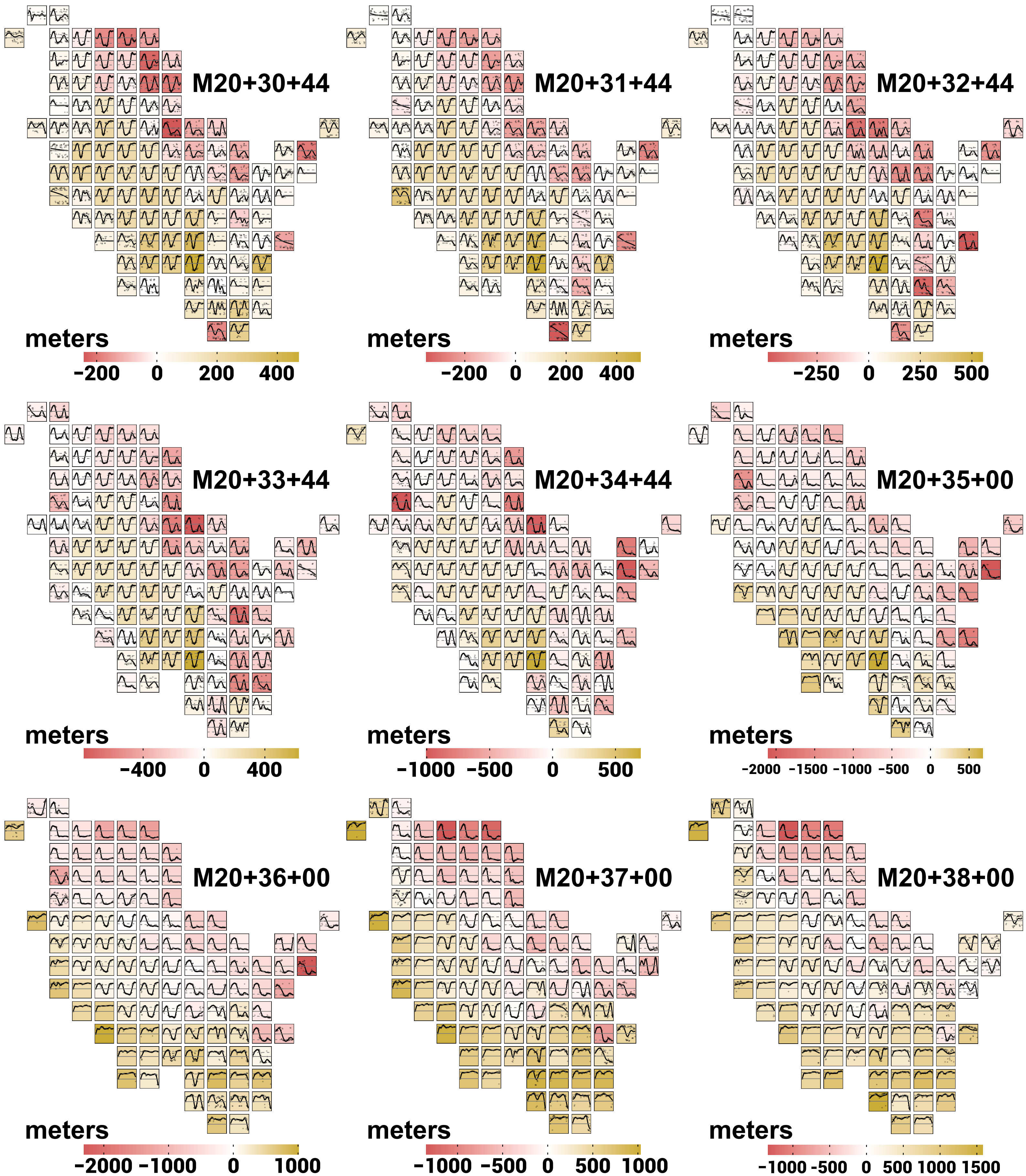

#### Section 2: difference in *population size*

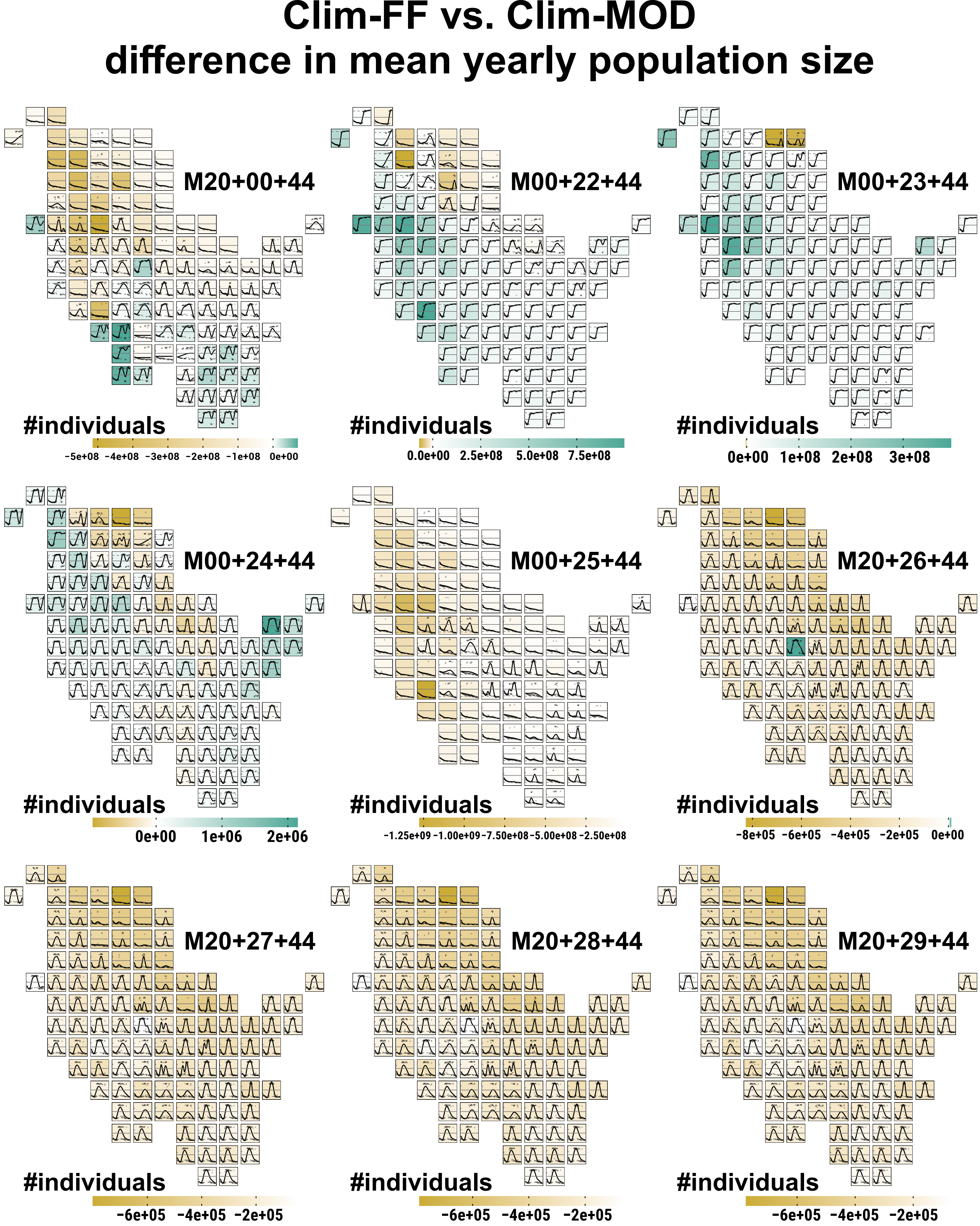
Clim-FF vs. Clim-MOD
difference in mean yearly population size
M20+00+44
#individuals
-5e+08 -4e+08 -3e+08 -2e+08 -1e+08 0e+00
M00+22+44
#individuals
0.0e+00 2.5e+08 5.0e+08 7.5e+08
M00+23+44
#individuals
0e+00 1e+08 2e+08 3e+08
M00+24+44
#individuals
0e+00 1e+06 2e+06
M00+25+44
#individuals
-1.25e+09 -1.00e+09 -7.50e+08 -5.00e+08 -2.50e+08
M20+26+44
#individuals
-8e+05 -6e+05 -4e+05 -2e+05 0e+00
M20+27+44
#individuals
-6e+05 -4e+05 -2e+05
M20+28+44
#individuals
-6e+05 -4e+05 -2e+05
M20+29+44
#individuals
-6e+05 -4e+05 -2e+05

### Clim-FF vs. Clim-MOD difference in mean yearly population size

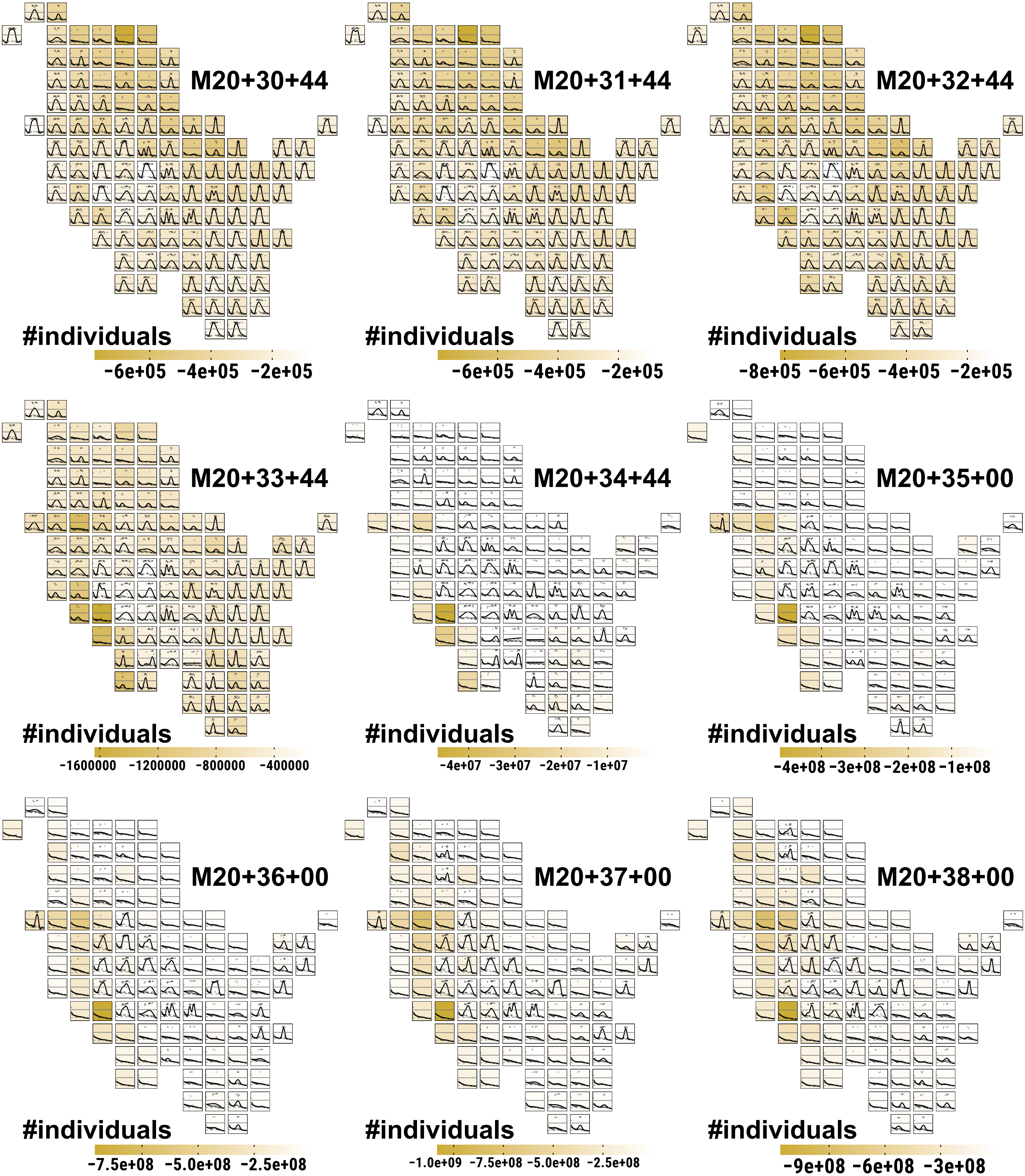

Clim-FF vs. Clim-BAU
difference in mean yearly population size

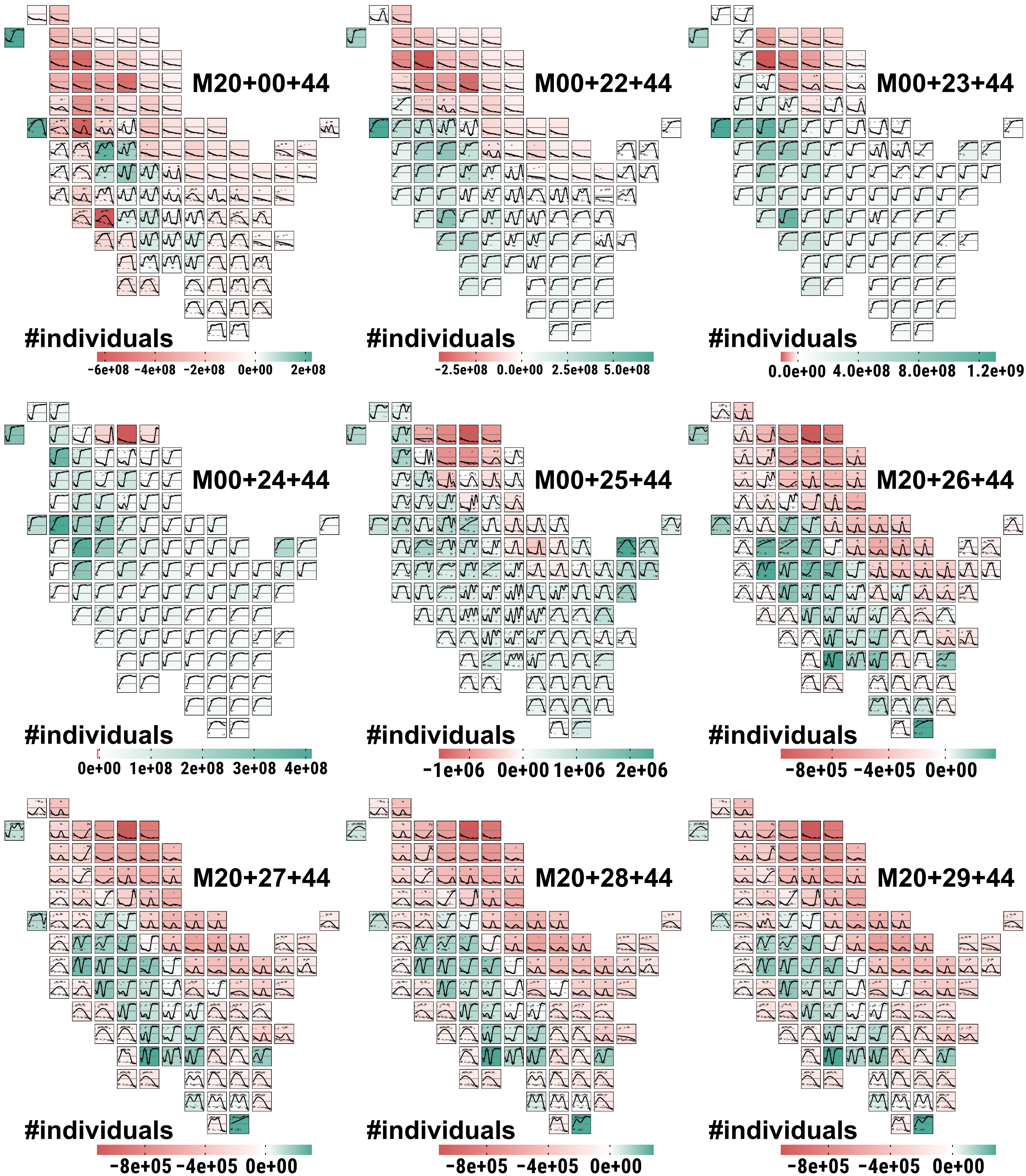

### Clim-FF vs. Clim-BAU
### difference in mean yearly population size

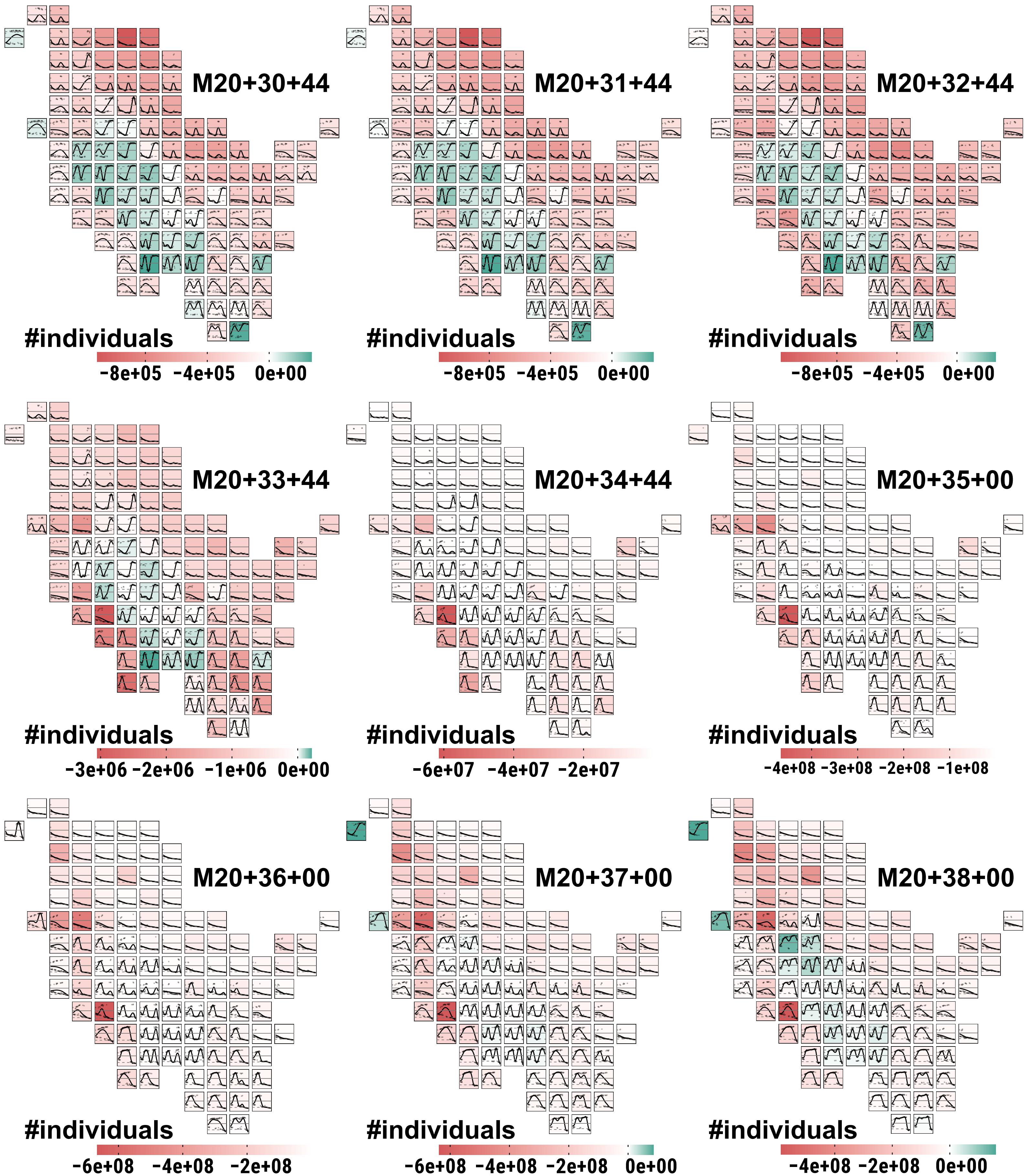

### Clim-MOD vs. Clim-BAU
### difference in mean yearly population size

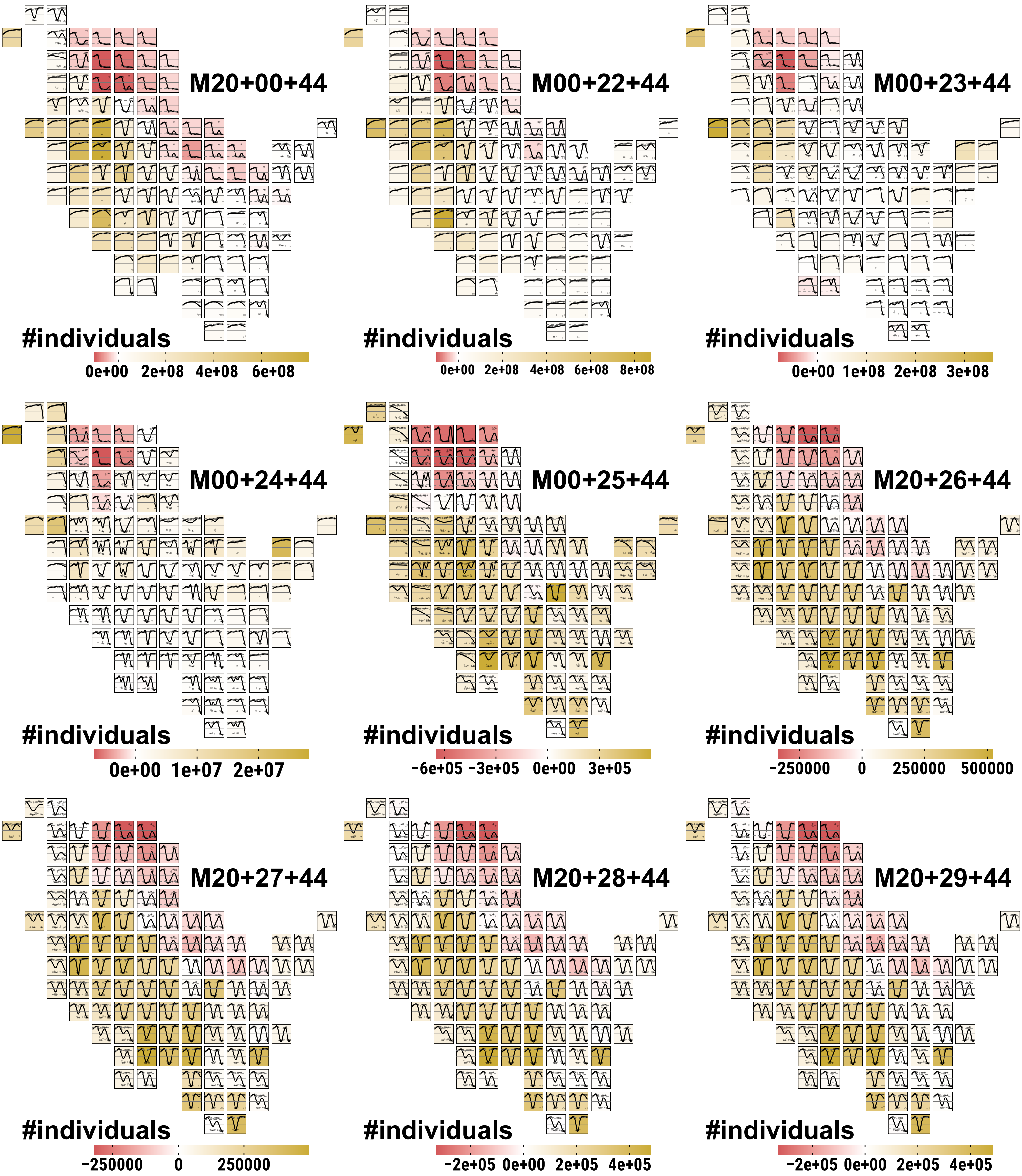

### Clim-MOD vs. Clim-BAU
### difference in mean yearly population size

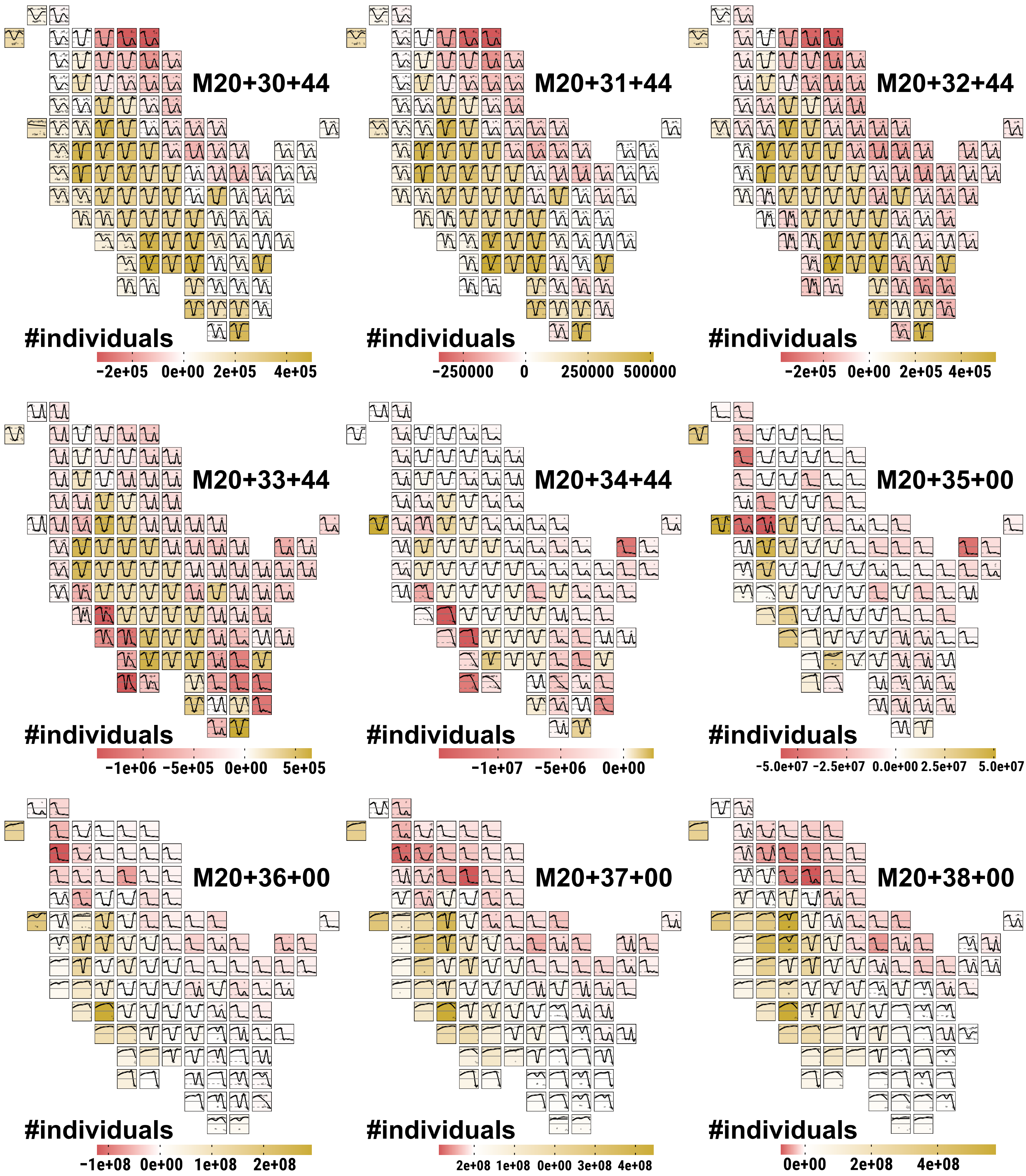

#### Section 3: difference in *population density*

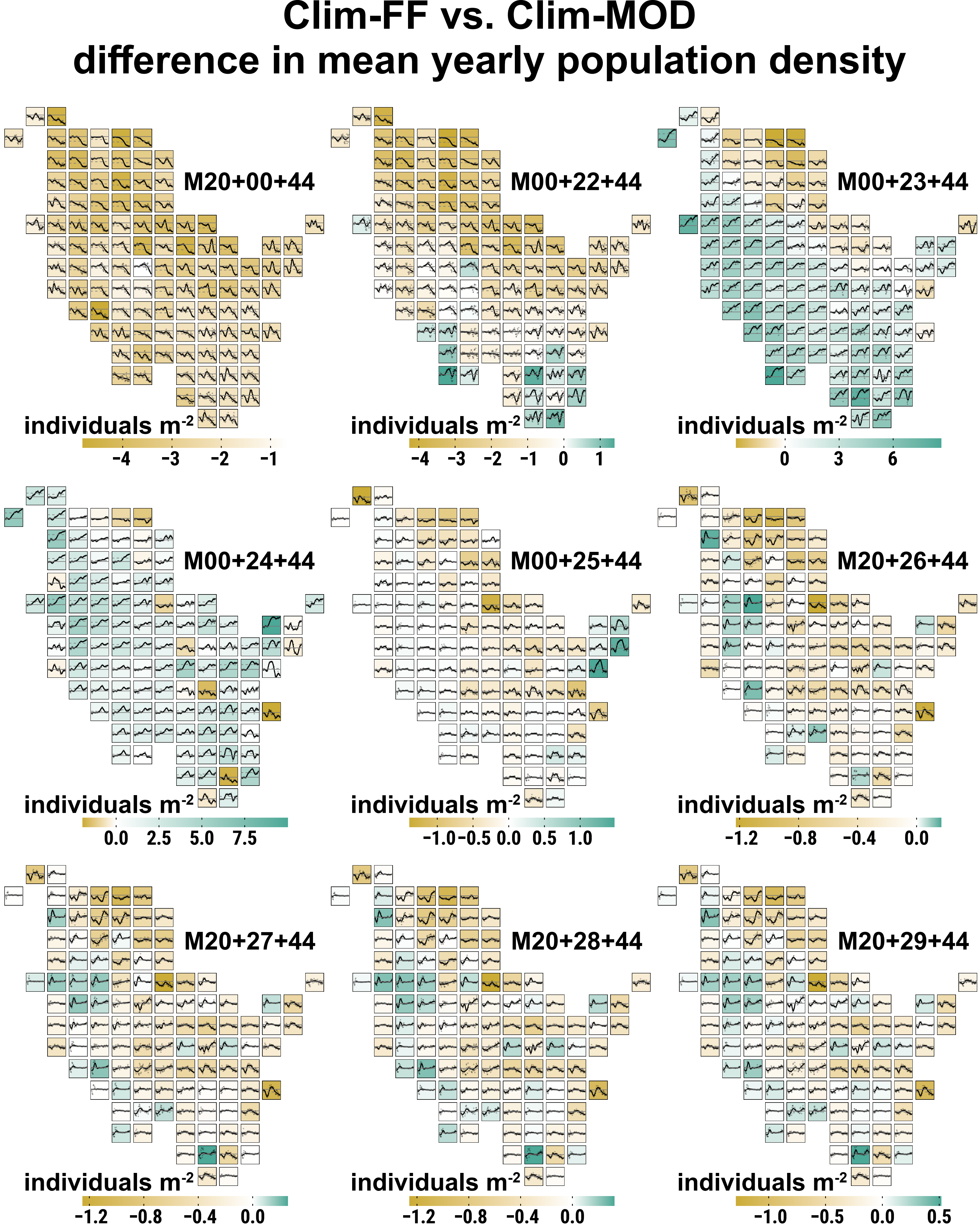
Clim-FF vs. Clim-MOD
difference in mean yearly population density
M20+00+44
individuals m-2
-4 -3 -2 -1
M00+22+44
individuals m-2
-4 -3 -2 -1 0 1
M00+23+44
individuals m-2
0 3 6
M00+24+44
individuals m-2
0.0 2.5 5.0 7.5
M00+25+44
individuals m-2
-1.0 -0.5 0.0 0.5 1.0
M20+26+44
individuals m-2
-1.2 -0.8 -0.4 0.0
M20+27+44
individuals m-2
-1.2 -0.8 -0.4 0.0
M20+28+44
individuals m-2
-1.2 -0.8 -0.4 0.0
M20+29+44
individuals m-2
-1.0 -0.5 0.0 0.5

### Clim-FF vs. Clim-MOD
### difference in mean yearly population density

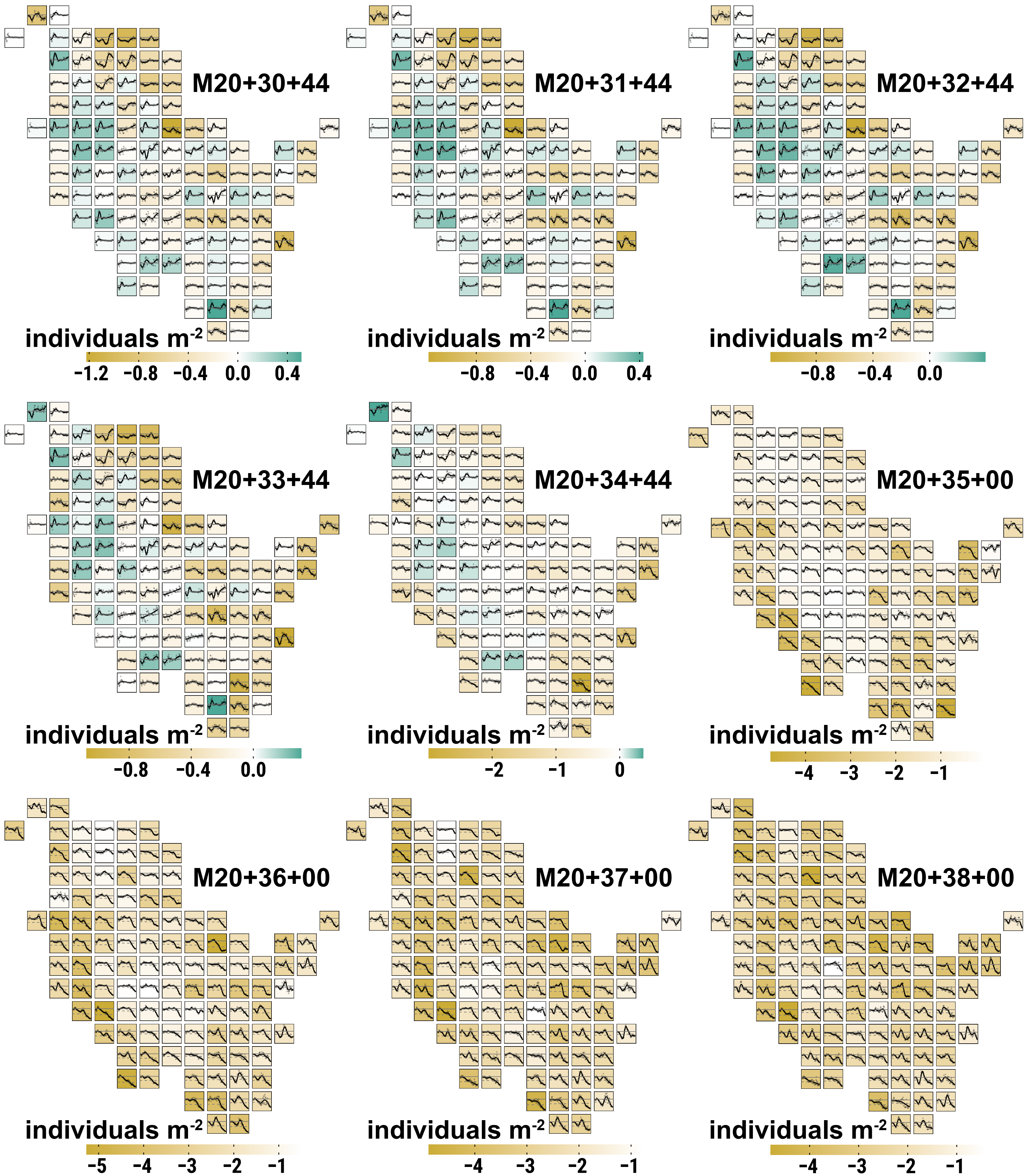

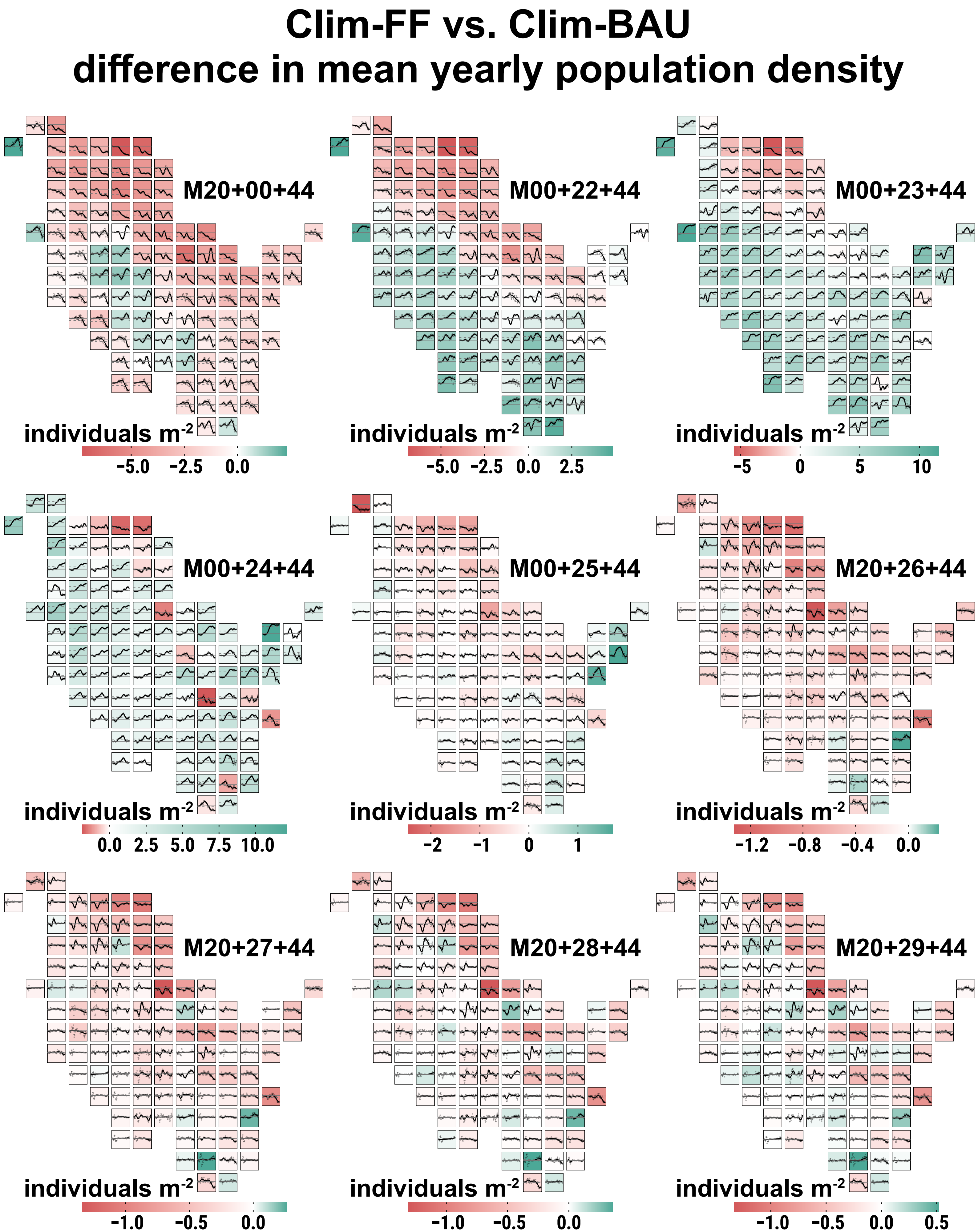

Clim-FF vs. Clim-BAU
difference in mean yearly population density
M20+00+44
individuals m-2
−5.0 −2.5 0.0
M00+22+44
individuals m-2
−5.0 −2.5 0.0 2.5
M00+23+44
individuals m-2
−5 0 5 10
M00+24+44
individuals m-2
0.0 2.5 5.0 7.5 10.0
M00+25+44
individuals m-2
−2 −1 0 1
M20+26+44
individuals m-2
−1.2 −0.8 −0.4 0.0
M20+27+44
individuals m-2
−1.0 −0.5 0.0
M20+28+44
individuals m-2
−1.0 −0.5 0.0
M20+29+44
individuals m-2
−1.0 −0.5 0.0 0.5

### Clim-FF vs. Clim-BAU
### difference in mean yearly population density

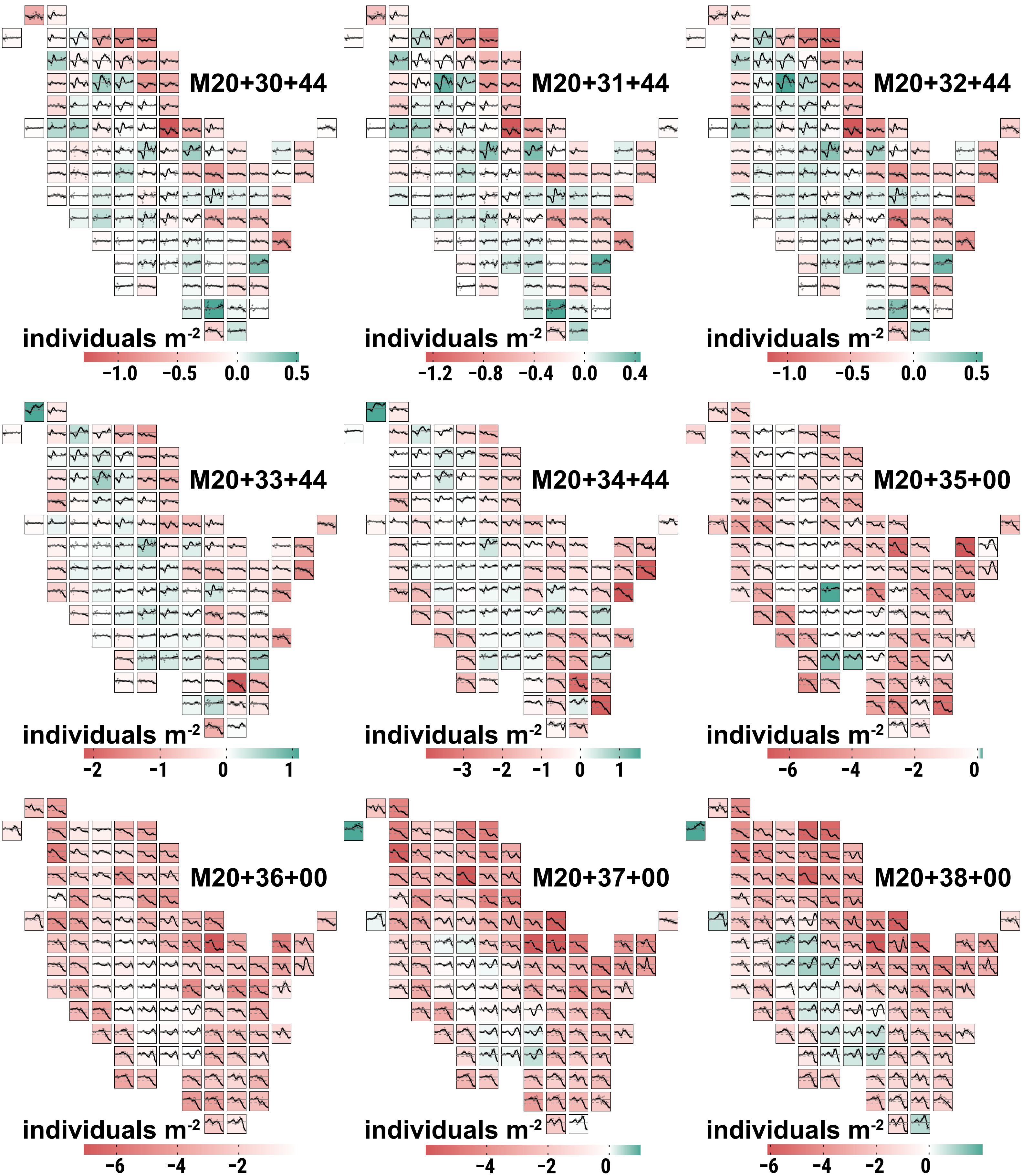

### Clim-MOD vs. Clim-BAU
### difference in mean yearly population density

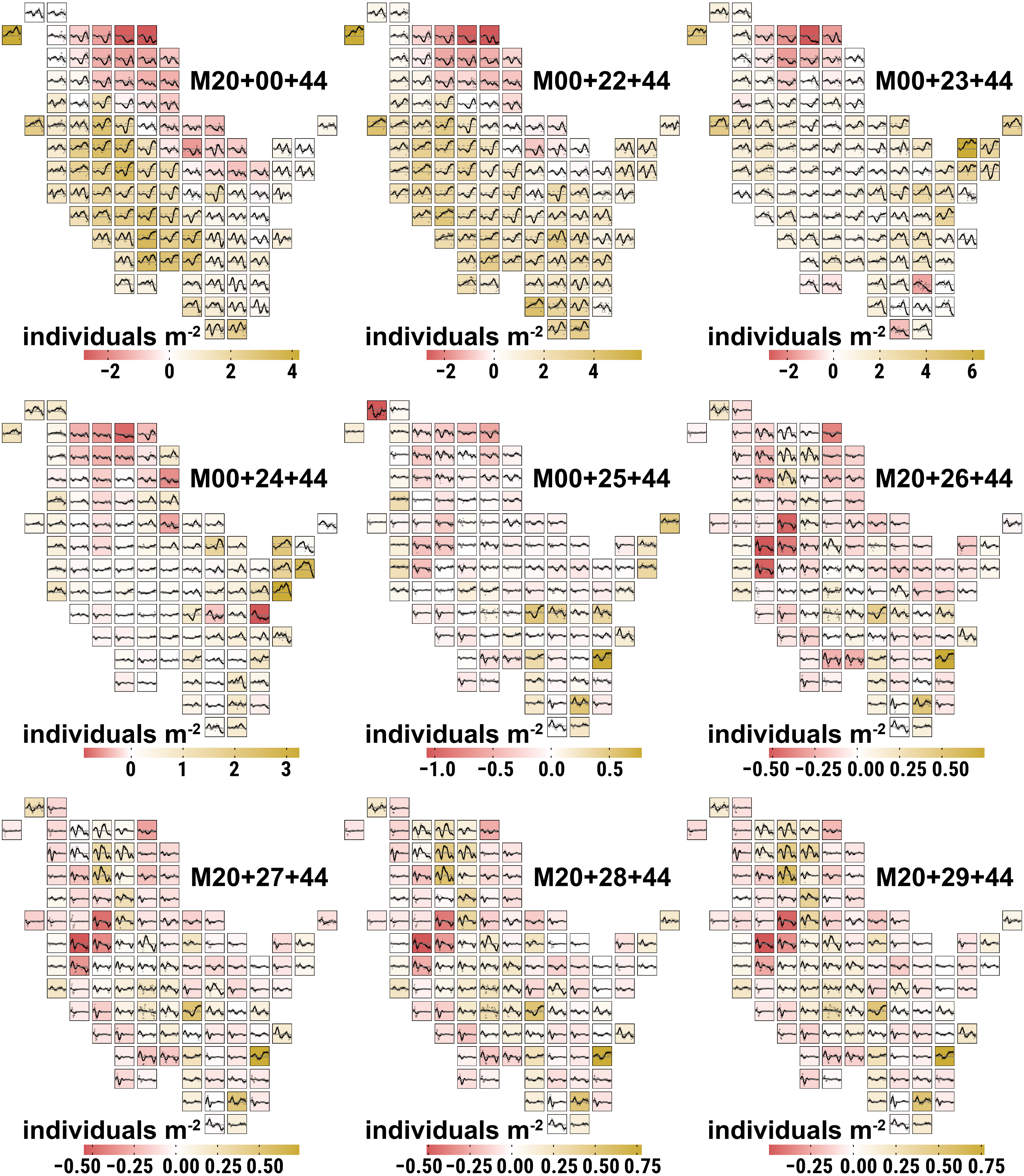

### Clim-MOD vs. Clim-BAU
### difference in mean yearly population density

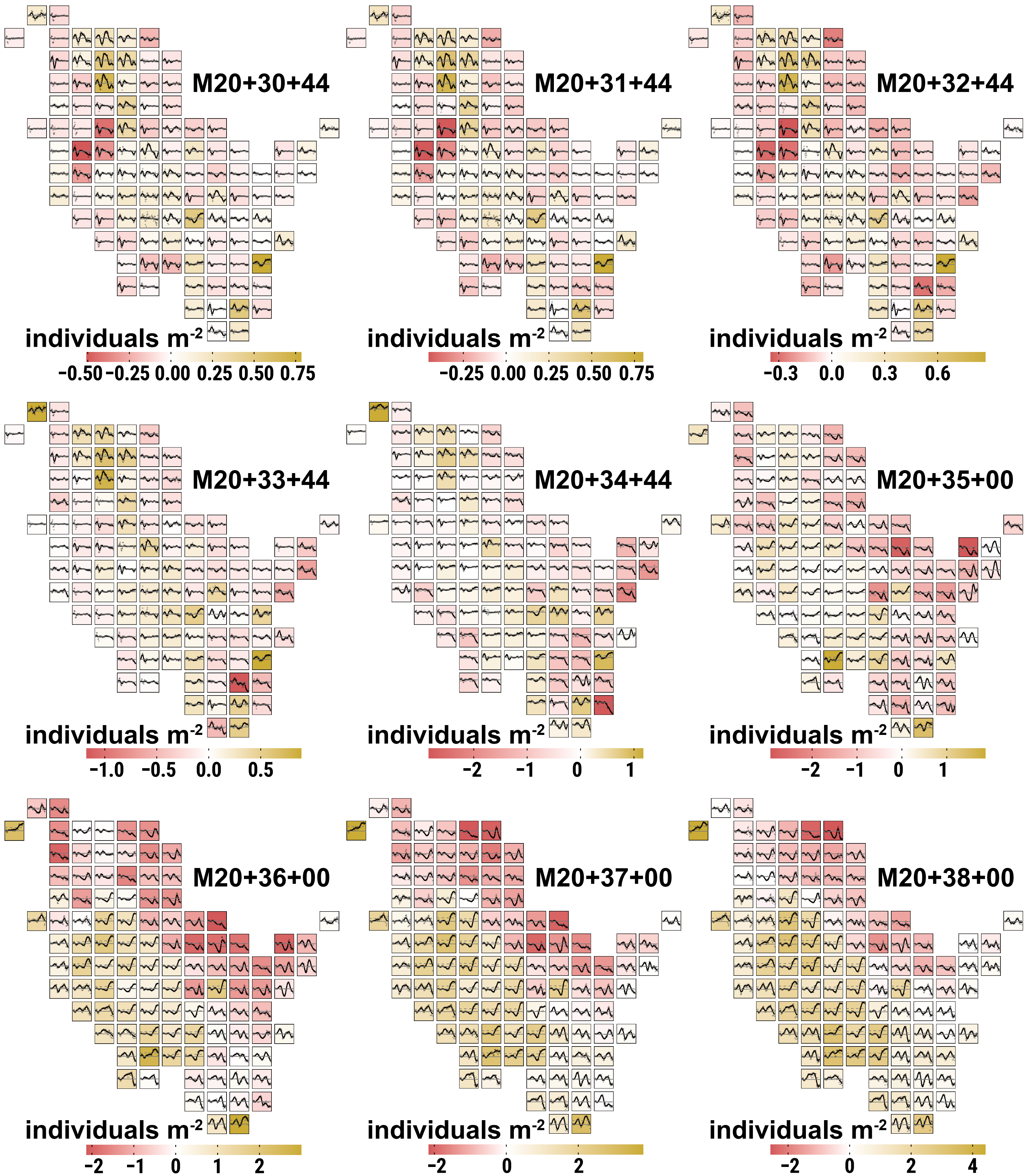
