## Supplementary material for "Large scale PVA modelling of insects in cultivated grasslands: the role of dispersal in mitigating the effects of management schedules under climate change": Specs of climate and grassland cells

This document describes the data in CSV files found at [ufz.de/record/dmp/archive/11900](ufz.de/record/dmp/archive/11900)

File *Supplement_S4-1_climate_cells.csv*:
Specifications per climate cell as defined in Section 2 of the main manuscript. The first column assigns a unique identifier (ID) to each cell (cf. Figure 2A, main manuscript). Columns 2-7 adapt specifications from the applied climate data (Section 2.3, main manuscript), where second and third column are the cell's Cartesian coordinates, fourth and fifth their longitude and latitude and sixth and seventh column their longitude and latitude in rotate pole grid coordinates. The last two columns represent a climate cell's geometric center in terms of the Cartesian coordinate system of the grassland cells (Supplement S4-2)

File *Supplement_S4-2_grassland_cells.csv*:
Specifications, mapping to climate cells and definition of weights applied for bilinear interpolation per grassland cell (Section 2.1, main manuscript). First and second column give the grassland cell's Cartesian coordinate. The remaining columns are subdivided into four three column blocks representing the four potential neighboring climate cells used for bilinear interpolation. Columns labeled ID_x contain the climate cell ID (Supplement S4-1) of a respective neighbor, columns labeled DIR_x the cardinal direction of this neighbor and columns labeled w_x the bilinear weight of this neighbor. The "x" in the previous description is replaced by the numbers 1-4, where the higher numbers represent the closer climate cells. The ID found in column ID_1 is the climate cell the grassland cell geographically belongs to. If ID_2-4, DIR_2-4 and w_2-4 are empty or contain a zero, no respective neighboring climate cell exists.
