## Supplementary material for "Large scale PVA modelling of insects in cultivated grasslands: the role of dispersal in mitigating the effects of management schedules under climate change": Illustration of correlation between evaluation parameters

The scatter plots below correlate three evaluation parameters (rows) from simulations with dispersal (population size [Σ eggs], population density [eggs $m^{-2}$], maximum established distance [meters]) with population density [eggs $m^{-2}$] stemming from simulations without dispersal as well as regional grassland cover [in %] in a radius ≤ 1,500 m of a simulation run's respective habitat of origin (columns). Each colored dot represents the correlated values of INDEPENDENT simulation runs, or rather their mean over 10 replicate runs (5 without dispersal), resulting from the population dynamics and dispersal process of a SINGLE initial population in the center of one out of 107 cells. The dot colors distinguish the three climate change scenarios (CCS) Clim-FF (RCP 2.6, green), Clim-MOD (RCP 4.5, brown) and Clim-BAU (RCP 8.5, pink). The colored numbers give the correlation coefficient ρ for the respective parameter combinations by CCS. The black lines represent the respective linear regression, where dashed=Clim-FF, dotted=Clim-MOD and dash dotted=Clim-BAU.

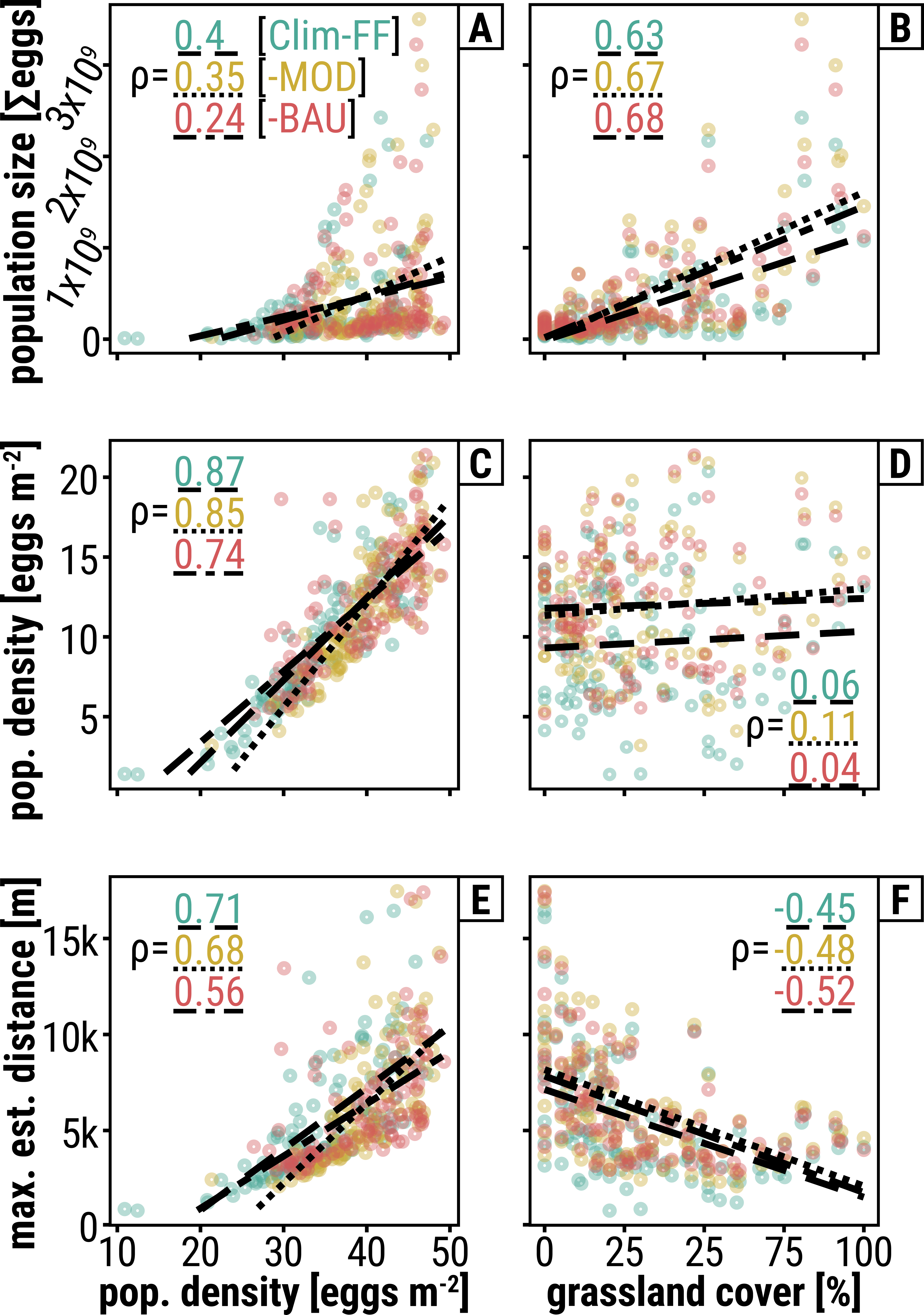

A
ρ= 0.4 [Clim-FF]
0.35 [-MOD]
0.24 [-BAU]
population size [Σeggs]
0
1x10⁹
2x10⁹
3x10⁹
B
ρ= 0.63
0.67
0.68
C
ρ= 0.87
0.85
0.74
pop. density [eggs m⁻²]
5
10
15
20
D
ρ= 0.06
0.11
0.04
E
ρ= 0.71
0.68
0.56
max. est. distance [m]
0
5k
10k
15k
F
ρ= -0.45
-0.48
-0.52
10
20
30
40
50
pop. density [eggs m⁻²]
0
25
25
75
100
grassland cover [%]
